# Hypermetabolism in mice carrying a near complete human chromosome 21

**DOI:** 10.1101/2023.01.30.526183

**Authors:** Dylan C. Sarver, Cheng Xu, Susana Rodriguez, Susan Aja, Andrew E. Jaffe, Feng J. Gao, Michael Delannoy, Muthu Periasamy, Yasuhiro Kazuki, Mitsuo Oshimura, Roger H. Reeves, G. William Wong

**Affiliations:** Department of Physiology, Johns Hopkins University School of Medicine, Baltimore, Maryland, USA; Center for Metabolism and Obesity Research, Johns Hopkins University School of Medicine, Baltimore, Maryland, USA; Department of Neuroscience, Johns Hopkins University School of Medicine, Baltimore, MD, USA; Department of Psychiatry and Behavioral Sciences, Johns Hopkins University School of Medicine, Baltimore, MD, USA; Department of Mental Health, Johns Hopkins Bloomberg School of Public Health, Baltimore, MD, USA; The Lieber Institute for Brain Development, Baltimore, MD, USA; Center for Computational Biology, Johns Hopkins University, Baltimore, MD, USA; Department of Genetic Medicine, Johns Hopkins University School of Medicine, Baltimore, MD, USA; Department of Biostatistics, Johns Hopkins Bloomberg School of Public Health, Baltimore, MD, USA; Department of Cell Biology, Johns Hopkins University School of Medicine, Baltimore, MD, USA; Department of Physiology and Cell Biology, The Ohio State University, Columbus, OH, USA; Burnett School of Biomedical Sciences, College of Medicine, University of Central Florida, Orlando, FL, USA; Division of Genome and Cellular Functions, Department of Molecular and Cellular Biology, School of Life Science, Faculty of Medicine, Tottori University, Tottori, Japan; Chromosome Engineering Research Center, Tottori University, Tottori, Japan

**Keywords:** Aneuploidy, trisomy, hypermetabolism, sarcolipin, SERCA pump, futile cycle

## Abstract

The consequences of aneuploidy have traditionally been studied in cell and animal models in which the extrachromosomal DNA is from the same species. Here, we explore a fundamental question concerning the impact of aneuploidy on systemic metabolism using a non-mosaic transchromosomic mouse model (TcMAC21) carrying a near complete human chromosome 21. Independent of diets and housing temperatures, TcMAC21 mice consume more calories, are hyperactive and hypermetabolic, remain consistently lean and profoundly insulin sensitive, and have a higher body temperature. The hypermetabolism and elevated thermogenesis are due to sarcolipin overexpression in the skeletal muscle, resulting in futile sarco(endo)plasmic reticulum Ca^2+^ ATPase (SERCA) activity and energy dissipation. Mitochondrial respiration is also markedly increased in skeletal muscle to meet the high ATP demand created by the futile cycle. This serendipitous discovery provides proof-of-concept that sarcolipin-mediated thermogenesis via uncoupling of the SERCA pump can be harnessed to promote energy expenditure and metabolic health.

## INTRODUCTION

The presence of an extra chromosome in mammals is generally lethal during fetal development, due to widespread cellular havoc caused by misregulated gene expression arising from gene dosage imbalance (1). Down syndrome (DS), resulting from trisomy of chromosome 21, is one of the rare aneuploidies compatible with life although as many as 80% of trisomy 21 conceptuses miscarry (2). The increased expression of ∼500 transcribed sequences of human chromosome 21 (Hsa21) affects many cell types and organ systems during development and in the postnatal period (3, 4). Humans with trisomy 21 have cognitive deficits, altered craniofacial development, and are at significantly higher risk for congenital heart defects, hearing and vision loss, leukemia, gastrointestinal disease, and early-onset dementia (2).

Given the significant impact of intellectual disability on the lives of individuals with DS, research emphasis has naturally focused on the neurological deficits underpinning trisomy 21 (5). In addition to developmental abnormalities associated with DS, there is an increasing awareness that adolescents and adults with DS also have an increased incidence of obesity, insulin resistance, and diabetes (6–8). Although this was first noted in the 1960s (9), the underlying cause for these metabolic dysregulations is mostly unknown and largely underexplored. Beyond clinical observations, limited studies have been conducted to determine the physiological underpinnings of metabolic impairments seen in DS [reviewed in (10)]. Our recent study on the Ts65Dn mouse model represents the most in-depth metabolic analysis, to date, of any Down syndrome mouse model (11). However, the segmental trisomic Ts65Dn mouse contains only ∼55% of the orthologous protein-coding genes found on Hsa21 (12). In addition, it contains additional trisomic genes from the centromeric region of mouse chromosome 17 (Mmu17) not found in Hsa21, thus complicating the genotype-phenotype relationships (13, 14).

In the past two decades, more than 20 mouse models of DS have been generated (15). Despite their utility in advancing DS research, none of these models recapitulate the full spectrum of human DS. With the exception of Tc1, all the DS mouse models are trisomic for some, but not all, of the orthologous mouse genes found in Hsa21 (15). Tc1 is the first mouse model with an independently segregating Hsa21 (16). However, Tc1 mice are missing >50 of the 220 protein coding genes on Hsa21 due to deletion and mutations (17). In addition, Tc1 mice show extensive mosaicism (i.e., the human chromosome is present in zygotes but is lost randomly from cells during development). As a consequence, every mouse has a unique developmental trajectory, complicating the interpretations of results obtained from Tc1 mice. To overcome the limitations of previously generated trisomic mouse models, a transchromosomic mouse model (TcMAC21) carrying an independently segregating and a near complete copy of Hsa21 was recently developed (18). TcMAC21 is not mosaic and contains 93% of HSA21q protein-coding genes (PCGs), and is considered the most representative mouse model of DS.

Both mouse and rat that carry a non-mosaic Hsa21 recapitulate many DS phenotypes related to the central nervous system (e.g., reduced cerebellum volume, learning and memory deficit), craniofacial skeleton, and heart (18, 19). The metabolic phenotype of TcMAC21, however, is unknown and has yet to be examined. The availability of the TcMAC21 mouse model has afforded a unique opportunity to address two fundamental questions: 1) what is the impact of aneuploidy on systemic metabolism; 2) what are the molecular, cellular, and physiological consequences of introducing a foreign (human) chromosome from an evolutionarily distant species into mice?

Unexpectedly, we discovered that TcMAC21 mice have all the hallmarks of hypermetabolism, driven by elevated mitochondrial respiration and futile sarco(endo)plasmic reticulum Ca^2+^ ATPase (SERCA) pump activity in the skeletal muscle as a consequence of endogenous sarcolipin (SLN) overexpression. Our study has provided further evidence and proof-of-concept that endogenous SLN-mediated uncoupling of the SERCA pump can be harnessed for energy dissipation, weight loss, and metabolic health.

## RESULTS

### Human chromosome 21 genes are differentially expressed and regulated in mouse adipose tissue, liver, and skeletal muscle

TcMAC21 mice carry a non-mosaic and independently segregating mouse artificial chromosome with a near complete copy of the long arm of human chromosome 21 (Hsa21q) (18). The Hsa21q in TcMAC21is comprised of ∼37 Mb and 199 protein coding genes (Fig. 1A). RNA-sequencing showed that TcMAC21 mice are capable of expressing Hsa21-derived transcripts in each of the tissues examined, and that gene expression is regulated in a tissue-specific manner. The transcriptional activity map of Hsa21 shows regions of gene expression and repression (Fig. 1B). There are three large regions of Hsa21 with little or no transcription activity: 29.5-31.1 Mb, 44.5-44.8 Mb, and 45.3-46.2 Mb. The first gap of transcriptional inactivity contains the protein-coding genes *Cldn8* and *Cldn17*, *Girk1*, and 33 distinct *Krtap* (keratin-associated protein) genes. The second transcriptionally inactive area contains 16 *Krtap* genes, and the third transcriptionally inactive area contains 8 protein-coding genes (*Col6a1, Col6a2, Col18a1, Fctd, Lss, Pcbp3, Slc19a1,* and *Spatc1l*). By filtering the transcriptional map to display expressed-genes only, we highlighted all the genes with their differential expression profiles across five major metabolic tissues— brown adipose tissue (BAT), inguinal white adipose tissue (iWAT), gonadal white adipose tissue (gWAT), liver, and skeletal muscle (Fig. 1C).

**Figure 1.**
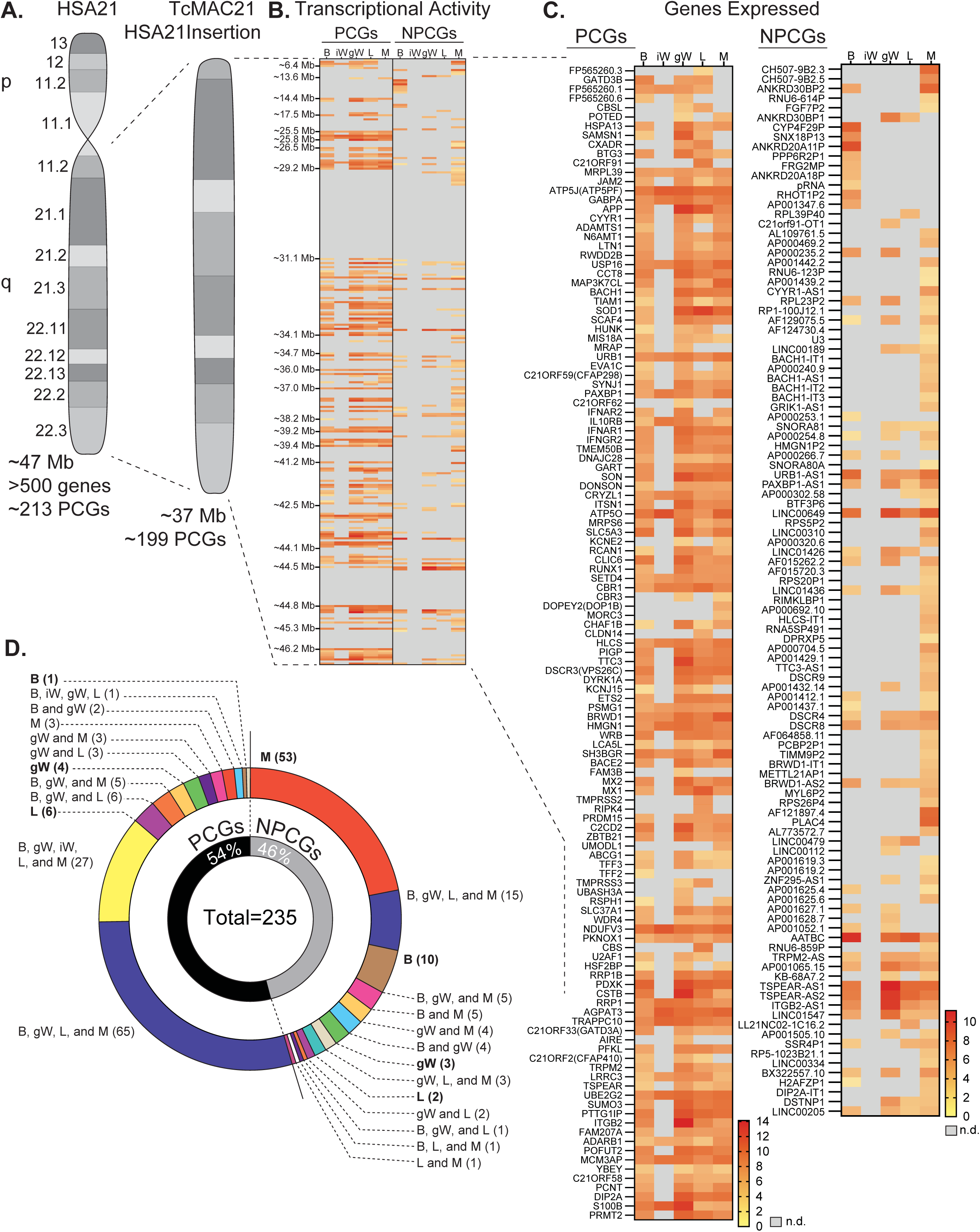
Human chromosome 21 genes are differentially expressed and regulated in mouse adipose tissue, liver, and skeletal muscle. **A)** Graphical representation of human chromosome 21 (Hsa21) and the entire long arm (Hsa21q) region carried by a mouse artificial chromosome in the transchromosomic mouse model (TcMAC21). Four deletions that occurred during generation of the transchromosomic mice eliminate 14/213 protein coding genes (PCGs; 7%) and 105/487 predicted or known non-protein coding genes (NPCGs; 22%) (18). **B)** Global view of transcriptionally expressed and repressed protein and non-protein coding gene regions over the entire Hsa21q across five tissues. Gray box denotes transcript that is not detected. **C)** Transcriptional activity map showing only Hsa21 genes expressed by at least one tissue. Gray box denotes transcript that is not detected. **D)** Overlap analysis showing shared expression of human protein coding and non-protein coding genes across five tissues. Of the 235 unique human genes expressed by the MAC21, 54% are protein coding and 46% are non-protein coding genes. PCGs, protein coding genes; NPCGs, non-protein coding genes; B, brown adipose tissue; iW, inguinal white adipose tissue; gW, gonadal white adipose tissue; L, liver; M, skeletal muscle (gastrocnemius); n.d., not detected. *n* = 5 RNA samples per group per tissue-type. Mice were on high-fat diet for 16 weeks at the time of tissue collection.

One of the more striking differences in expression profile is between visceral (gonadal) and subcutaneous (inguinal) white adipose tissue (gWAT and iWAT respectively). gWAT expresses 115 human protein-coding genes (PCGs) while iWAT expresses only 27. A similar pattern can be seen in the non-protein-coding genes (NPCGs). We were unable to detect any Hsa21-derived non-protein-coding genes in the iWAT, while in gWAT we observed 37. Overlap analysis was carried out to assess how similar expression profiles were between tissues (Fig. 1D). Of the 126 Hsa21-derived protein-coding genes expressed by at least one tissue, a majority (65 total) are shared between BAT, gWAT, liver, and skeletal muscle. Of the 109 Hsa21-derived non-protein-coding genes expressed by at least one tissue, the majority (53 total) are uniquely expressed by skeletal muscle. Of note, the liver uniquely expresses 6 PCGs and 2 NPCGs, gWAT 4 and 3, skeletal muscle 3 and 53, BAT 1 and 10, and iWAT 0 and 0. Together, these data indicate that Hsa21-derived transcripts are differentially expressed and regulated across major metabolic tissues in TcMAC21 mice.

### Hypermetabolism in TcMAC21 mice

Having established that Hsa21-derived transcripts are differentially expressed and regulated in mouse organs and tissues, we next asked the impact of the extra human genetic material and genes on systemic metabolism. As previously documented, TcMAC21 pups are born at the same weight as their euploid littermates (18). However, by 3 months of age TcMAC21 mice fed a standard chow weighed significantly less (∼8.5 g) than euploid littermates, and this weight difference remained stable over time (Fig. 2A). The size and body weight differences were not due to reduced plasma IGF-1 and growth hormone, as their circulating levels were in fact higher in TcMAC21 compared to euploid mice (Fig. S1). Body composition analysis showed that TcMAC21 have significantly reduced absolute and relative (normalized to body weight) fat mass (Fig. 2B). Although the absolute lean mass was reduced in TcMAC21 mice, the relative lean mass (normalized to body weight) was not different between genotypes. Tissue collection at termination of the study also showed smaller visceral and subcutaneous fat mass and liver weight in TcMAC21 mice (Table S1).

**Figure 2.**
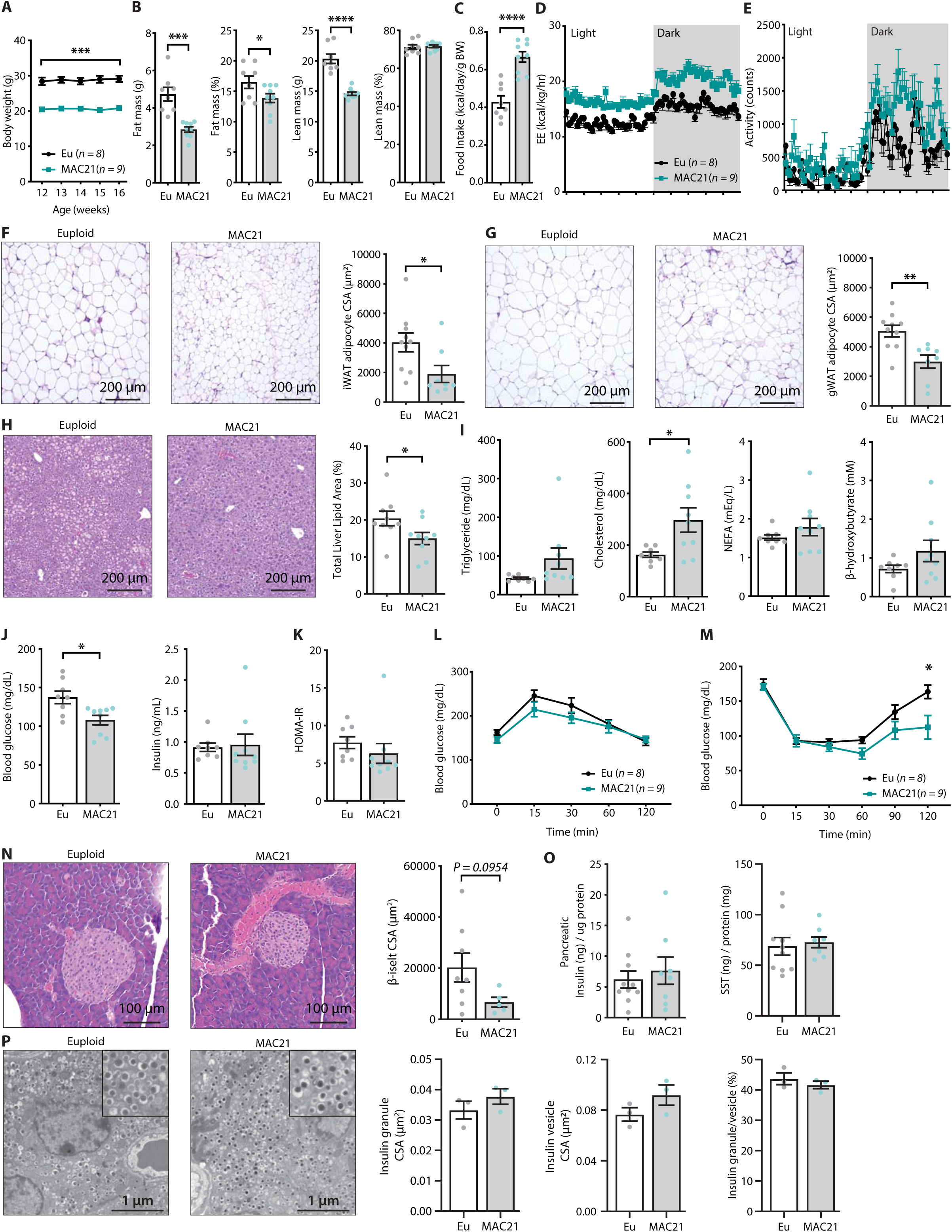
Hypermetabolism in TcMAC21 mice. **A)** Body weights of mice fed standard chow. **B)** Body composition analysis of fat and lean mass (relative to body weight). **C)** Food intake over a 24 h period. **D-E)** Energy expenditure (EE) and physical activity level over 24 hr period in chow-fed mice. **F)** Hematoxylin and eosin (H&E) stained sections of inguinal white adipose tissue (iWAT), and adipocyte cross-sectional area (CSA) quantification. **G)** Histology of gonadal white adipose tissue (gWAT), and adipocyte CSA quantification. **H)** Histology of liver tissues with quantification of area covered by lipid droplets per focal plane. **I)** Fasting serum triglyceride, cholesterol, non-esterified fatty acids (NEFA), β-hydroxybutyrate (ketone) levels. **J)** fasting blood glucose and insulin levels. **K)** Insulin resistance index (homeostatic model assessment for insulin resistance (HOMA-IR). **L)** Glucose tolerance tests. **M)** Insulin tolerance tests. **N)** Histology of pancreas and quantification of β-islet CSA. **O)** Pancreatic insulin and somatostatin (SST) contents (normalized to pancreatic protein input). **P)** Electron micrographs (EM) of pancreatic β-cells showing dense insulin granules and their surrounding vesicles, and the quantification of insulin granule CSA, insulin vesicle CSA, and the ratio of insulin granule to insulin vesicle. *n* = 8 euploid and 9 TcMAC21 mice for all graphs from A to Y. *n* = 8-10 euploid and 5-8 TcMAC21 samples used for pancreatic analysis by H&E and protein quantification, graphs N and P. *n* = 3 euploid and 3 TcMAC21 used for EM quantification; each data point represents 1,200 insulin granules and 1,200 insulin vesicles quantified across 6 unique locations per mouse, graphs P.

Differences in body weight were not due to reduced caloric intake, as TcMAC21 mice actually consumed a significantly higher amount of food relative to their body weight than euploid controls (Fig. 2C). Indirect calorimetry analysis indicated that TcMAC21 mice—regardless of the photocycle (light or dark phase) and metabolic states (*ad libitum* fed, fasted, refed)—were expending ∼25% more energy and were significantly more active compared to euploid controls (Fig. 2D-E and Fig. S2 and Table S2). Despite much higher caloric intake per gram body mass, TcMAC21 mice were much leaner due to substantially elevated physical activity and energy expenditure. Hyperactivity and elevated energy expenditure were not due to altered circulating thyroid hormones, as serum triiodothyronine (T_3_, the active form of TH) levels were not different between chow fed TcMAC21 and euploid mice (Fig. S1). Serum level of thyroxine (T_4_), the precursor of T_3_, were modestly elevated in TcMAC21 relative to euploid mice.

In accordance with the lean phenotype, TcMAC21 mice had significantly smaller adipocyte cell size in both subcutaneous (inguinal) and visceral (gonadal) fat depots (Fig. 2F-G), as well as significantly reduced fat accumulation in liver (Fig. 2H). Fasting triglyceride, non-esterified fatty acid (NEFA), and β-hydroxybutyrate levels were not different between genotypes; fasting cholesterol, however, was higher in TcMAC21 mice (Fig. 2I). Although fasting insulin levels were not different between groups, fasting blood glucose was significantly lower in TcMAC21 mice (Fig. 2J). The insulin resistance index (HOMA-IR), along with glucose and insulin tolerance tests suggested modest improvements in insulin sensitivity in TcMAC21 relative to euploid mice (Fig. 2K-M). Assessment of the pancreas showed that TcMAC21 mice have similar β-islet cross-sectional area (CSA), insulin and somatostatin content, and insulin granule and vesicle size compared to euploid controls (Fig. 2N-P). Taken together, these data indicate that chow-fed TcMAC21 mice at baseline are lean despite increased caloric intake, and this is largely due to elevated physical activity and energy expenditure.

### MAC21 mice are resistant to diet-induced obesity and metabolic dysfunction

The hypermetabolic phenotypes seen in chow-fed TcMAC21 predicted that these mice would be resistant to diet-induced obesity and metabolic dysfunction. Indeed, after 8 weeks on high-fat diet (HFD), TcMAC21 mice gained only ∼3 g of body weight, whereas the euploid controls gained >15 g of body weight over the same period. Consequently, TcMAC21 mice weighed ∼50% less than euploid controls (Fig. 3A). Consistent with the lean phenotype, the absolute and relative (normalized to body weight) fat mass were markedly reduced compared to euploid controls (Fig. 3B). The weights of other organs (liver, kidney, BAT) at time of termination were also lower in TcMAC21 mice, but tibia length was not different between genotypes (Fig. S3 and Table S3). Complete blood count revealed no differences in erythroid, lymphoid, and myeloid cell numbers between genotypes (Table S4). Because relative lean mass was higher in TcMAC21 compared to euploid mice (Fig. 3B), the lean phenotype seen in HFD-fed TcMAC21 is largely due to reduced adiposity. Accordingly, TcMAC21 had significantly smaller adipocyte cell size in both subcutaneous (inguinal) and visceral (gonadal) fat depots, and a marked reduction in lipid accumulation in the liver (Fig. 3C-E).

**Figure 3.**
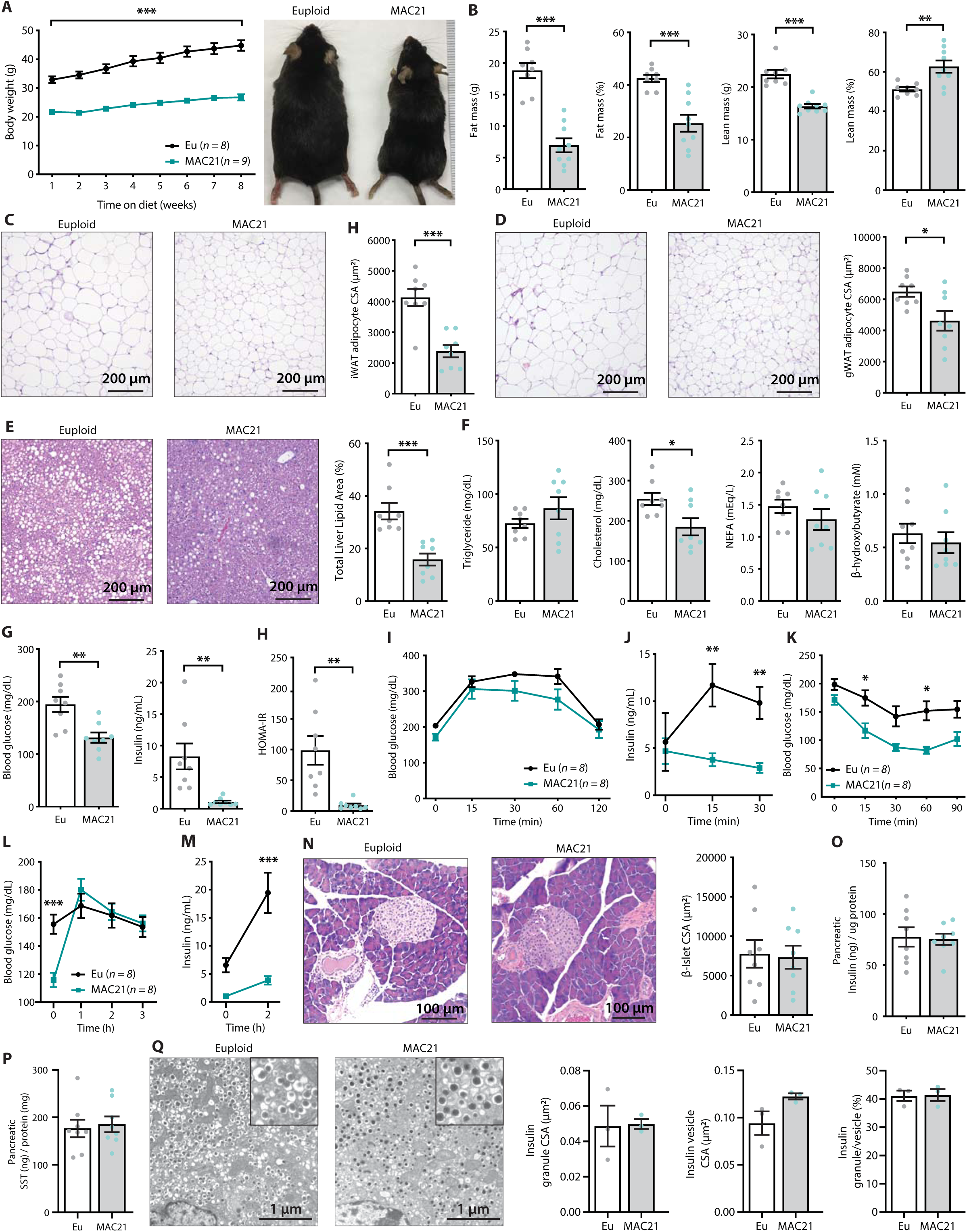
TcMAC21 mice are resistant to diet-induced obesity and metabolic dysfunction. **A)** Body weights over time on a high-fat diet and representative mouse images. **B)** Body composition analysis of fat and lean mass. **C)** Histology of inguinal white adipose tissue (iWAT), and quantification of adipocyte cross-sectional area (CSA). **D)** Histology of gonadal white adipose tissue (gWAT), and quantification of adipocyte CSA. **E)** Histology of liver tissues, and quantification of area covered by lipid droplets per focal plane. **F)** Fasting serum triglyceride, cholesterol, non-esterified fatty acids (NEFA), β-hydroxybutyrate (ketone) levels. **G)** Fasting blood glucose and insulin levels. **H)** Insulin resistance index (homeostatic model assessment for insulin resistance (HOMA-IR). **I)** Glucose tolerance tests (GTT). **J)** Serum insulin levels during GTT**. K)** Insulin tolerance tests. **L)** Blood glucose levels after an overnight (16 h) fast and 1, 2, and 3 h of food reintroduction. **M)** Serum insulin levels after a 16 h fast and 2 h of refeeding. **N)** Pancreas histology and quantification of β-islet CSA. **O-P)** Pancreatic insulin and somatostatin (SST) contents (normalized to pancreatic protein input). **Q)** Electron micrographs (EM) of pancreatic β-cells showing dense insulin granules and their surrounding vesicles, and quantification of insulin granule CSA, insulin vesicle CSA, and the ratio of insulin granule to insulin vesicle. *n* = 8 euploid and 8-9 TcMAC21 mice for all graphs from A to P. *n* = 3 euploid and 3 TcMAC21 used for EM quantification; each data point represents 1,200 insulin granules and 1,200 insulin vesicles quantified across 6 unique locations per mouse, graphs Q.

Although fasting serum triglyceride, NEFA, and β-hydroxybutyrate levels were not different between genotypes, serum cholesterol was significantly lower in TcMAC21 mice (Fig. 3F). Fasting glucose and insulin levels, and the insulin resistance index (HOMA-IR), were markedly lower in TcMAC21 mice relative to euploid controls (Fig. 3G-H), indicative of enhanced insulin sensitivity. In glucose tolerance tests (GTT), even though the rate of glucose disposal was similar between TcMAC21 and euploid mice, the amount of serum insulin present during GTT (time 0, 15, and 30 min) was dramatically lower in TcMAC21 (Fig. 3I-J). This indicates that a substantially lower amount of insulin is sufficient to promote glucose clearance in TcMAC21 at a rate comparable to euploid mice, consistent with elevated insulin sensitivity in the peripheral tissues. Indeed, when we directly assessed insulin action via insulin tolerance tests (ITT), TcMAC21 mice clearly exhibited higher insulin sensitivity as indicated by the significant differences in insulin-stimulated glucose disposal (Fig. 3K).

To independently confirm TcMAC21 mice are more insulin sensitive, we fasted the mice overnight (16 hr) then reintroduced them to food. Under this fasting-refeeding condition, we could clearly see the resumption of food intake was successful at increasing blood glucose in TcMAC21 (Fig. 3L); however, the insulin response to food intake in TcMAC21 mice was strikingly smaller in magnitude compared to euploid controls (Fig. 3M). Again, these data indicate that TcMAC21 mice are significantly more insulin sensitive since a substantially lower insulin response during fasting-refeeding is sufficient for glucose clearance at a rate comparable to euploid mice.

These results prompted us to determine if there were developmental changes in the pancreas leading to reduced insulin secretion in response to glucose administration or food intake, independent of elevated insulin sensitivity in peripheral tissues. Quantification of β-islet size, pancreatic insulin and somatostatin content, as well as insulin granule and vesicle size did not reveal any intrinsic differences between TcMAC21 and euploid mice (Fig. 3N-Q), thus ruling out a developmental cause and arguing in favor of enhanced insulin action. Pancreatic acinar zymogen granule size was also not different between genotypes (Fig. S4), suggesting normal development of the exocrine pancreas. Taken together, these data indicate that TcMAC21 mice are remarkably resistant to HFD-induced obesity and insulin resistance.

### Hypermetabolism of TcMAC21 mice is uncoupled from changes in adipose and liver transcriptomes

Next, we sought to uncover the physiological mechanisms responsible for TcMAC21 resistance to weight gain and developing insulin resistance when fed a high-fat diet. First, we wanted to rule out whether there is a change in caloric intake. TcMAC21 mice actually consumed a significantly higher amount of food (relative to their body weight) compared to euploid controls (Fig. 4A and Table S5). To rule out any potential dysfunction of the gut that might adversely affect nutrient absorption, we collected, counted, weighed, and subjected fecal samples of each mouse to fecal bomb calorimetry. Neither fecal frequency, average fecal pellet weight, nor fecal energy content was different between TcMAC21 and euploid mice (Fig. 4B and Fig. S5). Fecal energy content trended lower in TcMAC21 (Fig. 4B), implying a modest increase in the efficiency of nutrient absorption. Since energy input (caloric intake per gram body weight) was significantly higher in TcMAC21 and outputs (left over fecal energy) were similar across genotypes, the data strongly support hypermetabolism being the cause of the lean phenotype seen in TcMAC21 mice. Indeed, when we measured energy expenditure (EE) and physical activity of both groups, we found that TcMAC21 mice have markedly higher EE and physical activity irrespective of circadian cycle and metabolic states (Fig. 4C-D and Fig. S6 and Table S5). The striking difference in EE was very similar to TcMAC21 fed a standard chow (Fig. 2D), but to an even greater extent when mice were fed a high-fat diet, presumably due to the greater availability of calorie-dense lipid substrates for oxidation.

**Figure 4.**
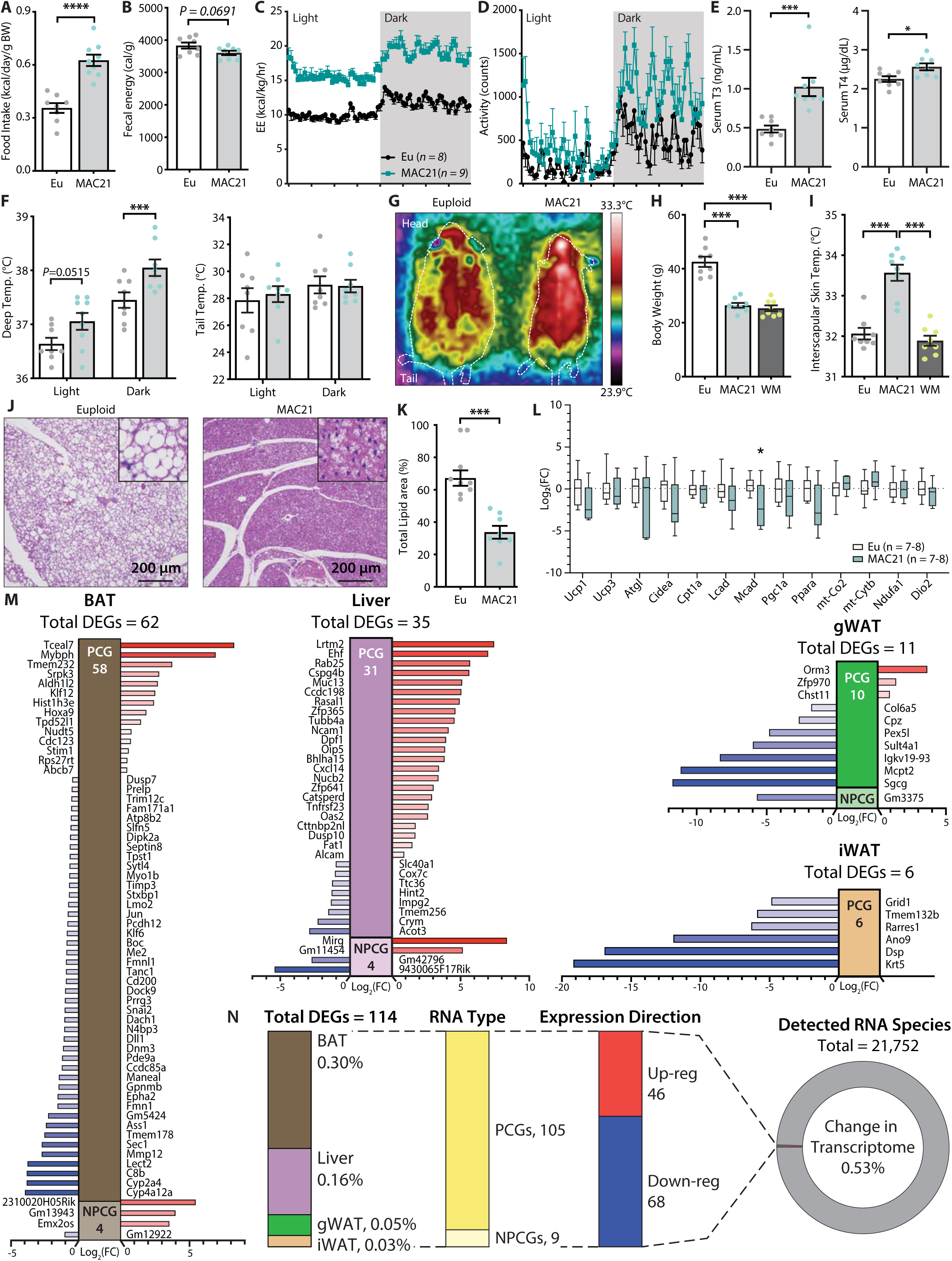
Hypermetabolism of TcMAC21 mice on HFD is uncoupled from changes in adipose and liver transcriptomes. **A)** Food intake in mice fed a high-fat diet (HFD). **B)** Fecal energy content. **C-D)** Energy expenditure (EE) and activity level over 24 h period in HFD-fed mice. **E)** Serum Triiodothyronine (T_3_) and Thyroxine (T_4_) levels. **F)** Deep colonic and tail temperature measured over three days in both the light and dark cycle. **G)** Representative infrared images of mice. **H)** Body weights of euploid, TcMAC21, and weight-matched (WM) control C57BL/6 mice. **I)** Interscapular skin temperature of euploid, TcMAC21, and WM control mice. **J)** Representative histology of brown adipose tissue (BAT). **K)** Quantification of percent total lipid area coverage per focal plane in BAT of euploid and TcMAC21. **L)** Expression of mouse genes (by qPCR) known to play major metabolic roles in BAT. **M)** Differentially expressed mouse genes (DEGs), both protein coding (PCG) and non-protein coding genes (NPCG) in BAT, liver, gonadal white adipose tissue (gWAT), and inguinal white adipose tissue (iWAT). All data is relative to euploid, and presented as Log2(FC). The list of genes shown is all the up and down regulated mouse genes (significant by adjusted p-value cut-off) for all 4 tissues. The red bars indicate up regulated genes and the blue bars indicate down regulated genes. **N)** General view and summary of transcriptional changes in BAT, Liver, gWAT, and iWAT to highlight the strikingly minimal changes in the mouse transcriptome across the four tissues. Only a combined total of 114 differentially expressed genes (DEGs) across four tissues, with the relative percentage (out of the 114 DEGs) shown for each tissue. Of the 114 DEGs, 105 are protein-coding genes (PCGs; dark yellow bar) and 9 are non-protein coding genes (NPCGs; light yellow bar). Of the 114 DEGs, 46 are upregulated (red bar) and 68 are down regulated (blue bar). In total, only a combined 0.53% change is noted in the transcriptome of all four tissues (out of the 21,752 RNAs detected).

Because TcMAC21 mice burned a large excess of energy, we measured the circulating levels of thyroid hormones as they are known to increase metabolic rate and energy expenditure (20). Both serum T_3_ (the active form) and T_4_ (precursor of T_3_) levels were significantly higher in TcMAC21 relative to euploid mice (Fig. 4E). T_3_ hormone, however, was not elevated in HFD-fed mice housed at thermoneutrality (30°C) (Fig. S1). If energy expenditure was elevated, body temperature of TcMAC21 mice would likely increase. Indeed, deep colon temperatures of TcMAC21 were elevated, most notably in the dark cycle when mice are active (Fig. 4F). Assessment with thermal imaging showed an elevated skin temperature around the interscapular region of TcMAC21 mice, whereas the tail skin temperature was not different between groups (Fig. 4F-G). Importantly, the differences in interscapular skin temperatures persisted in TcMAC21 even when compared to weight-matched wild-type mice (Fig. 4H-I), thus ruling out body weight (and hence surface area/volume ratio) as the cause of greater heat generation to compensate for greater heat loss. Elevated body temperature, however, was not observed in chow-fed mice (Fig. S7), even though chow-fed TcMAC21 were also hyperactive and had higher energy expenditure.

Consistent with the thermal imaging data, histological analysis of the interscapular brown adipose tissue (BAT) revealed a marked reduction in fat accumulation and a “healthy” brown appearance in TcMAC21 mice when compared to euploid controls (Fig. 4J-K), presumably due to excess lipids being utilized. We therefore expected several key metabolic genes in BAT to be upregulated in TcMAC21. Surprisingly, we found minimal differences in key thermogenic and fat oxidation genes between the two groups of mice (Fig. 4L and Fig. S8). This led us to assume the differences in gene expression must be due to non-canonical and potentially novel pathways. To test this, we conducted an unbiased RNA-sequencing analysis of BAT, liver, gWAT, and iWAT. Again, to our surprise and contrary to expectation, we found the transcriptomes of BAT, liver, gWAT, and iWAT in TcMAC21 to be remarkably similar to euploid controls (Fig. 4M).

There was not a single tissue that had more than 0.3% of its transcriptome significantly altered (Fig. 4N and Table S6-S9). None of the significant changes in gene expression—small in number—could readily account for the striking differences in phenotypes between TcMAC21 and euploid mice. Corroborating the RNA-seq results, quantitative real-time PCR analyses of select metabolic genes in BAT, liver, gWAT, and iWAT also showed minimal changes in TcMAC21 regardless of diet (chow or HFD) and temperature (22°C or 30°) (Fig. S8-12). Together, these data indicate that TcMAC21 mice are hyperactive and hypermetabolic with elevated thermogenesis, but these phenotypes are largely uncoupled from transcriptomic changes in BAT, liver, gWAT, and iWAT.

### Sarcolipin overexpression in skeletal muscle drives TcMAC21 hypermetabolism

Given the remarkable differences seen in body weight, adiposity, tissue histology and lipid contents, insulin sensitivity, body temperature, physical activity and metabolic rate between TcMAC21 and euploid mice, we were surprised by how few changes in transcriptomes—both in number and magnitude—were occurring in BAT, liver, and two major white adipose depots. This prompted us to examine the transcriptome of skeletal muscle, by far the largest metabolically active tissue that can substantially contribute to overall energy expenditure. RNA-sequencing of TcMAC21 skeletal muscle (gastrocnemius) revealed it to be the most transcriptionally dynamic tissue, with 4.2% of the transcriptome changed relative to euploid controls (Fig. 5A). Within the skeletal muscle of TcMAC21, there were 432 up-regulated genes and 501 down-regulated genes (Fig. S13 and Table S10). Gene ontology analysis of the differentially expressed genes highlighted up-regulated pathways related to thermogenesis, mitochondrial activity, and amino acid metabolism; and down-regulated pathways related to TGF-β, insulin, growth hormone, calcium, adrenergic, serine/threonine kinase, and cGMP-PKG signaling pathways (Fig. 5B).

**Figure 5.**
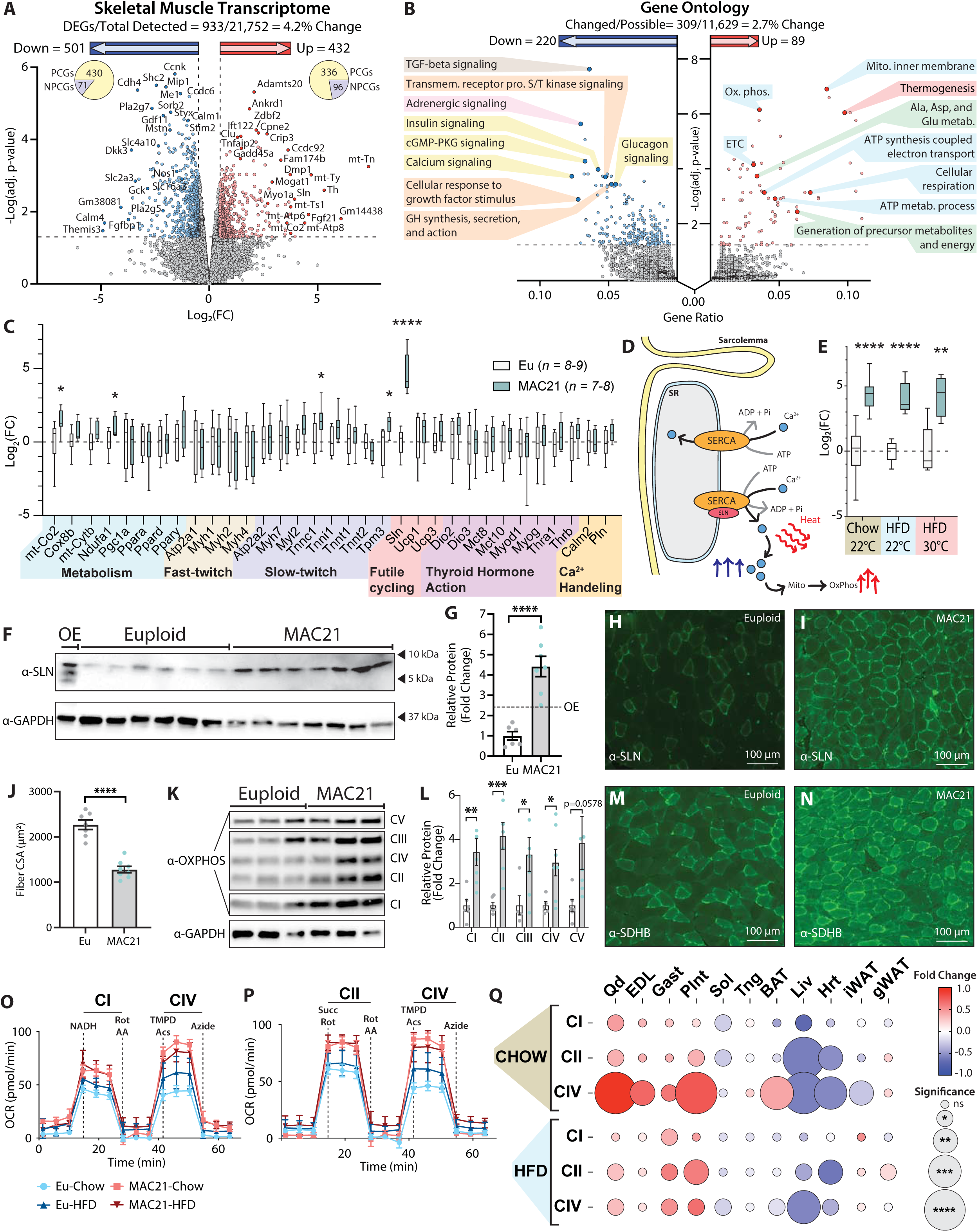
Sarcolipin overexpression in skeletal muscle drives TcMAC21 hypermetabolism. **A)** RNA-sequencing analysis reveals changes in TcMAC21 relative to euploid skeletal muscle (gastrocnemius). Volcano plot of skeletal muscle transcriptome (transcripts from the mouse genome only). The lower dotted line denotes significance at the adjusted *P* value cut-off (adj. *P* = 0.05). The vertical dotted lines denote a Log2(FC) of -0.5 or 0.5. **B)** Gene Ontology analysis of RNA-sequencing results using ClusterProfiler (77). **C)** Expression of genes (by qPCR) known to be involved in metabolism, fast- or slow-twitch fiber types, futile cycling, thyroid hormone action, and calcium handling in skeletal muscle (gastrocnemius). **D)** Graphical representation of the sarco(endo)plasmic reticulum Ca^2+^ ATPase (SERCA) and its regulator, sarcolipin (SLN). SERCA pump uses the energy derived from ATP hydrolysis to translocate Ca^2+^ from the cytosol back into the sarcoplasmic reticulum (SR), promoting muscle relaxation and restoring intracellular Ca^2+^ level following muscle contraction. SLN binds to SERCA and uncouples its Ca^2+^ transport activity from ATP hydrolysis and heat generation, thus promoting futile SERCA activity. **E)** qPCR analysis of sarcolipin (*Sln*) expression in euploid and TcMAC21 mice fed a standard chow, high-fat diet (HFD), and HFD while housed at thermoneutrality (30°C). **F)** Immunoblot of SLN and GAPDH (loading control) in muscle lysates of Euploid, TcMAC21, and SLN overexpression (OE) transgenic mice. **G)** Immunoblot quantification of SLN using GAPDH as a loading control. The dotted line marks the OE (SLN overexpression mouse model) level of SLN expression. **H-I)** Gastrocnemius immunofluorescent labeling of SLN-expressing muscle fibers. **J)** Muscle fiber cross-sectional area (CSA) quantification from WGA-stained gastrocnemius. **K)** Immunoblot of OXPHOS complex levels, with GAPDH as a loading control. **L)** Quantification of OXPHOS complex levels relative to GAPDH. **M-N)** Gastrocnemius immunofluorescent labeling of Succinate dehydrogenase subunit B (SDHB)-expressing muscle fibers. **O-P)** Seahorse respirometry analyses of frozen tissue samples. Shown here are the quadricep group average tracings for oxygen consumption rate (OCR) across the experimental time course. **Q)** OCR of TcMAC21 mitochondrial complex I, II, and IV relative to Euploid, for eleven separate tissues (quadricep, Qd; extensor digitorum longus, EDL; gastrocnemius, Gast; Plantaris, Plnt; Soleus, Sol; Tongue, Tng; Brown adipose tissue, BAT; Liver, Liv; Heart, Hrt; Inguinal white adipose tissue, iWAT; Gonadal white adipose tissue, gWAT) and two dietary conditions (Chow and HFD). DEGs, differentially expressed genes; PCGs, protein coding genes; NPCGs, non-protein coding genes. *n* = 5 euploid and 4 TcMAC21 for RNA-sequencing experiments. *n* = 8-9 euploid and 7-8 TcMAC21 for HFD qPCR. *n* = 7- 10 euploid and TcMAC21 for Chow qPCR. *n* = 4-5 euploid and TcMAC21 for HFD + thermoneutrality qPCR. *n* = 6 euploid and 6 TcMAC21 for all immunoblots. *n* = 7 Euploid and 8 TcMAC21 for gastrocnemius CSA quantification. *n* = 6 euploid and 4 TcMAC21 for all mitochondrial respiration assays; each biological replicate represents the average of three technical replicates.

To further confirm the RNA-sequencing results, we performed qPCR on a number of key genes related to skeletal muscle metabolism, fast- and slow-twitch fiber-types, futile-cycling, thyroid hormone action, and general calcium handling (Fig. 5C and Fig. S14). While a few of the genes were significantly different in expression, the most notable upregulated gene with the biggest magnitude of change was *Sln*, encoding the 31 amino acid single-pass membrane protein, sarcolipin (SLN). SLN is a regulator and an uncoupler of the sarco(endo)plasmic reticulum calcium ATPase (SERCA) (21, 22). Under normal circumstances, SERCA uses the energy derived from ATP hydrolysis to transport calcium from the cytosol back into the sarcoplasmic reticulum (SR) (23). When SLN binds to SERCA, it uncouples ATP hydrolysis from calcium transport into the SR (24); this results in futile SERCA pump activity, ATP hydrolysis, and heat generation (Fig. 5D) (22, 25, 26). In addition, increased cytosolic calcium transients (due to less being transported into the SR) promotes calcium entry into mitochondria and activates mitochondrial respiration, as well as, calcium-dependent signaling that enhances oxidative metabolism in skeletal muscle (27, 28).

Interestingly, regardless of diet (chow or HFD) and temperature (22°C or 30°C), TcMAC21 mice consistently had ∼20-30 fold upregulated *Sln* expression (Fig. 5E). The expression of other calcium handling genes—phospholamban (*Pln*), calmodulin (*Calm2*), SERCA1a (*Atp2a1*), SERCA2a (*Atp2a2*)—were not significantly different between TcMAC21 and euploid mice. Likewise, overexpressing SLN in skeletal muscle of transgenic mice also did not alter SERCA expression (29). The *Sln* gene is known to be upregulated in skeletal muscle by cold exposure (30–32). In contrast, the markedly upregulated *Sln* expression seen in TcMAC21 mice housed at ambient temperature (22°C) remained high even when the animals were housed at thermoneutrality (30°C). Consistent with the mRNA data, SLN protein levels were also strikingly upregulated in TcMAC21 skeletal muscle (gastrocnemius) (Fig. 5F-G). Immunofluorescence also indicated substantially more SLN positive muscle fibers in TcMAC21 mice (Fig. 5H-I). We included skeletal muscle lysate from SLN over-expression (OE) transgenic mouse (28) as our positive control. While the transgenic mouse had ∼2.5 fold higher level of SLN compared to controls, the SLN protein levels in TcMAC21 were ∼4.5 fold higher than the euploid mice (Fig. 5G). It is known that *Sln* can be induced in skeletal muscle as a compensatory response to muscle atrophy, dystrophy, and injury (33). However, none of the genes involved in muscle repair, wasting, and atrophy were significantly upregulated (Table S12), suggesting that *Sln* overexpression is not due to structural or functional deficit of the skeletal muscle in TcMAC21 mice. Collectively, our data suggest that a mechanism exists that can achieve upregulation of endogenous mouse SLN at a level substantially higher than that seen in artificially overexpressed transgenic mice under non-pathological condition.

Quantification of histological sections revealed that the average gastrocnemius muscle fiber cross-sectional area (CSA) in TcMAC21 was significantly smaller compared to euploid mice (Fig. 5J and Fig. S15). This suggests a shift from being a predominantly glycolytic muscle to a more oxidative muscle, as a smaller muscle fiber cross-sectional area is associated with a more oxidative muscle phenotype (34). In accordance, protein levels of mitochondrial complex I-V were significantly higher in TcMAC21 gastrocnemius muscle compared to euploid controls (Fig. 5K-L). Corroborating the Western blot data, immunofluorescence also showed substantially higher succinate dehydrogenase (SDHB) staining (marker of oxidative capacity) in gastrocnemius muscle of TcMAC21 mice (Fig. 5M-N).

To determine if TcMAC21 mice have higher mitochondrial activity compared to euploid controls, we conducted mitochondrial respiration analyses in liver, BAT, iWAT, gWAT, and different muscle types (quadriceps, extensor digitorum longus, gastrocnemius, plantaris, soleus, tongue, and heart). Except soleus and heart, most muscle types from TcMAC21 fed either chow or HFD—regardless of whether they were predominantly slow-twitch (oxidative), fast-twitch (glycolytic), or mixed—showed significantly elevated oxygen consumption due to elevated mitochondrial respiration (Fig. 4O-Q and Fig. S16-20). Enhanced mitochondrial complex I, II, and IV activities were more pronounced in glycolytic muscle tissues, suggesting that these normally glycolytic muscle fibers may assume a more oxidative phenotype in TcMAC21 mice. The greater oxidative phenotype in glycolytic muscle tissues was not due to changes in fiber type composition, as none of the fiber type-specific transcripts were different between genotypes (Fig. 5C). These data are consistent with similar results reported for skeletal muscle-specific SLN overexpression mice (27–29).

In contrast to muscle, mitochondrial activities of complex I and II in interscapular BAT was largely unchanged in TcMAC21 (Fig. 5Q and Fig. S18), despite the pronounced differences in BAT histology and lipid content between TcMAC21 and euploid mice (Fig. 4J-K). In chow-fed mice, however, mitochondrial complex IV activity was higher in BAT of TcMAC21. In subcutaneous (inguinal) white adipose tissue, mitochondrial respiration was significantly reduced in chow-fed TcMAC21 mice but unchanged in HFD-fed mice (Fig. S21). In visceral (gonadal) white adipose tissue, mitochondrial respiration was also largely unchanged in either chow or HFD-fed TcMAC21 mice (Fig. S21). Unexpectedly and in striking contrast to muscle and BAT, mitochondrial complex I, II, IV activity was markedly reduced in the liver and heart of both chow- and HFD-fed TcMAC21 compared to euploid mice (Fig. 5Q and Fig. S19-20), despite the liver of TcMAC21 mice being significantly “healthier” with much reduced lipid accumulation (Fig. 2H and 3E).

All together, these data suggest that SLN-mediated futile SERCA activity in skeletal muscle is likely the dominant driver underlying the hypermetabolism phenotypes seen in TcMAC21 mice. Because the futile SERCA pump activity consumes ATP to generate heat without transporting calcium into SR (22, 24, 26, 35, 36), this creates a huge energy demand and markedly drives up the activity of mitochondrial respiration to supply the ATP needed for the futile cycling of Ca^2+^. Presumably, this dominant effect results in systemic channeling of metabolic substrates (e.g., lipids) into skeletal muscle to fuel the elevated mitochondrial respiration rate. This in turn leads to secondary improvements and protection—reduced fat accumulation and smaller adipocyte—we observed in liver, BAT, gWAT, and iWAT despite minimal changes in the transcriptomes of these tissues.

### Potential regulators of sarcolipin expression in skeletal muscle

Having established that SLN-mediated futile SERCA activity underlies hypermetabolism in TcMAC21 mice, we next asked what regulator(s) promote the upregulation of endogenous SLN, as little is known about what controls its expression in skeletal muscle. We took advantage of the observation that SLN overexpression appears to be locked in the “on” position in skeletal muscle and could not be downregulated to control levels at thermoneutrality (Fig. 5E). Thyroid hormone (T_3_) is likely not the driver of SLN overexpression since serum T_3_ was not different between TcMAC21 and euploid housed at thermoneutrality (Fig. S1); in addition, SLN was shown to be upregulated in hypothyroid mice compared to euthyroid controls (37). When housed at thermoneutrality (30°C), TcMAC21 mice maintained their lean phenotypes with lower fasting insulin levels and elevated metabolic rate and energy expenditure despite higher caloric intake (Fig. 6A-E). Accordingly, TcMAC21 mice housed at thermoneutrality had smaller adipocyte cell size and lower lipid contents in BAT and liver (Fig. S22). Tissue collection at termination of the study also showed significantly smaller fat mass and reduced BAT and liver weight in TcMAC21 mice (Table S13).

**Figure 6.**
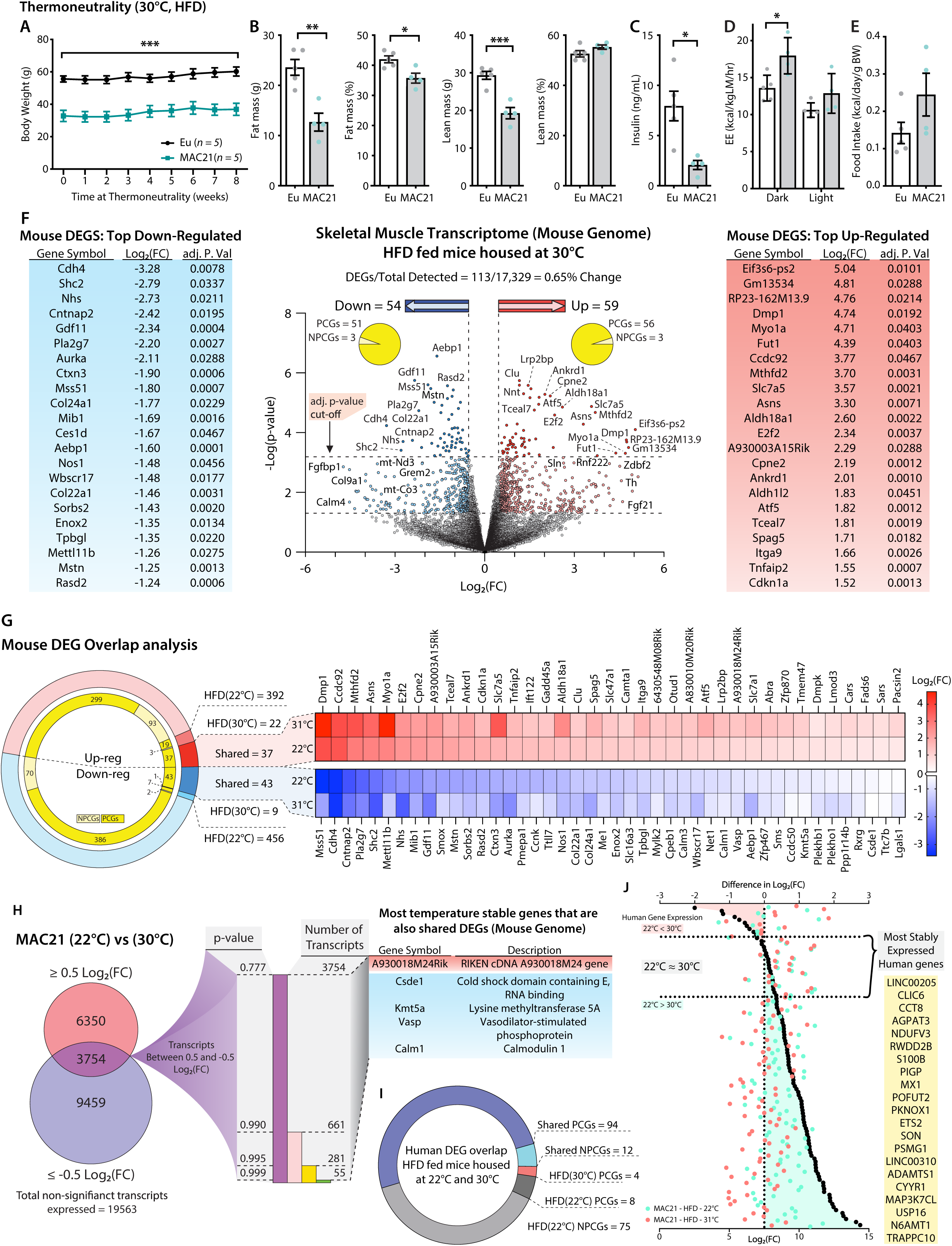
Potential regulators of sarcolipin expression in skeletal muscle. A.) Body weights at thermoneutrality (30°C) for 8 weeks. B) Body composition analysis of fat and lean mass. C) Fasting insulin levels. D) Energy expenditure in the dark and light cycles. E) Food intake. F) Volcano plot of skeletal muscle transcriptome (transcripts from the mouse genome only). The lower dotted line denotes significance at the *P* value cut-off (*P* = 0.05), and the upper dotted line denotes significance at the adjusted *P* value cut-off (adj. *P* = 0.05). The vertical dotted lines denote a Log 2 (Fold Change) of -0.5 or 0.5. Flanking the Volcano plot are the top down- and up-regulated mouse genes. G) Comparison of differentially expressed genes (DEGs) shared and not shared by TcMAC21 mice housed at 22°C vs. 30°C. Heat map showing all significantly up- or down-regulated shared genes. H) Direct comparison of TcMAC21 mice housed at 22°C and 30°C for most stably expressed mouse genes. Data filtered first by genes with Log2(FC) within ±0.5 (least change), then by lowest significance, and finally compared to the shared DEGs found in G. Table shows DEGs with least variation in expression at 22°C and 30°C. I) Overlap analysis of Hsa21-derived human transcripts expressed in TcMAC21 mice (skeletal muscle) housed at 22°C and 30°C. J) Graph showing the most stably expressed Hsa21-derived human genes in the TcMAC21 gastrocnemius. Top axis refers to the difference in Log2(FC) between human genes in TcMAC21 mice housed at 22°C and 30°C. Bottom axis shows the Log2(FC) value of a particular gene at 22°C and 30°C. The gene list shows all the human genes expressed by both groups with the least amount of change between the two temperatures.

From these data we surmised that the regulator(s) of SLN would likewise remain unchanged (i.e., continued to be up- or down-regulated) at thermoneutrality. We therefore performed RNA-sequencing on skeletal muscle isolated from HFD-fed TcMAC21 housed at thermoneutrality. RNA-seq data showed a total of 113 differentially expressed genes (DEGs) in the skeletal muscle of TcMAC21 mice, with 59 being up-regulated and 54 being down-regulated (Fig. 6F and Table S14). Of the 59 up-regulated genes, 56 were protein coding (PCGs) and 3 were non-protein coding (NPCGs); the down-regulated group had 51 PCGs and 3 NPCGs. Next, we compared the skeletal muscle transcriptomes of euploid vs TcMAC21 mice housed at 22°C with the transcriptomes of euploid vs TcMAC21 mice housed at 30°C. This tells us which genes are differentially expressed by TcMAC21 mice relative to euploid controls at both temperatures. There were 848 genes differentially expressed by only TcMAC21 mice housed at 22°C. This includes 392 up-regulated (299 PCGs and 93 NPCGs) and 456 down-regulated (386 PCGs and 70 NPCGs) genes (Fig. 6G). There were 31 genes differentially expressed by only TcMAC21 mice housed at 30°C, with 22 up-regulated (19 PCGs and 3 NPCGs) and 9 down-regulated (7 PCGs and 2 NPCGs) genes. Notably, there were 81 genes differentially expressed by TcMAC21 at both temperatures, with 37 up-regulated (all PCGs) and 44 down-regulated (43 PCGs and 1 NPCG) genes.

To further narrow our candidate *Sln* regulators, we directly compared the expression of mouse genes in the skeletal muscle of TcMAC21 mice housed at 22°C and 30°C. Because *Sln* expression was similarly upregulated in the skeletal muscle of TcMAC21 mice housed at 22°C and 30°C, we presumed the regulator(s) of *Sln* would also display minimal variation across both temperatures. We therefore focused on genes that remained unchanged at 22°C and 30°C in TcMAC21 skeletal muscle. We first filtered out all genes above or below Log2(FC) of 0.5 or -0.5, which left us with 3754 transcripts whose expression remained relatively unchanged at both temperatures. We next sorted the data by *P*-value, then compared the entire list of unchanged transcripts to the list of shared DEGs shown in Fig. 6G, and found that only 5 genes were present in both (Fig. 6H). This list contains one up-regulated gene (A930018M24Rik) and four down-regulated PCGs (*Csde1*, *Kmt5a*, *Vasp*, and *Calm1*). Again, these are the genes that are differentially expressed by TcMAC21 mice housed at 22°C and 30°C as compared to euploid, which remain unchanged in TcMAC21 mice across temperature. Thus, these are potential *Sln* regulators that are encoded in the mouse genome.

Alternatively, the regulator(s) of *Sln* expression could be one or more Hsa21-drived human genes. There was a total of 106 human transcripts expressed by TcMAC21 mice housed at 22°C and 30°C, with 94 PCGs and 12 are NPCGs (Fig. 6I). There were 4 human PCGs only expressed in TcMAC21 mice housed at 30°C and 83 human genes whose expression was turned off at 30°C. Interestingly, a majority (75 out of 83) of the human genes that were turned off at 30°C are NPCGs. The human genes that we postulated as most likely to regulate *Sln* expression would be those that change the least at 22°C and 30°C. We first calculated the difference in Log2(FC) values of all human genes expressed at 22°C and 30°C in TcMAC21 skeletal muscle [i.e., Log2(FC at 22°C) - Log2(FC at 30°C)]. Those values closest to zero represent genes with the most consistent expression across both housing temperatures. Genes with a negative Log2(FC) difference are expressed at higher levels at 30°C, and genes with a positive Log2(FC) difference are expressed at higher levels at 22°C. The ∼20 genes with the closest-to-zero Log2(FC) difference are listed in yellow (Fig. 6J), and these would be considered potential regulators of *Sln* derived from Hsa21. Altogether, our analysis has yielded a defined number of potential *Sln* regulators, from the mouse or human genome, that can be experimentally tested (either singly or in combination) in follow-up studies.

## DISCUSSION

While assessing the systemic metabolic impact of cross-species aneuploidy (a human chromosome in mice), we made several striking and unexpected discoveries. Given that TcMAC21 mice recapitulate multiple features of human trisomy 21 — most notably structural and functional neurological deficits and craniofacial skeletal alterations (18) and also a small body size compared to their euploid littermates —we initially expected the animals to show metabolic changes such as obesity and glucose intolerance reminiscent of the DS population (7) or the metabolic dysregulation (e.g., insulin resistance and dyslipidemia) seen in the Ts65Dn Down syndrome mouse model (11). Instead, the TcMAC21 mice display hallmarks of hypermetabolism. They are lean despite greater caloric intake, and have markedly elevated oxidative metabolic rate as indicated by increased VO_2_, energy expenditure, mitochondrial respiration in skeletal muscle, and by a shift to more a oxidative phenotype in skeletal muscle.

Although the higher prevalence of obesity and diabetes is well documented in the DS population (7), the mechanism of how gene dosage imbalance causes metabolic dysregulation *in vivo* is largely unknown and limited to only a few studies (10, 38). Until our recent study (11), the scope of the previous metabolic studies using DS mouse models was limited. Our present study represents one of the most comprehensive metabolic analyses carried out in DS mouse models to date. To our surprise, TcMAC21 phenotypes appear to be the opposite of what is seen in DS. We can speculate on possible reasons for this: 1) The strong and dominant physiological effects of *Sln* overexpression in the skeletal muscle of TcMAC21 override and mask potential deleterious effects of high-fat feeding seen in the euploid controls; 2) The unexpected phenotypes of TcMAC21 could be the result of complex interactions between human genes and the mouse genome that we do not fully understand. Further, the human proteins may alter the stoichiometric compositions of multi-protein complexes in an unpredictable way. For this reason, and for comparison, it is imperative to carry out similar comprehensive metabolic studies in other established DS mouse models (e.g., Dp(16)1Yey/+ and Ts66Yah) where all the trisomic genes are derived from the mouse instead of human, and also without extra trisomic genes not found in Hsa21 (39, 40). It is possible the trisomic mouse orthologs of Hsa21 genes are over-expressed and interact with the rest of the mouse genome in a manner that more closely reflects what might be seen in the adipose tissue, liver, and skeletal muscle of DS. This is an important issue to resolve in terms of selecting appropriate DS mouse models to best reflect the metabolic phenotypes seen in DS.

What could be the mechanism underlying the striking metabolic phenoytpes of TcMAC21? The key driver of hypermetabolism appears to be SLN overexpression in the skeletal muscle of TcMAC21 mice, leading to persistent uncoupling of the SERCA pumps, heat generation, and energy dissipation. This mechanism could explain multiple unexpected observations: despite a marked reduction in lipid accumulation in the liver, BAT, and WAT, surprisingly few changes were seen in the transcriptomes of these tissues, and none could account for the striking tissue histology. Importantly, none of the key genes (e.g., *Ucp1* in BAT and browning/beiging genes in iWAT) involved in thermogenesis and lipid metabolism were significantly different between TcMAC21 and euploid mice, and yet the mice had elevated deep colon temperature. While the metabolic activity (i.e., mitochondrial respiration) of BAT was largely unchanged, complex I, II, and IV activity in the liver were surprisingly downregulated. Mitochondrial respiration was either downregulated or unchanged in iWAT and gWAT. All these striking phenotypes could be explained by metabolic substrates (e.g., lipid) channeling away from liver, BAT, and WAT, and into skeletal muscle to fuel elevated mitochondrial respiration, as this was needed to meet the high ATP demand created by futile SERCA pump activity.

We largely ruled out the involvement of thyroid hormones in promoting hypermetabolism in TcMAC21 mice. Under the basal state when mice were fed a standard chow, or HFD-fed mice housed at thermoneutrality, circulating T_3_ levels were not different between groups despite the hyperactivity and hypermetabolism of TcMAC21 mice. Although serum T_3_ levels were elevated in HFD-fed TcMAC21 housed at room temperature (22°C), the predicted T_3_ effects were largely absent. It is known that T_3_ negatively regulates the expression of SLN, as hypothyroidic mice have significantly upregulated SLN mRNA and protein in skeletal muscle compared to euthyroid mice (37). Instead of the expected downregulation, SLN expression was dramatically elevated in the skeletal muscle of TcMAC21 mice. Through its nuclear hormone receptor, T_3_ regulates many metabolic genes in adipose tissue, liver, and skeletal muscle (20), and yet none of the well-known T_3_-regulated genes were upregulated in these tissues despite elevated T_3_ levels. In addition, T_3_ is a potent inducer of *Ucp1* expression in BAT (41) and white adipose tissue beiging (42), but *Ucp1* transcript levels were not upregulated in BAT or the white adipose tissue of TcMAC21 mice. This apparent T_3_ resistance in HFD-fed TcMAC21 mice is likely a compensatory response to an already elevated body temperature and the hypermetabolism induced by SLN-mediated futile SERCA activity and heat generation in skeletal muscle. If T_3_ further increased the metabolic rate of TcMAC21 mice, this would likely overtax the heat dissipation mechanism leading to hyperthermia and possible death.

By collapsing the proton motive force within the intermembrane space of mitochondria, UCP1-mediated mitochondrial uncoupling in BAT could raise body temperature and enhance energy expenditure (43). Although thermal imaging highlighted increased skin temperature around the interscapular BAT region of TcMAC21 mice, the thermal signal could be produced by local muscle around the interscapular area. Our gene expression, BAT transcriptomics, and functional data, however, do not support UCP1-mediated uncoupling in BAT as a probable mechanism for the hypermetabolism and elevated thermogenesis seen in TcMAC21 mice. Neither the expression of *Ucp1* nor mitochondrial activity was consistently elevated in the BAT of TcMAC21 mice fed chow or HFD. Housing mice at a thermoneutral temperature would suppress BAT activity and beigeing in iWAT (44), yet TcMAC21 mice housed at thermoneutrality for an extended period (>2 months) retained elevated energy expenditure and lean body weight comparable to TcMAC21 mice housed at 22°C. Together, these data argue against BAT being a significant contributor to the thermogenesis seen in TcMAC21 mice.

Recent studies have also shown that creatine-driven futile substrate cycling and SERCA2b-mediated calcium cycling in beige fat can promote energy expenditure and thermogenesis (45, 46). Regardless of housing temperatures, none of the genes involved in creatine/phosphocreatine futile cycle (e.g, *Slc6a8*, *Gatm*, *Gamt*, *Ckmt1*, *Ckmt2*) or the SERCA2b-RyR2 pathway (*Atp2a2*, *Ryr2*) were upregulated in iWAT, suggesting that these pathways are likely not involved in the hypermetabolic phenotypes of TcMAC21. Promoting metabolic inefficiency in skeletal muscle by UCP1 or UCP3 overexpression can increase metabolic rate and prevent diet-induced obesity (47–49). In the skeletal muscle of TcMAC21 mice, *Ucp1* and *Ucp3* transcripts were not upregulated, suggesting that mitochondrial uncoupling by UCP1 or UCP3 is also unlikely to contribute to the observed lean and hypermetabolic phenotypes.

SLN plays a role in modulating body weight by promoting energy expenditure through its effects on the SERCA pump (25, 28, 50, 51). Binding of SLN to SERCA does not interfere with ATP hydrolysis of the pump, but reduces calcium transport into SR lumen through a calcium “slippage” mechanism (22, 26, 35, 36). Thus, SLN promotes futile cycling of the SERCA pumps, ATP hydrolysis, and heat generation. Consistent with this model, mice lacking SLN are cold intolerant and gain more weight on an HFD due to reduced energy expenditure (25, 50, 51); conversely, transgenic overexpression of SLN in skeletal muscle promotes a lean phenotype (28).

Under normal situations, SLN is mainly expressed in slow-twitch oxidative muscle tissues such as the soleus and diaphragm, with little expression in fast-twitch glycolytic muscle tissues such as the gastrocnemius and quadriceps (52). In the case of hypothyroid mice, endogenous SLN is only markedly upregulated in oxidative muscle fibers (soleus and diaphragm) but not the large glycolytic muscle tissues (37). It has been argued that in rodents the futile SERCA activity and energy expenditure occurring in soleus and diaphragm (both are small in size) is quantitatively insufficient to effect significant changes in body weight (53). Notwithstanding the artificial overexpression of SLN in both fast-twitch glycolytic and slow-twitch oxidative muscle fibers (28), it is unclear whether endogenous SLN can be substantially induced in large glycolytic muscle tissues such as the gastrocnemius and quadriceps under non-pathological conditions. Our data helps to resolve this issue. In TcMAC21 mice, endogenous SLN is markedly upregulated in a largely glycolytic muscle (gastrocnemius), suggesting that a mechanism indeed exists that can substantially elevate SLN transcript and protein in muscle fiber types that normally have low expression. Corroborating this is the observation that mitochondrial respiration is significantly elevated in most muscle types—gastrocnemius (fast-twitch), quadriceps (fast-twitch), plantaris (mixed), and extensor digitorum longus (mixed).

The huge ATP demand created by the futile SERCA pump activity appears to be the cause of upregulated protein expression of complex I-IV, elevated mitochondrial respiration, and the assumption of an oxidative phenotype in muscle tissues that are normally glycolytic. This reprogramming of glycolytic muscle tissues into an oxidative phenotype is likely due to elevated cytosolic calcium transients (as less calcium is being transported into the SR due to uncoupling of the SERCA pumps), leading to calcium-dependent activation of mitochondrial respiration and calcium-dependent signaling that drives increased production of OXPHOS proteins (27, 28, 54–57). Since skeletal muscle makes up ∼40% of the body’s weight and is a major energy consuming tissue (58, 59), elevated energy expenditure via futile SERCA pump activity—especially the big muscles (gastrocnemius and quadriceps)—will have substantial impact on total energy balance and body weight. As an added benefit, increased fuel consumption for non-shivering thermogenesis driven by the futile SERCA pump would channel lipid substrates into skeletal muscle for oxidation and prevent excess lipid accumulation in liver, BAT, and white adipose tissue, thus conferring an overall healthy metabolic profile with preserved systemic insulin sensitivity.

Although TcMAC21 mice also have increased voluntary physical activity, our data suggest that futile SERCA activity in skeletal muscle is the main driver of the hypermetabolic phenotype. Overexpression of SLN in skeletal muscle does not significantly alter voluntary physical activity at ambient room temperature or at thermoneutrality, yet the SLN transgenic mice consistently gained significantly less weight in response to high-fat feeding (28, 50). Further, SLN appears to be required for effective adiposity reduction in HFD-fed mice with access to a running wheel (60). Elevated locomotor activity in TcMAC21 mice, however, would likely amplify the effect of sarcolipin-mediated uncoupling of the SERCA pumps, as muscle movement results in greater SR Ca^2+^ cycling. In cultured myotubes, caffeine-induced release of Ca^2+^ from the SR via the ryanodine receptor/channel (RYR1) would only activate calcium-dependent signaling in the presence, but not absence, of SLN; conversely, inhibiting Ca^2+^ release through RYR1 using dantrolene blocks calcium-dependent signaling only in SLN-expressing, but not SLN-deficient, myotubes (27). Thus, increased SR Ca^2+^ cycling due to elevated physical activity would promote energy expenditure and also create a favorable cellular context to amplify the effects of sarcolipin overexpression on uncoupling of the SERCA pumps. Accordingly, energy expenditure and the degree of leanness seen in TcMAC21 mice are significantly greater in magnitude than the SLN transgenic mice.

To fully harness SLN-mediated futile SERCA activity for energy dissipation and weight loss requires an understanding of how endogenous SLN expression is regulated, but the mechanism of which remains largely unknown. TcMAC21 mice offer critical insights into the switches that control SLN expression in skeletal muscle. Regardless of temperatures (ambient or thermoneutral), elevated *Sln* expression in TcMAC21 appeared to be in the “on” position and could not be turned off. We exploited this information to zoom in on a defined set of mouse and Hsa21-derived transcripts—some of which are transcription factors (e.g., *PKNOX1*, *ETS2*), RNA binding proteins (e.g., *Csde1*), and epigenetic regulators (e.g., *Kmt5a*)—that remain significantly up- or down-regulated in skeletal muscle at both 22°C and 30°C, and show <1% variability across the two temperatures. We presume that one or more of these genes could potentially be the sought after “switches” of *Sln* expression. Additional studies are needed to identify such critical on/off switches. A recent study has suggested that the C-terminal cleavage fragment of TUG can upregulate *Sln* expression in the skeletal muscle when it enters the nucleus and forms a complex with PPAR-γ and PGC-1α (61). However, this pathway is largely abrogated in mice fed an HFD; thus, we think this transcriptional mechanism is likely not involved in upregulating *Sln* expression in TcMAC21 mice fed an HFD.

Weight loss can be achieved by reducing caloric intake or promoting energy expenditure, and the latter can be physiologically accomplished by increasing physical activity (e.g., exercise) or through shivering and non-shivering thermogenesis (43, 62). SLN-mediated non-shivering thermogenesis offers an attractive approach for promoting energy expenditure and weight loss: Expression of SLN is restricted to the striated muscle of all mammals, including humans (63–66). Compared to BAT (∼1.5% of body weight in young men) (67), the total mass of skeletal muscle (∼40% of body weight) makes futile SERCA activity in this large tissue especially effective in energy dissipation (68). Importantly, overexpression of SLN in skeletal muscle does not appear to have adverse effects; rather, SLN transgenic mice have higher endurance capacity and improved muscle performance due to enhanced oxidative capacity (29). In summary, our work provides further proof-of-concept that endogenous SLN-mediated uncoupling of SERCA pumps to enhance energy expenditure can potentially be harnessed for systemic metabolic health.

## MATERIALS AND METHODS

### Mouse model

The transchromosomic TcMAC21 (simply referred to as MAC21) mice carrying a near complete human chromosome 21were generated and genotyped as previously described (18). Because MAC21 male mice are infertile, all the female mice were used for breeding. Hence, metabolic analyses were conducted on male mice only. MAC21 mice were initially maintained on an outbred background (ICR strain mice). After crossing for eight generations onto BDF1 (C57BL/6J (B6) x DBA/2J (D2)), MAC21 were transferred to the Riken Animal Resource (BRC No. RBRC05796 and STOCK Tc (HSA21q-MAC1)). Characterizations of MAC21 were performed on mice (75% B6/25% D2 on average) produced by crossing B6 males with trisomic B6D2 females (18). Euploid littermates were used as controls throughout the studies.

Mice were fed a standard chow (Envigo; 2018SX) or high-fat diet (HFD; 60% kcal derived from fat, #D12492, Research Diets, New Brunswick, NJ). Mice were housed in polycarbonate cages on a 12h:12h light-dark photocycle with ad libitum access to water and food. Standard chow was provided for 8 weeks, beginning at 12 weeks of age; HFD was provided for 16 weeks, beginning at 5 months of age. At termination of the study, all mice were fasted for 2 h and euthanized. Tissues were collected, snap-frozen in liquid nitrogen, and kept at 80°C until analysis.

For thermoneutral studies, mice were housed in a temperature regulated (ambient temp. maintained at 30 ± 1°C) animal facility at Johns Hopkins University School of Medicine. Mice were acclimatized for two full weeks prior to experimentation. All experimental procedures conducted on these mice (i.e., glucose and insulin tolerance tests, fasting-refeeding blood collections, tissue dissections, etc.) were completed within the 30°C housing unit to prevent temperature fluctuation.

All mouse protocols were approved by the Institutional Animal Care and Use Committee of the Johns Hopkins University School of Medicine (animal protocol # MO19M481). All animal experiments were conducted in accordance with the National Institute of Health guidelines and followed the standards established by the Animal Welfare Acts.

### Body composition analysis

Body composition analyses for total fat and lean mass, and water content were determined using a quantitative magnetic resonance instrument (Echo-MRI-100, Echo Medical Systems, Waco, TX) at the Mouse Phenotyping Core facility at Johns Hopkins University School of Medicine.

### Complete blood count analysis

A complete blood count on blood samples was performed at the Pathology Phenotyping Core at Johns Hopkins University School of Medicine. Tail vein blood was collected using EDTA-coated blood collection tubes (Sarstedt, Nümbrecht, Germany) and analyzed using Procyte Dx analyzer (IDEXX Laboratories, Westbrook, ME).

### Indirect calorimetry

Chow or HFD-fed MAC21 male mice and euploid littermates were used for simultaneous assessments of daily body weight change, food intake (corrected for spillage), physical activity, and whole-body metabolic profile in an open flow indirect calorimeter (Comprehensive Laboratory Animal Monitoring System, CLAMS; Columbus Instruments, Columbus, OH) as previously described (69). In brief, data were collected for three days to confirm mice were acclimatized to the calorimetry chambers (indicated by stable body weights, food intakes, and diurnal metabolic patterns), and data were analyzed from the subsequent three days. Mice were observed with ad libitum access to food, throughout the fasting process, and in response to refeeding. Rates of oxygen consumption (*V̇* _O2_; mL·kg^-1^·h^-1^) and carbon dioxide production (*V̇* _CO2_; mL·kg^-1·^h^-1^) in each chamber were measured every 24 min. Respiratory exchange ratio (RER = *V̇* _CO2_/*V̇* _O2_) was calculated by CLAMS software (version 4.93) to estimate relative oxidation of carbohydrates (RER = 1.0) versus fats (RER = 0.7), not accounting for protein oxidation. Energy expenditure (EE) was calculated as EE= *V̇* _O2_× [3.815 + (1.232 × RER)] and normalized to lean mass. Physical activities (total and ambulatory) were measured by infrared beam breaks in the metabolic chamber. Average metabolic values were calculated per subject and averaged across subjects for statistical analysis by Student’s t-test.

### Fecal bomb calorimetry and assessment of fecal parameters

Fecal pellet frequency and average fecal pellet weight were monitored by housing each mouse singly in clean cages for 3 days and counting the number of fecal pellets and recording their weight at the end of each 24 h period. The average of the three days was used to generate a mouse average, which were then averaged within a group for comparison across genotype. Fecal pellets from 3 days were combined and shipped to the University of Michigan Animal Phenotyping Core for fecal bomb calorimetry. Briefly, fecal samples were dried overnight at 50°C prior to weighing and grinding them to powder. Each sample was mixed with wheat flour (90% wheat flour, 10% sample) and formed into 1.0 g pellet, which was then secured into the firing platform and surrounded by 100% oxygen. The bomb was lowered into water reservoir and ignited to release heat into the surrounding water. Together these data were used to calculate fecal pellet frequency (bowel movements/day), average fecal pellet weight (g/bowel movement), fecal energy (cal/g feces), and total fecal energy (kcal/day).

### Thermography tests

Deep colonic temperature was measure by inserting a lubricated (Medline, water soluble lubricating jelly, MDS032280) probe (Physitemp, BAT-12 Microprobe Thermometer) into the anus at a depth of 2 cm. Stable numbers were recorded on three separate days in both the dark and light cycle for each mouse. Skin temperature measurements and images of the tail, abdomen, and suprascapular regions of mice were taken across three days, in both the dark and light cycle, using a thermal imaging camera (Teledyne FLIR, Sweden, FLIR-C2) set at a constant distance of ∼16 inches away from the specimen.

### Bone length measurements

Tibias were dissected and excess tissue was cleared away. A Mitutoyo Corp. digital caliper (500-196-30) was used to measure end-to-end bone length.

### Glucose and insulin tolerance tests

For glucose tolerance tests (GTTs), mice were fasted for 6 h before glucose injection. Glucose (Sigma, St. Louis, MO) was reconstituted in saline (0.9 g NaCl/L), sterile-filtered, and injected intraperitoneally (i.p.) at 1 mg/g body weight. Blood glucose was measured at 0, 15, 30, 60, and 120 min after glucose injection using a glucometer (NovaMax Plus, Billerica, MA). Blood was collected at 0, 15, and 30 min time points for serum isolation followed by Insulin ELISA. For insulin tolerance tests (ITTs), food was removed 2 h before insulin injection. Insulin was diluted in saline, sterile-filtered, and injected i.p. at 1.0 U/kg body weight. Blood glucose was measured at 0, 15, 30, 60, and 90 min after insulin injection using a glucometer (NovaMax Plus).

### Fasting-Refeeding insulin tests

Mice were fasted overnight (∼16 h) then reintroduced to food. Blood glucose was monitored at the 16 h fast time point (time = 0 h refed) and at 1, 2, and 3 hours into the refeeding process. Blood was collected at the 16 h fast and 2 h refed time points for insulin ELISA. Homeostatic model assessment for insulin resistance (HOMA-IR) was calculated as follows (70): [fasting insulin (uIU/mL) X blood glucose (mmol/L)] / 22.5.

### Blood and tissue chemistry analysis

Tail vein blood samples were allowed to clot on ice and then centrifuged for 10 min at 10,000 x *g*. Serum samples were stored at -80°C until analyzed. Serum triglycerides (TG) and cholesterol were measured according to manufacturer’s instructions using an Infinity kit (Thermo Fisher Scientific, Middletown, VA). Non-esterified free fatty acids (NEFA) were measured using a Wako kit (Wako Chemicals, Richmond, VA). Serum β-hydroxybutyrate (ketone) concentrations were measured with a StanBio Liquicolor kit (StanBio Laboratory, Boerne, TX). Serum insulin (Crystal Chem, 90080), adiponectin (MilliporeSigma, EZMADP-60K), leptin (MilliporeSigma, EZML-82K), growth hormone (MilliporeSigma, EZRMGH-45K), IGF-1 (Crystal Chem, 80574), T3 (Calbiotech, T3043T-100), and T4 (Calbiotech, T4044T-100) levels were measured by ELISA according to manufacturer’s instructions. Pancreatic insulin and somatostatin (SST) were isolated from pancreas samples using acid-ethanol extraction. Briefly, pancreatic samples were in a solution of acid-ethanol (1.5% HCl in 70% EtOH) and incubated overnight at -20°C. They were then homogenized and incubated for an additional night at - 20°C. The following day, samples were centrifuged at 4°C to pellet debris. The aqueous protein-rich solution was transferred to a new tube and neutralized with 1M Tris pH7.5. Prior to Insulin quantification by ELISA, samples were diluted 1:1000 with ELISA sample diluent. Following ELISA quantification of pancreatic insulin (Mercodia, 10-1247-01) and SST (Phoenix Pharmaceuticals Inc., EK-060-03) protein concentrations of pancreatic samples were quantified using a Bradford assay (Sigma-Aldrich, B6916). Insulin and SST values were then normalized to protein concentration, averaged, and compared across groups.

### Histology and quantification

Inguinal (subcutaneous) white adipose tissue (iWAT), gonadal (visceral) white adipose tissue (gWAT), liver, pancreas, suprascapular brown adipose tissue (BAT), and gastrocnemius muscle were dissected and fixed in formalin. Paraffin embedding, tissue sectioning, and staining with hematoxylin and eosin were performed at the Pathology Core facility at Johns Hopkins University School of Medicine. Images were captured with a Keyence BZ-X700 All-in-One fluorescence microscope (Keyence Corp., Itasca, IL). Adipocyte (gWAT and iWAT) and gastrocnemius cross-sectional area (CSA), as well as the total area covered by lipid droplets in hepatocytes and brown adipose tissue adipocytes, were measured on hematoxylin and eosin-stained slides using ImageJ software (71). For CSA measurements, all cells in one field of view at 100X magnification per tissue section per mouse were analyzed. For iWAT and gWAT adipocyte CSA, at least 150 cells were quantified per mouse. For pancreatic β-islet CSA, 40X magnification images were stitched together from sections at two depths of the pancreas for a single mouse. All β-islets across the two stitched images were quantified per mouse. Image capturing and quantifications were carried out blinded to genotype.

### Immunohistochemistry and quantification

All paraffin sections used for immunofluorescence staining were deparaffinized with SafeClear (Fisher HealthCare, 23-044192), rehydrated using graded concentrations of ethanol in water (100% EtOH, 95%, 75%, 50%, then dH_2_O), then subjected to antigen retrieval with a pressure cooker for 5 min in sodium citrate buffer (10 mM sodium citrate, 0.05% Tween 20, pH 6.0). Specimens were washed with a TBS + 0.025% Trion X-100 buffer, blocked with 10% normal goat serum at RT for 1 h, and incubated at 4°C overnight with the appropriate primary antibody. Muscle cells expressing sarcolipin or oxidative fibers were labeled with a rabbit polyclonal anti-sarcolipin Ab (Millipore Sigma, ABT13) or mouse monoclonal Ab to SDHB (Abcam, ab14714), respectively. Primary antibodies were used at a concentration of 1:200 in TBS buffer (with 1% BSA), followed by the appropriate secondary antibody at 1:1000 (diluted in TBS + 1% BSA) and incubated for 1 h at RT. Secondary antibodies used were goat anti-mouse (Invitrogen, Alexa Fluor 594, A21135) and goat anti-rabbit (Invitrogen, Alexa Fluor 488, A11008). All specimens were additionally stained with WGA (Wheat Germ Agglutinin, Alexa Fluor 647, W32466) at a concentration of 1:250 in TBS and mounted with a coverslip using ProLong Gold antifade reagent with DAPI (Invitrogen, P36935). Images were captured with a Keyence BZ-X700 All-in-One fluorescence microscope (Keyence Corp., Itasca, IL). Muscle cell cross-sectional area analysis was completed on specimens stained with WGA. At least 1000 cells of the gastrocnemius muscle were quantified per mouse using ImageJ (71).

### Western blots analysis

Protein was isolated from skeletal muscle samples using RIPA buffer as previously described (72). Protein lysates used for sarcolipin immunoblots were boiled for 5 min in a loading buffer (50 mM Tris, 2% SDS, 1% β-ME, 6% glycerol, 0.01% bromophenol blue), while those used for mitochondrial oxidative phosphorylation complex immunoblots were heated for 3 min at 50°C in the same buffer. Total protein was quantified by BCA assay (Thermo Scientific, 23225), loaded in equal amounts and volume, and run on a 4-20% gradient gel (Biorad, 4561096). Protein was transferred to nitrocellulose or PVDF membrane (sarcolipin and OXPHOS blots, respectively) and blocked in PBS containing 0.2% Tween 20 and 5% non-fat milk for 1 h, then incubated overnight at 4°C on a shaker with the antibody. Sarcolipin was detected using the Millipore Sigma antibody (ABT13, rabbit polyclonal) at a concentration of 1:500. Mitochondrial oxidative phosphorylation complexes were detected with the Abcam OXPHOS cocktail antibody (ab110413, mouse monoclonal) at a concentration of 1:5000. GAPDH was detected using the Proteintech antibody (60004-1-Ig, mouse monoclonal) at a concentration of 1:20,000. Anti-rabbit or anti-mouse secondary antibodies conjugated to HRP were used to recognize the primary antibody. Immunoblots were developed using HRP substrate ECL (GE Healthcare), visualized with a MultiImage III FluorChem Q (Alpha Innotech), and quantified with ImageJ. For the OXPHOS membrane, the blot was re-probed with the GAPDH antibody after stripping with ReBlot Plus Strong Antibody Stripping Solution (MilliporeSigma, 2504).

### Electron microscopy and quantification

Mouse pancreas was dissected and sectioned into six pieces (∼1 mm^3^) for fixation with freshly prepared electron microscopy-grade 2% paraformaldehyde, 2% glutaraldehyde in 100 mM Sorenson’s phosphate buffer containing 3 mM MgCl_2_, pH 7.4, and 1,144 mOsm overnight at 4 °C on slow rocker. Samples were rinsed with the same buffer containing 3% sucrose, 316 mOsm, and osmicated in 2% osmium tetroxide reduced in 1.5% potassium ferrocyanide for 2 h at 4°C. Tissue was then rinsed in 100 mM maleate buffer containing 3% sucrose pH 6.2 and en bloc stained in 2% uranyl acetate in maleate buffer for 1 h at 4°C in the dark. Samples were dehydrated in a graded ethanol series, brought to room temperature in 70% ethanol, and completely dehydrated in 100% ethanol. Samples were resin-embedded (Epon 812, T. Pella) after a propylene oxide transition step and further infiltrated and cured the next day. 80-nm-thin compression free sections were obtained with a Diatome diamond knife (35 degree). Sections were picked up on 1 x 2-mm formvar-coated copper slot grids (Polysciences) and further stained with uranyl acetate followed by lead citrate. Grids were examined on a Hitachi H-7600 transmission electron microscope operating at 80 kV. Images of β-cells and acinar cells were digitally captured at magnifications of 5,000 and 10,000X from six unique locations in the pancreas of each mouse with an AMT XR 50-5 megapixel CCD camera. Within each of the six locations, at least 600 zymogen granules and 200 insulin granules (dense insulin core and vesicle) were measured for CSA quantification using ImageJ software. Analyses were performed on a total of 6 randomly selected male chow-fed mice (WT, *n* = 3; KO, *n* = 3) and 6 randomly selected HFD-fed mice (WT, *n* = 3; KO, *n* = 3). Of the six unique locations from a single mouse, values were averaged to generate a mouse average. Therefore, each data point represents a mouse average comprised of at least 3,600 zymogen granules, 1,200 dense insulin cores, or 1,200 insulin vesicles.

### Quantitative real-time PCR analysis

Total RNA was isolated from tissues using Trizol reagent (Thermo Fisher Scientific) according to the manufacturer’s instructions. Purified total RNA was reverse transcribed using an iScript cDNA Synthesis Kit (Bio-rad). Real-time quantitative PCR analysis was performed on a CFX Connect Real-Time System (Bio-rad) using iTaq^TM^ Universal SYBR Green Supermix (Bio-rad) per manufacturer’s instructions. Data were normalized to *36B4* gene (encoding the acidic ribosomal phosphoprotein P0) and expressed as relative mRNA levels using the ΔΔCt method (73). Fold change data were log transformed to ensure normal distribution and statistics were performed. Real-time qPCR primers used are listed in supplemental Table S14.

### RNA-sequencing analysis

A total of 50 samples were sequenced with paired-end 50-bp reads (2×50bp) across two batches that were balanced for genotype (*n* = 40 across liver, brown fat, subcutaneous fat and visceral fat and *n* = 10 of skeletal muscle). Raw sequencing reads from all samples were aligned to a custom concatenated Gencode hg38 + mm10 reference genome using HISAT2 2.0.4 (74). Genes were quantified with featureCounts v1.5.0-p3 (75) to a custom concatenated hg38+mm10 gtf file within each sample. We retained 24,016 genes with mean RPKM > 0.1 across the mouse genome (N=23,801 genes) and chr21 in the human genome (N=215 genes) above this cutoff, and subsequently dropped two samples with outlier RNA-seq quality metrics (one SubQ_MAC21 sample with low gene assignment and high mitochondrial rates and one Skeletal_MAC21 sample with low read alignment rate), leaving a total of 48 samples. High-throughput sequencing data from this study have been submitted to the NCBI Sequence Read Archive (SRA) under accession number PRJNA877694.

We interrogated the effects of MAC21 genotype, tissue, and the statistical interaction between genotype and tissue on the transcriptome, further adjusting for confounders of gene assignment rate and mitochondrial mapping rate, using the limma voom approach (76). Given that four of the tissues were derived from the same animals, we used linear mixed effect using the animal ID as a random intercept. We accounted for multiple testing by controlling for the Benjamini-Hochberg false discovery rate across all mouse genes (since human genes were highly enriched for being DEGs given these defined the genotype effect). As a secondary analysis, we modeled the effect of MAC21 genotype within each tissue (also adjusting for the same two sequencing-derived confounders as the full model) using linear regression (since there were no repeated measures within a tissue). Gene set enrichment analyses were performed using clusterProfiler (77) and accounted for the false discovery rate.

### Respirometry of frozen tissue samples

Respirometry was conducted on frozen tissue samples to assay for mitochondrial activity as described previously (78). Briefly, all tissues were dissected, snap frozen in liquid nitrogen, and stored at -80°C freezer for later analysis. Samples were thawed in MAS buffer (70mM sucrose, 220 mM mannitol, 5 mM KH_2_PO_4_, 5 mM MgCl_2_, 1 mM EGTA, 2 mM HEPES pH 7.4), finely minced with scissors, then homogenized with a glass Dounce homogenizer. The resulting homogenate was spun at 1000 *g* for 10 min at 4°C. The supernatant was collected and immediately used for protein quantification by BCA assay (Thermo Scientific, 23225). Each well of the Seahorse microplate was loaded with 8 µg of homogenate protein for all tissue types, except for brown adipose tissue (BAT) and heart of which 4 µg was loaded. Each biological replicate is comprised of three technical replicates. Samples from all tissues were treated separately with NADH (1 mM) as a complex I substrate or Succinate (a complex II substrate, 5 mM) in the presence of rotenone (a complex I inhibitor, 2 µM), then with the inhibitors rotenone (2 µM) and Antimycin A (4 µM), followed by TMPD (0.45 mM) and Ascorbate (1 mM) to activate complex IV, and finally treated with Azide (40 mM) to assess non-mitochondrial respiration.

### Statistical analyses

All results are expressed as mean ± standard error of the mean (SEM). Statistical analysis was performed with Prism 9 software (GraphPad Software, San Diego, CA). Data were analyzed with two-tailed Student’s *t*-tests or by repeated measures ANOVA. For two-way ANOVA, we performed Bonferroni post hoc tests. *P* < 0.05 was considered statistically significant.

## ACKNOWLEDGEMENTS

The work was funded, in part, by grants from the National Institutes of Health (DK084171 to GWW, HD098540 and HD038384 to RHR), Japan Society for the Promotion of Science (25221308 to MO), and the Core Research for Evolutional Science and Technology (JPMJCR18S4 to YK). The presented information and its interpretation do not necessarily reflect those of the funding agencies. We thank Nanami Senoo from the Claypool lab for technical advice on conducting the mitochondrial respiration experiments using Seahorse machine.

## AUTHOR CONTRIBUTIONS

DCS, RHR, GWW conceived the project. DCS, CX, SR, SA, MD, GWW performed the experiments. DCS, SA, AJ, GWW analyzed and interpreted the data. FG bred and maintained the mouse lines. YK, MO, and RHR provided the TcMAC21 mouse model. MP provided reagents and inputs to the project. GWW and DCS wrote the manuscript with inputs from all authors.

---

**Figure S1.**
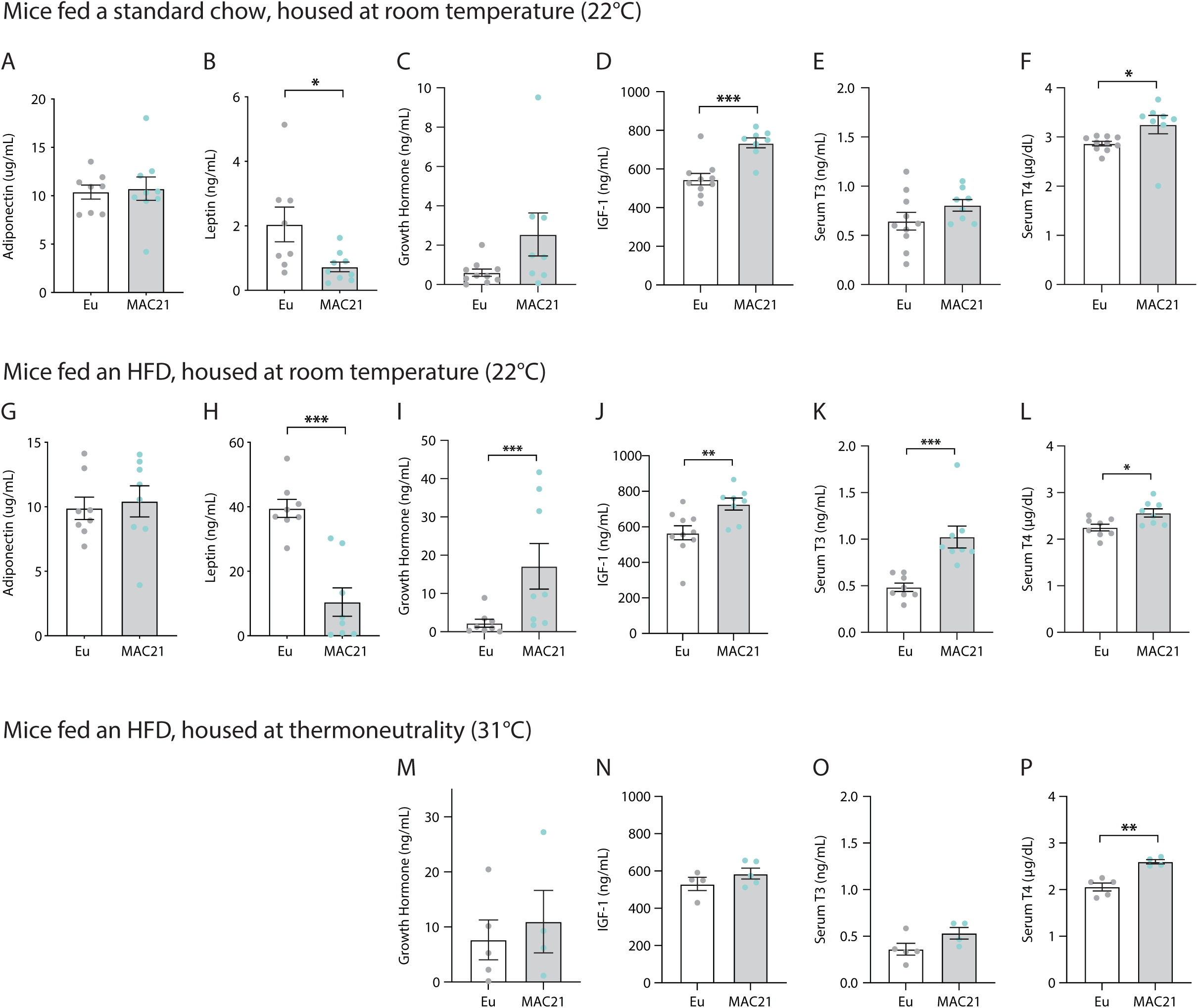
ELISA data from Euploid and MAC21 mice. Circulating levels of adiponectin (A), leptin (B), growth hormone (C), IGF-1 (D), thyroid hormone (T3, E), and thyroxine (T4, F) in chow-fed mice housed at room temperature (22°C). Circulating levels of adiponectin (G), leptin (H), growth hormone (I), IGF-1 (J), T3 (K), and T4 (L) in HFD-fed mice housed at 22°C. Circulating levels of growth hormone (M), IGF-1 (N), T3 (O), and T4 (P) in HFD-fed mice housed at thermoneutrality (30°C).

**Figure S2.**
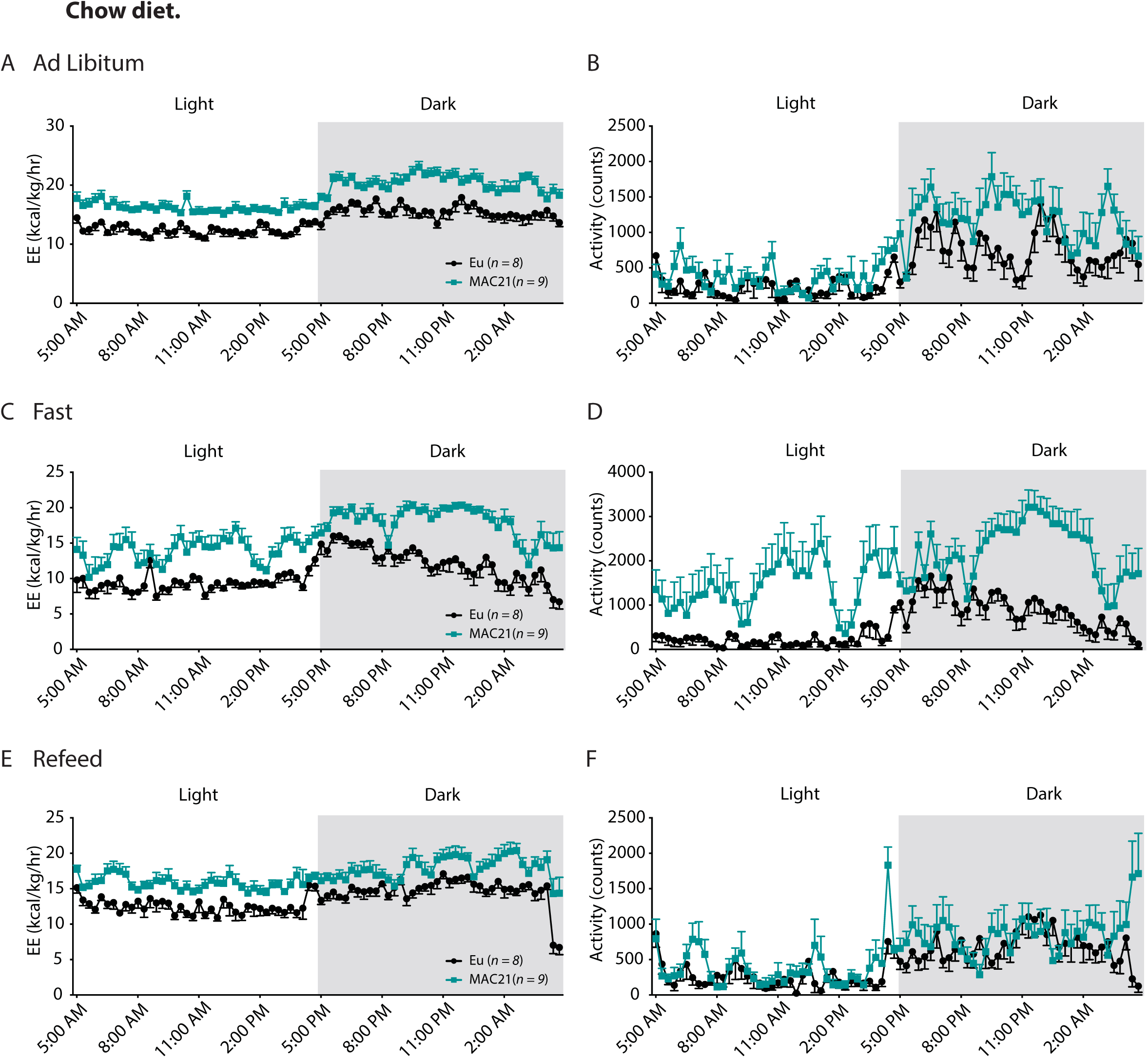
Indirect calorimetry of energy expenditure and physical activity in Euploid and MAC21 mice fed a standard chow. **A-B)** Energy expenditure (EE) and physical activity over a 24 h period in ad libitum chow-fed mice. **C-D)** Energy expenditure (EE) and physical activity over a 24 h period in fasted mice. **E-F)** Energy expenditure (EE) and physical activity over a 24 h period of refeeding after a fast.

**Figure S3.**
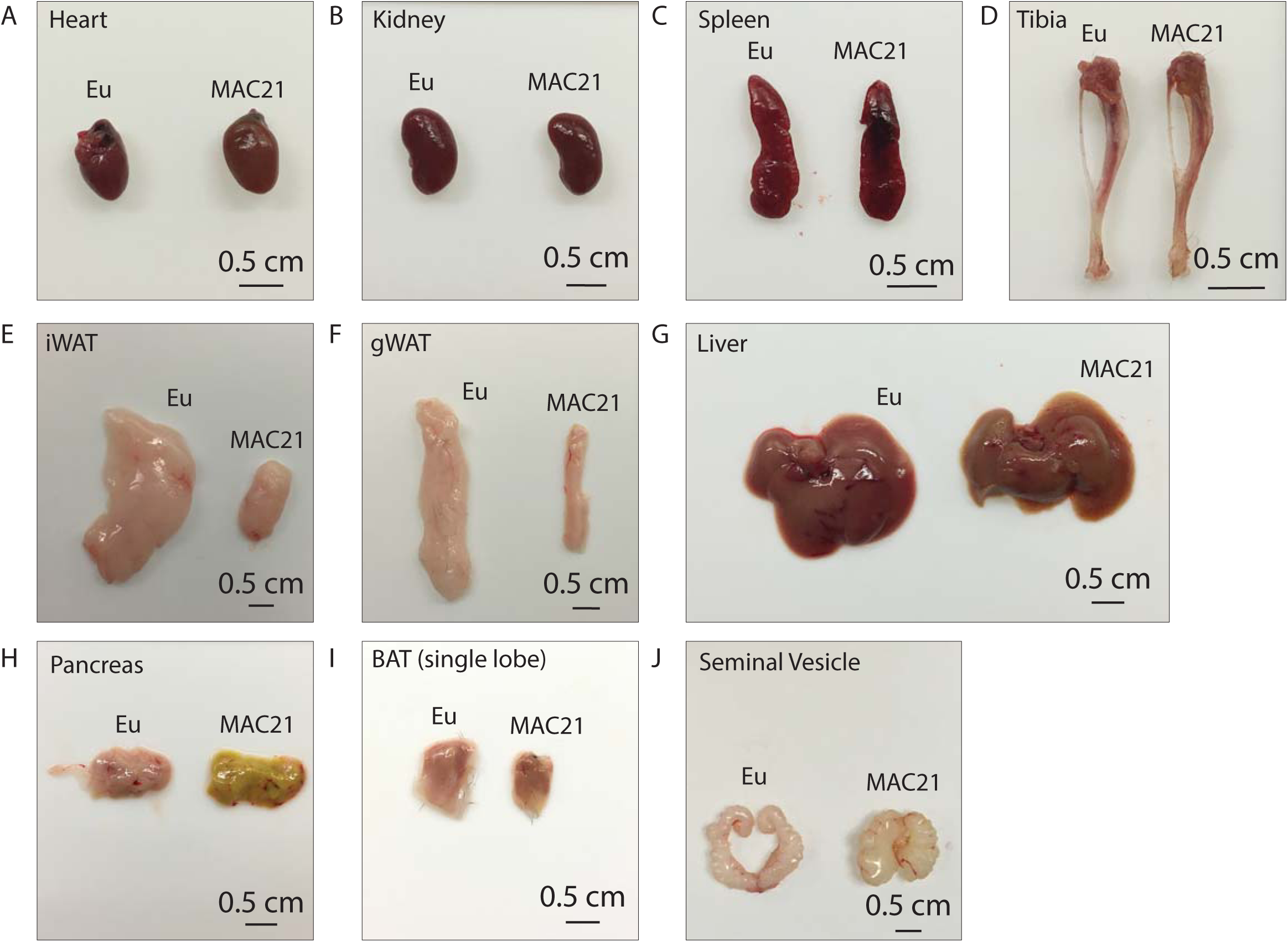
Representative tissue dissection images of Euploid and MAC21 mice fed a high-fat diet. Representative side-by-side dissection images of **A)** heart, **B)** kidney, **C)** spleen, **D)** tibia, **E)** inguinal white adipose tissue (iWAT), **F)** gonadal white adipose tissue (gWAT), **G)** liver, **H)** pancreas, **I)** brown adipose tissue (BAT), and **J)** seminal vesicle. Mice were housed at ambient room temperature (22°C).

**Figure S4.**
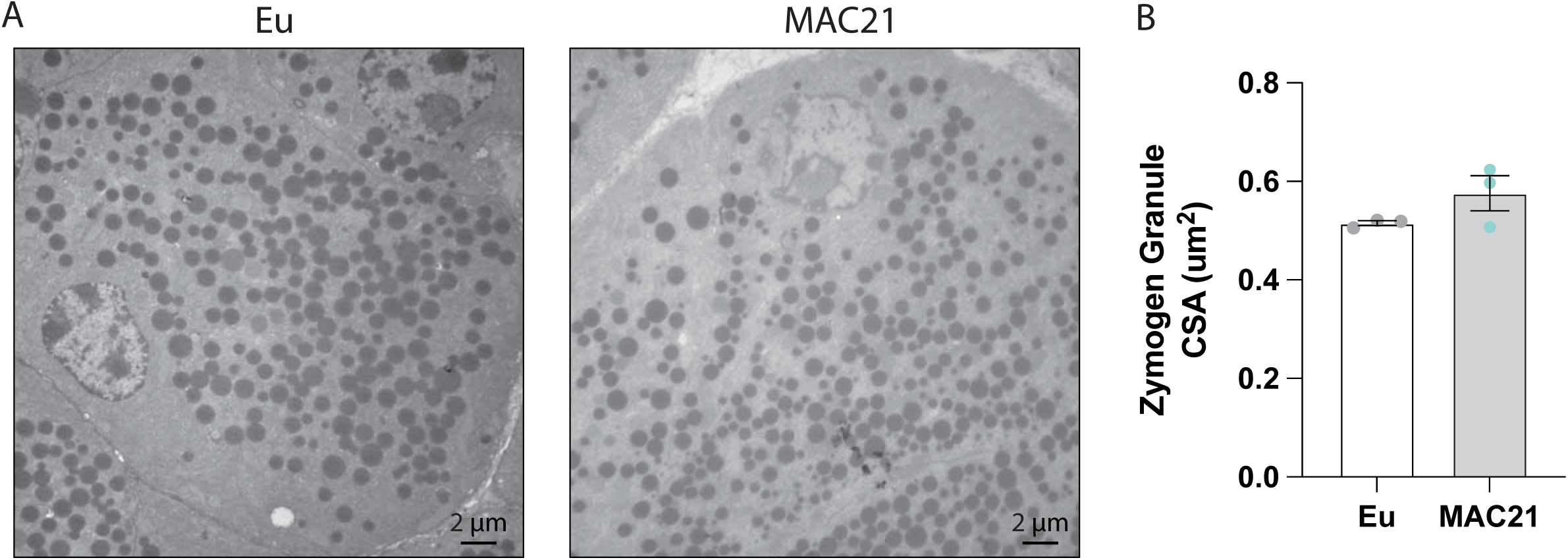
Electron micrograph quantification of pancreatic acinar cell zymogen granules. **A and B)** Representative Euploid and MAC21 acinar cells. **C)** Average zymogen granule cross-sectional area (CSA) quantification. Each data point represents a mouse average comprised of at least 3,600 zymogen granules from six unique locations within the pancreas. Analyses were performed on a total of 6 randomly selected HFD-fed mice (WT, *n* = 3; KO, *n* = 3).

**Figure S5.**
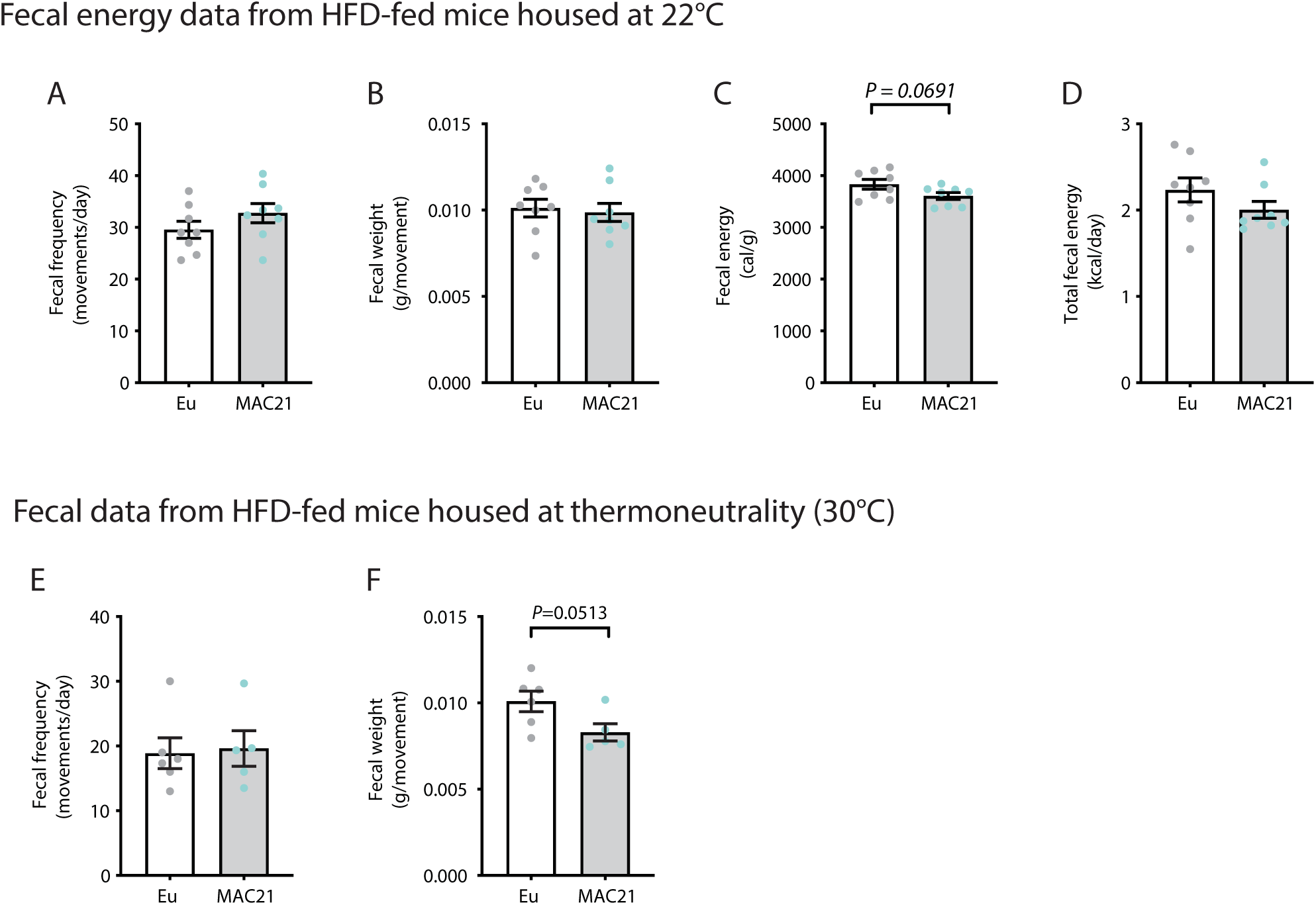
Fecal data from Euploid and MAC21 mice fed a high-fat diet at room temperature (22°C) or thermoneutrality (30°C). Fecal energy data from HFD-fed mice housed at room temperature (22°C) showing **A)** rate of fecal pellet production (bowel movement), **B)** Average fecal pellet weight, and **C-D)** Fecal energy composition as measure by fecal bomb calorimetry. Fecal data from mice fed a high-fat diet housed at thermoneutrality showing **E)** rate of fecal pellet production and **F)** average fecal weight per pellet.

**Figure S6.**
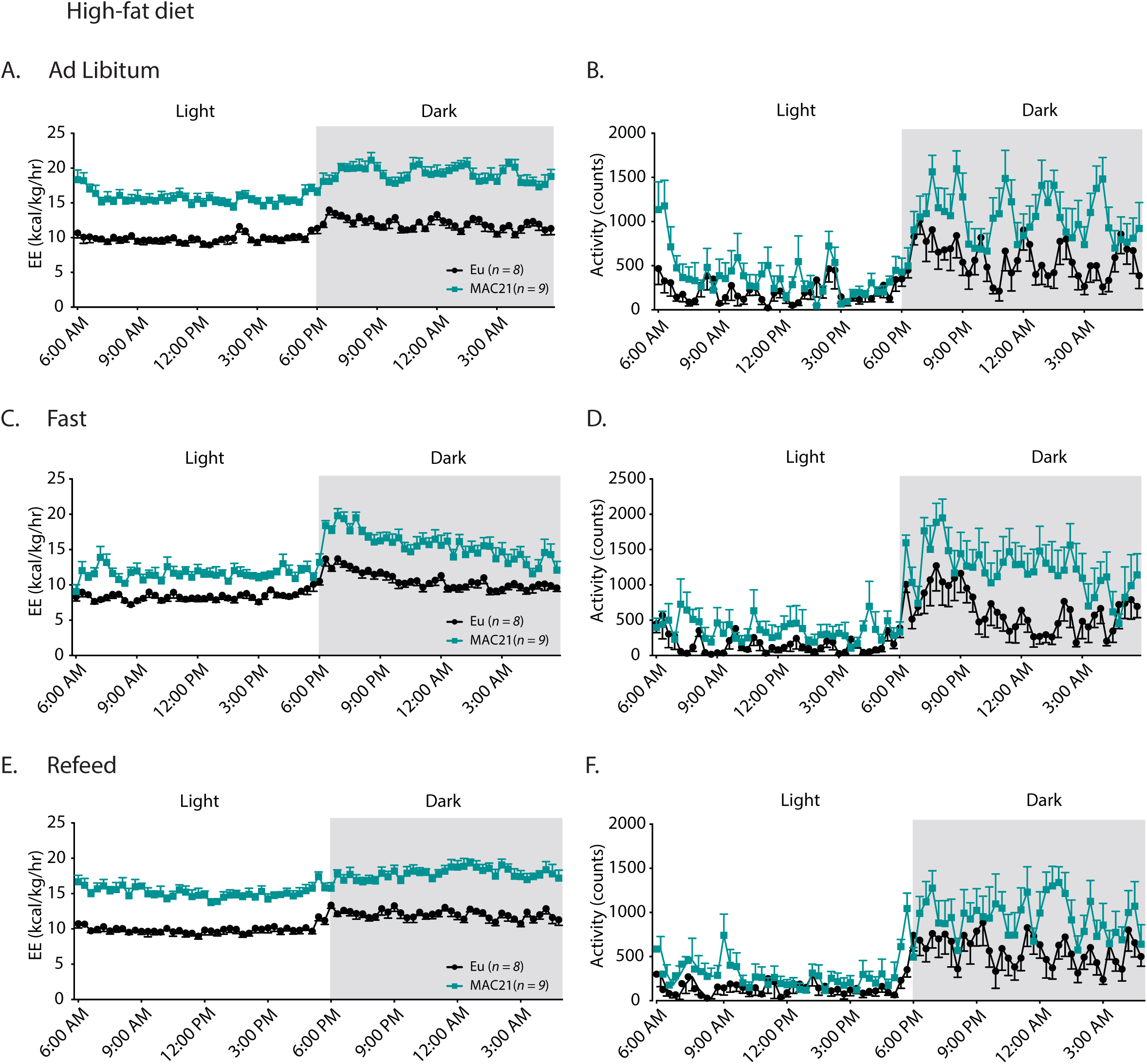
Indirect calorimetry of energy expenditure and physical activity in Euploid and MAC21 mice fed a high-fat diet. **A-B)** Energy expenditure (EE) and physical activity over a 24 h period in ad libitum HFD-fed mice. **C-D)** Energy expenditure (EE) and physical activity over a 24 h period in fasted mice. **E-F)** Energy expenditure (EE) and physical activity over a 24 h period of refeeding after a fast.

**Figure S7.**
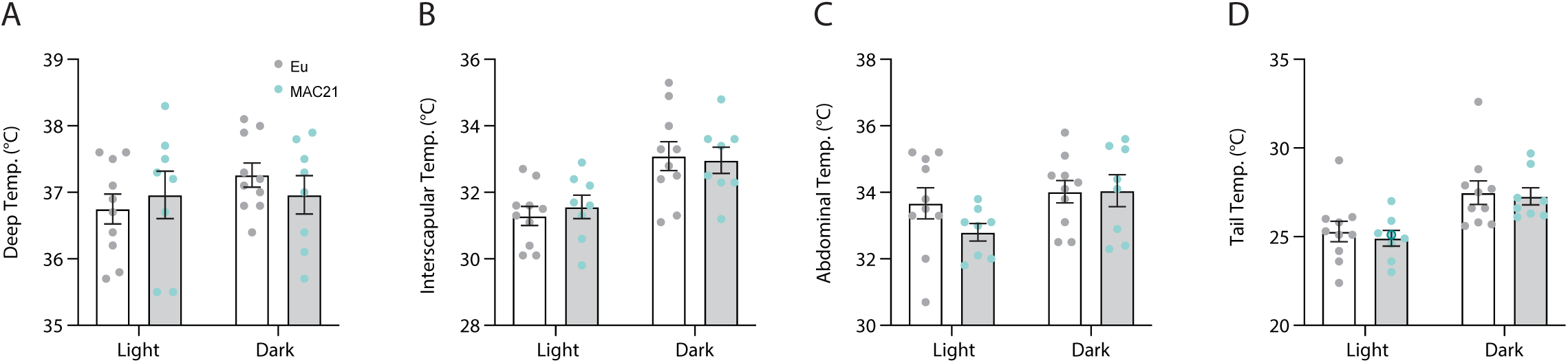
Light and dark cycle body temperature of Euploid and MAC21 mice fed a standard chow. **A)** Deep colonic temperature. **B)** Interscapular skin temperature. **C)** Abdominal Skin temperature. **D)** Tail skin temperature. Each data point represents the average of three days of independent data collection.

**Figure S8.**
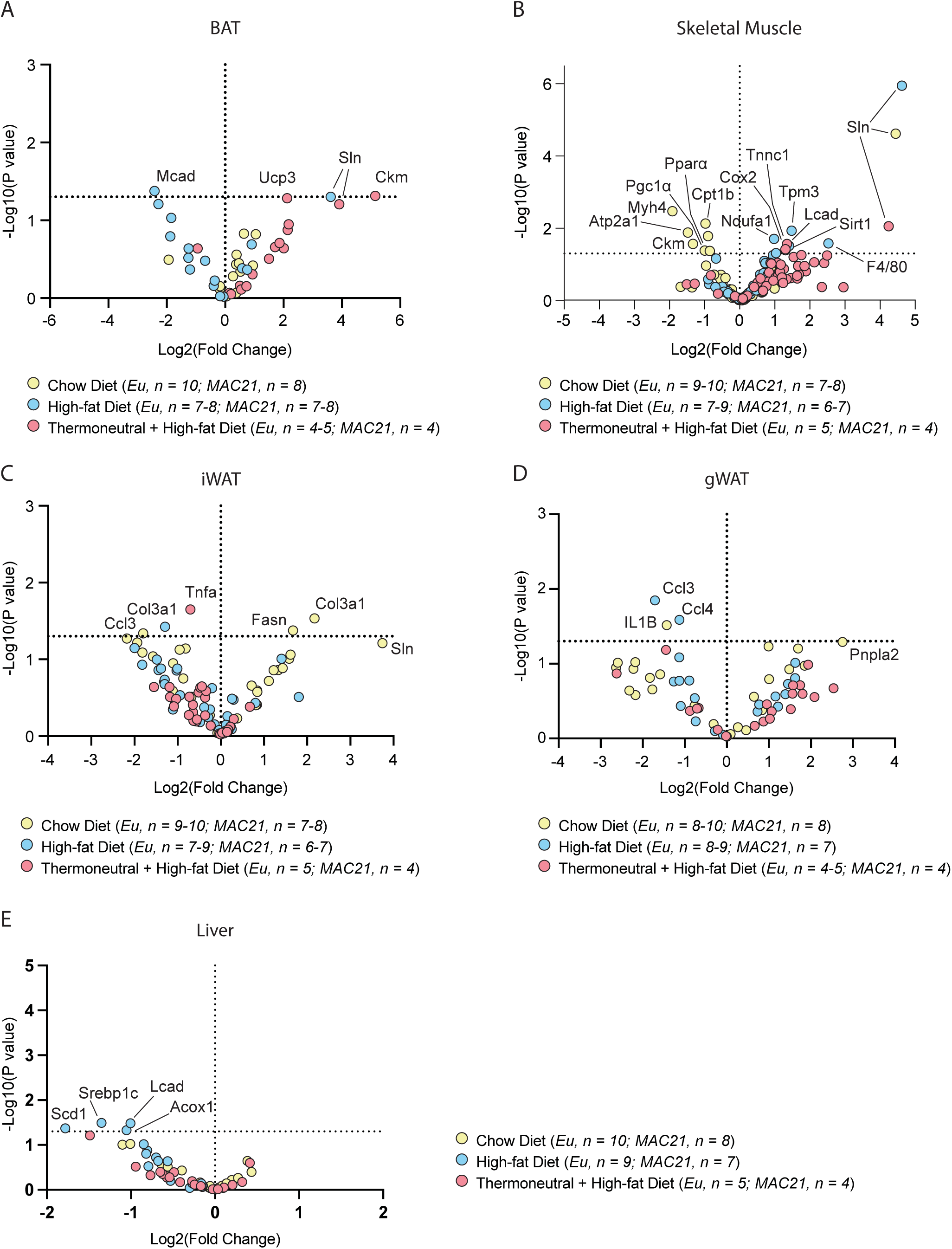
Volcano plot of all qPCR results from BAT, skeletal muscle, and white adipose tissue. Overlaid results of Euploid and MAC21 mice across all conditions assayed (chow diet at 22°C, high-fat diet at 22°C, and high-fat diet at 30°C) for **A)** Brown adipose tissue (BAT), **B)** Gastrocnemius (skeletal muscle), **C)** inguinal white adipose tissue (iWAT), **D)** gonadal white adipose tissue (gWAT), and **E)** Liver. Significant and/or relevant genes are labeled. Thermoneutral = 30°C housing condition.

**Figure S9.**
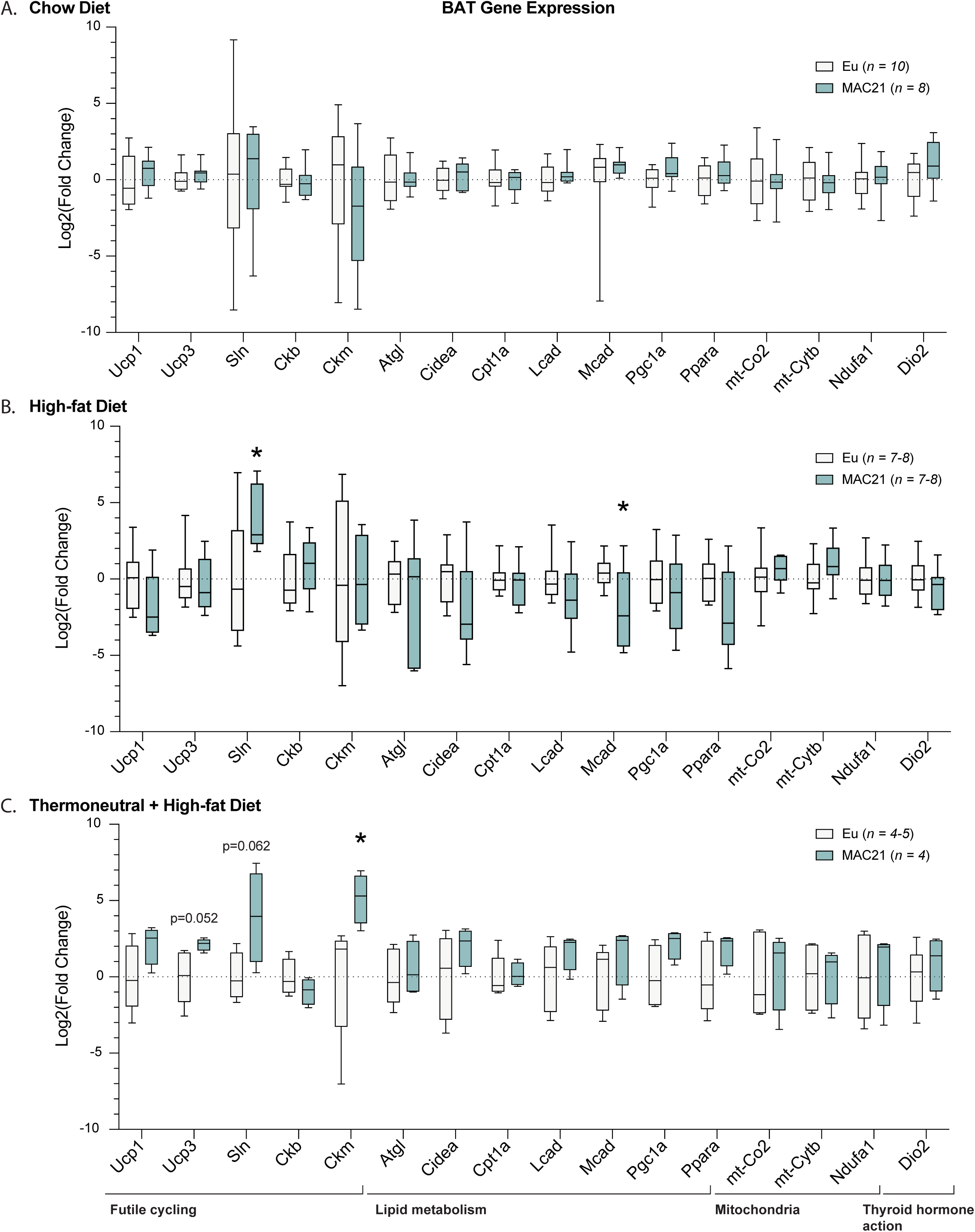
Gene expression analysis in brown adipose tissue. qPCR analysis of brown adipose tissue genes important for futile cycling, lipid metabolism, mitochondrial function, and thyroid hormone action in Euploid and MAC21 male mice fed a standard chow and housed at 22°C (A), fed a high-fat diet and housed at 22°C (B), and fed a high-fat diet and housed at thermoneutral 30°C (C).

**Figure S10.**
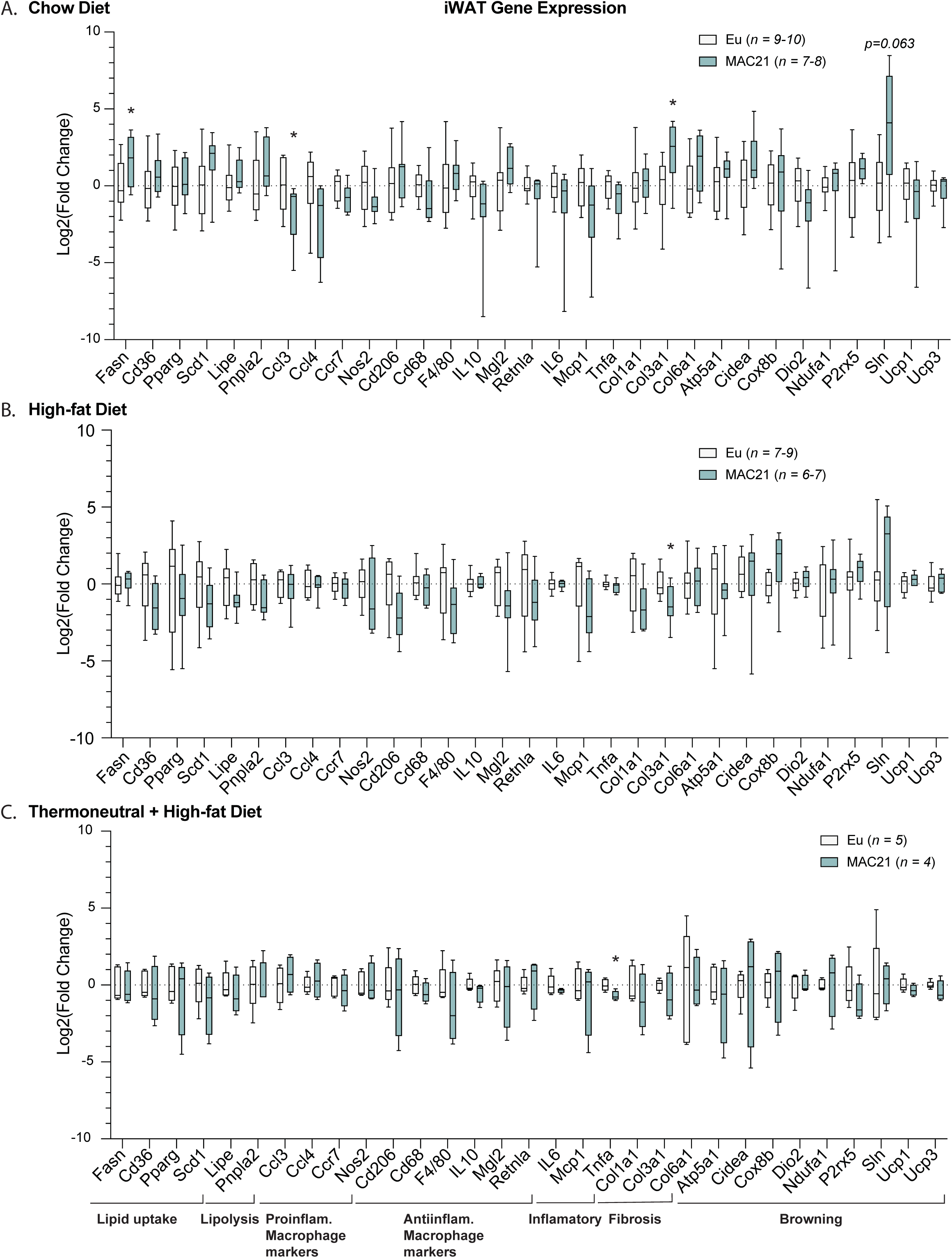
Gene expression analysis in inguinal white adipose tissue. qPCR analysis of inguinal white adipose tissue (iWAT) genes important for lipid uptake, lipolysis, inflammation, fibrosis, and browning in Euploid and MAC21 male mice fed a standard chow diet and housed at 22°C (A), fed a high-fat diet and housed at 22°C (B), and fed a high-fat diet and housed at thermoneutral 30°C (C).

**Figure S11.**
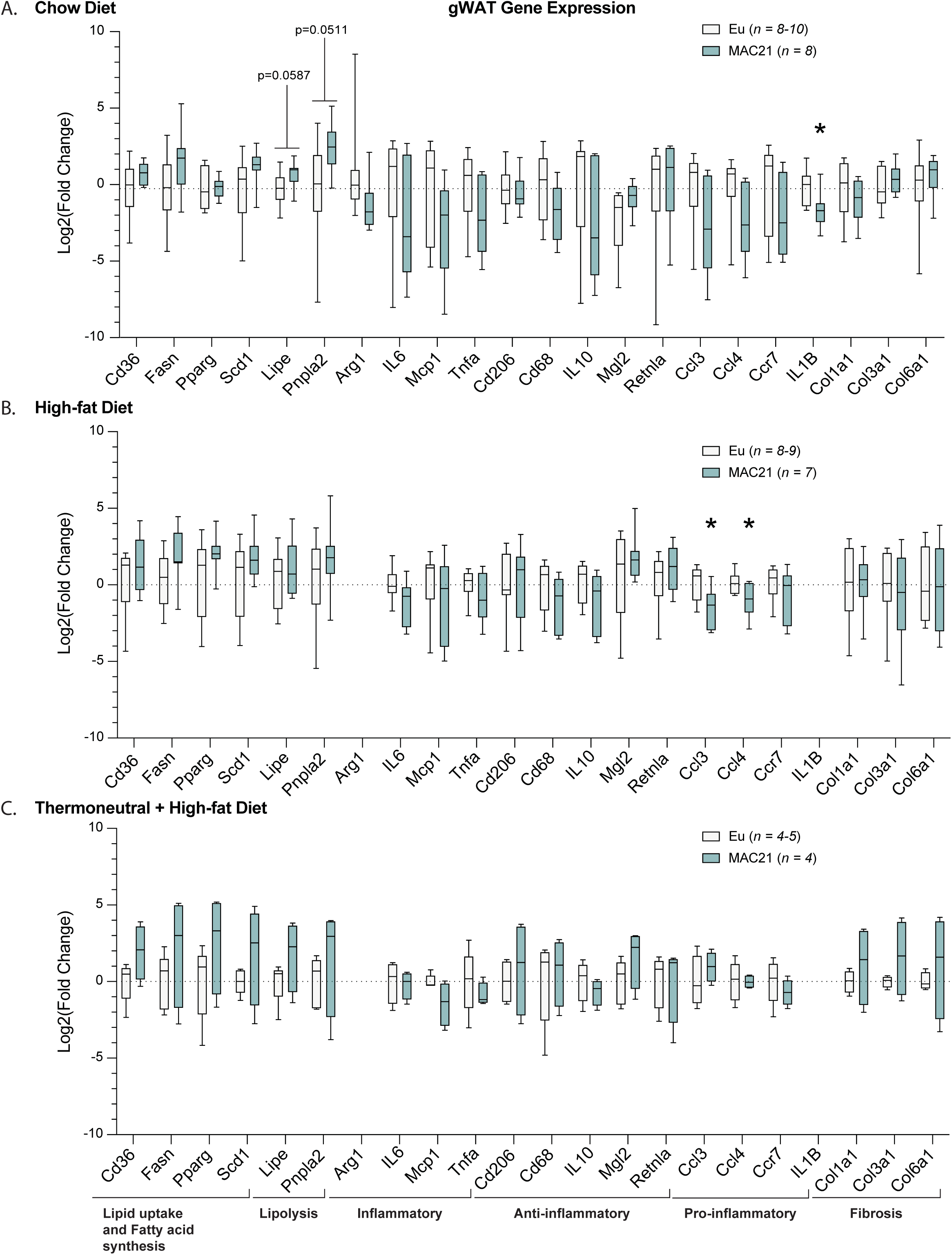
Gene expression analysis in gonadal white adipose tissue. qPCR analysis of gonadal white adipose tissue (gWAT) genes important for lipid uptake, fatty acid synthesis, lipolysis, inflammation, and fibrosis in Euploid and MAC21 male mice fed a standard chow diet and housed at 22°C (A), fed a high-fat diet and housed at 22°C (B), and fed a high-fat diet and housed at thermoneutral 30°C (C).

**Figure S12.**
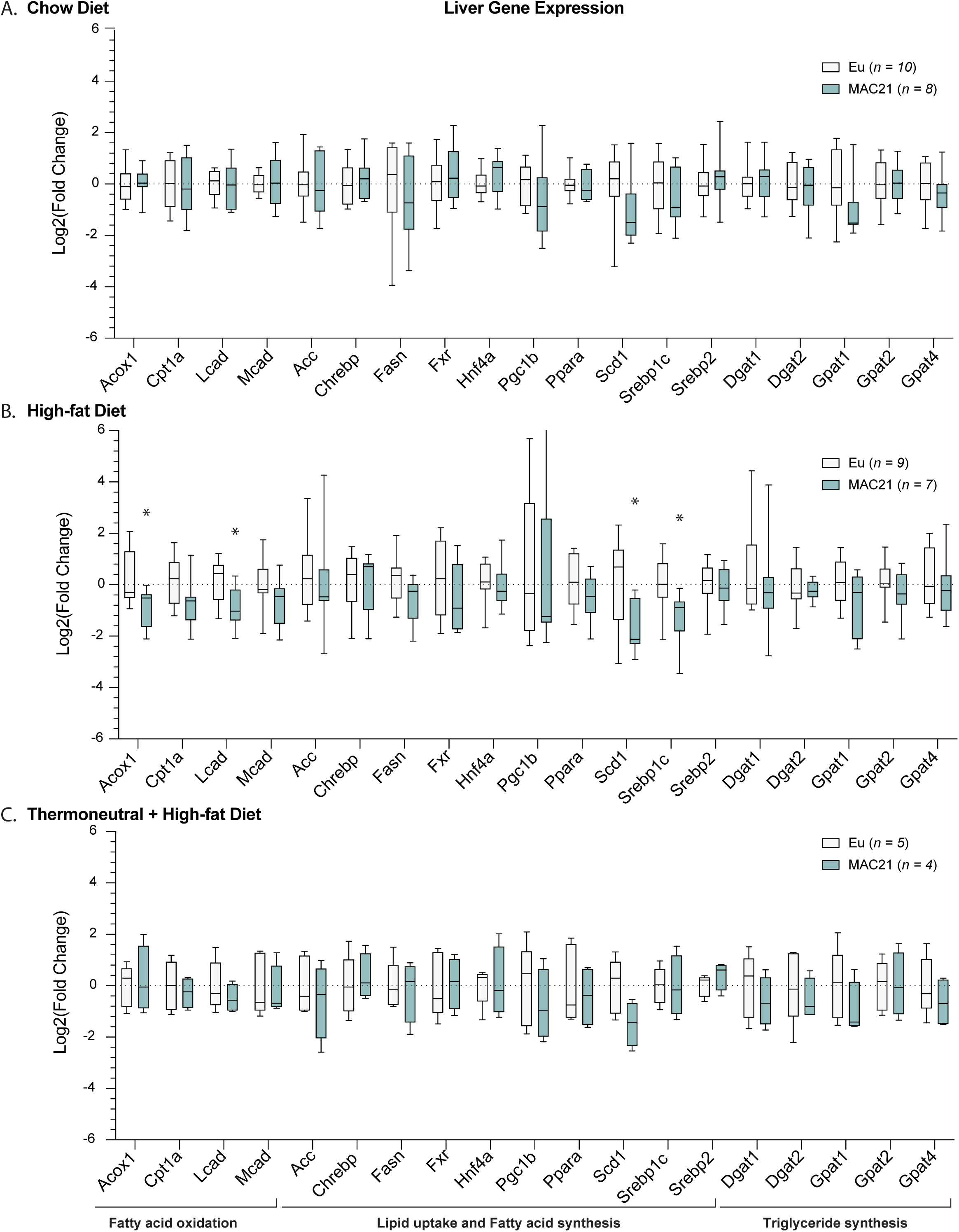
Gene expression analysis in liver. qPCR analysis of liver genes important for lipid uptake, fatty acid synthesis and oxidation, and triglyceride synthesis in Euploid and MAC21 male mice fed a standard chow and housed at 22°C (A), fed a high-fat diet and housed at 22°C (B), and fed a high-fat diet and housed at thermoneutral 30°C (C).

**Figure S13.**
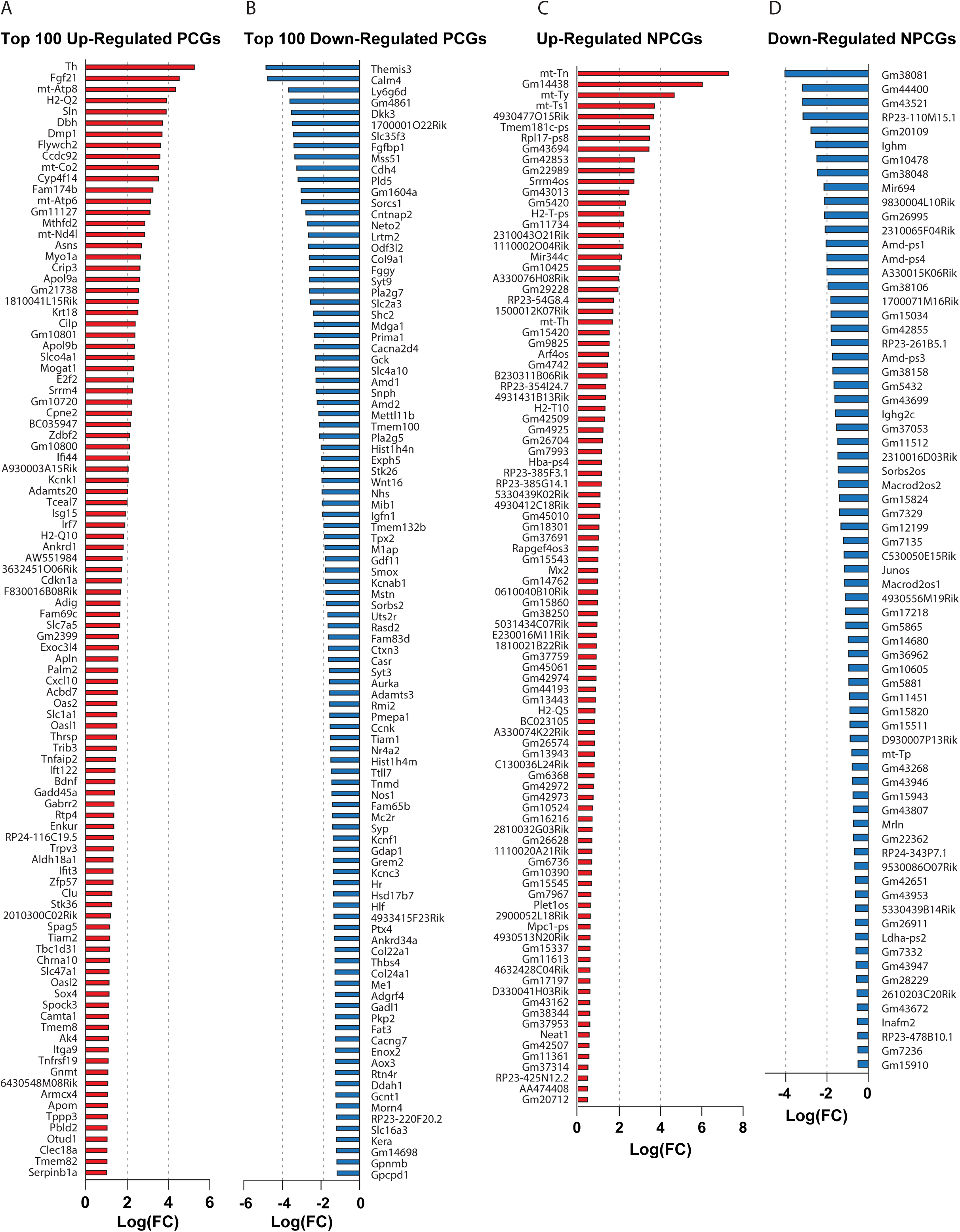
The top up- and down-regulated genes in skeletal muscle based on RNA-seq data. **A)** Top 100 up-regulated protein-coding genes. **B)** Top 100 down-regulated protein-coding genes. **C)** All up-regulated non-protein-coding genes. **D)** All down-regulated non-protein-coding genes. *n* = 5 Euploid and 4 MAC21 for RNA-sequencing experiments. All mice were fed a high-fat diet and housed at ambient room temperature (22°C).

**Figure S14.**
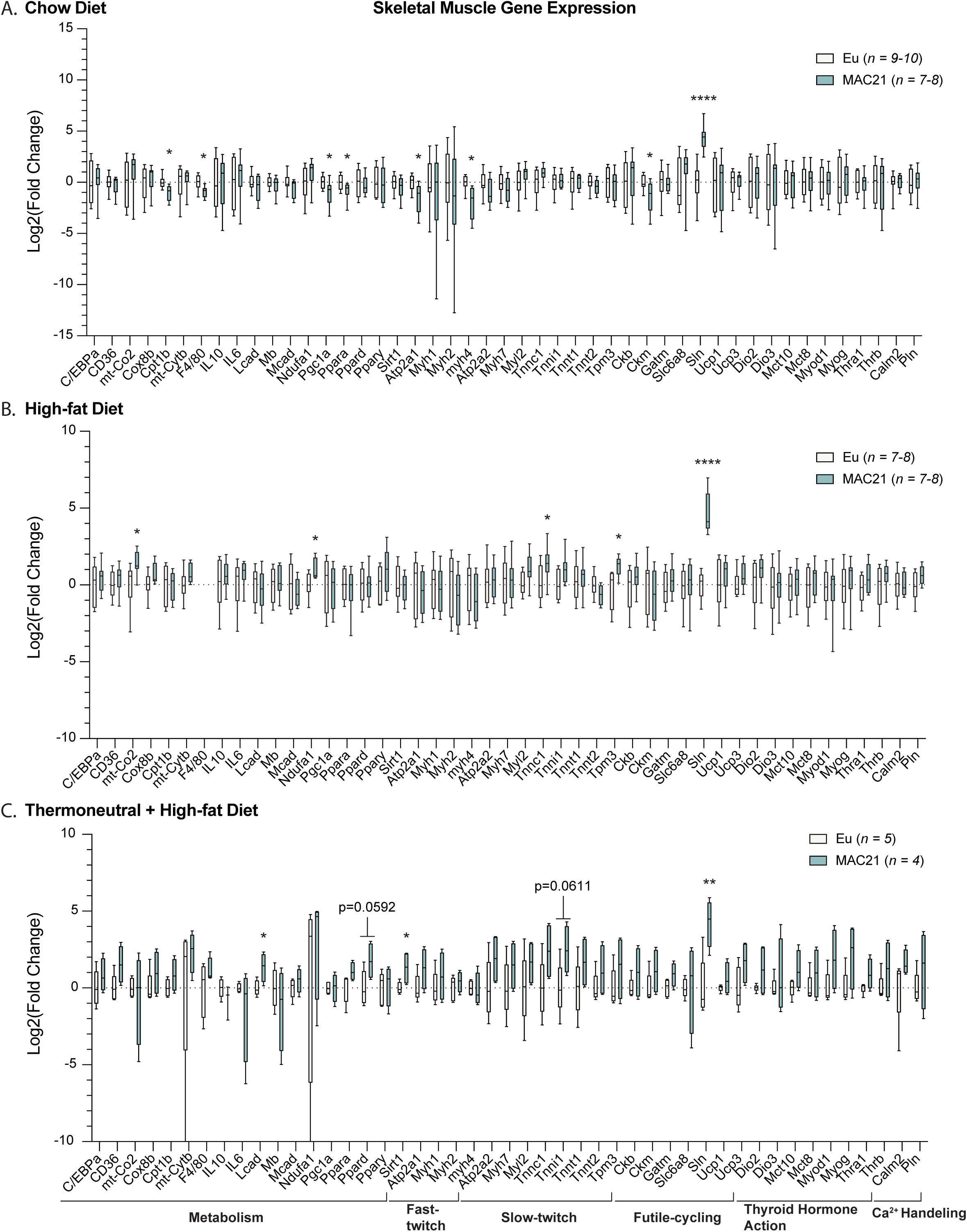
Gene expression analysis in skeletal muscle. qPCR analysis of gastrocnemius skeletal muscle genes important for general metabolism, fast- and slow-twitch fiber types, futile cycling, thyroid hormone action, and calcium handling in Euploid and MAC21 male mice fed a standard chow diet and housed at 22°C (A), fed a high-fat diet and housed at 22°C (B), and fed a high-fat diet and housed at thermoneutral 30°C (C).

**Figure S15.**
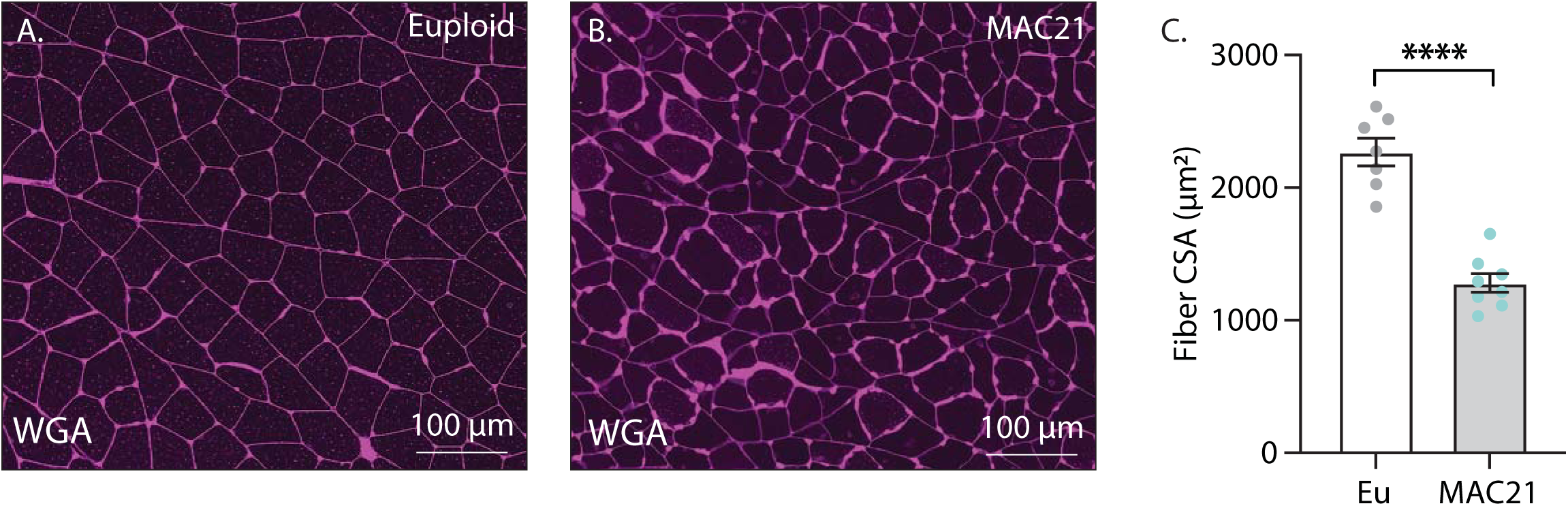
Skeletal muscle cross-sectional area analysis of MAC21 and Euploid mice fed a high-fat diet. **A-B)** Representative wheat-germ agglutinin (WGA) stained gastrocnemius skeletal muscle samples**. C)** Muscle fiber cross-sectional area (CSA) analysis. Mice were housed at ambient room temperature (22°C).

**Figure S16.**
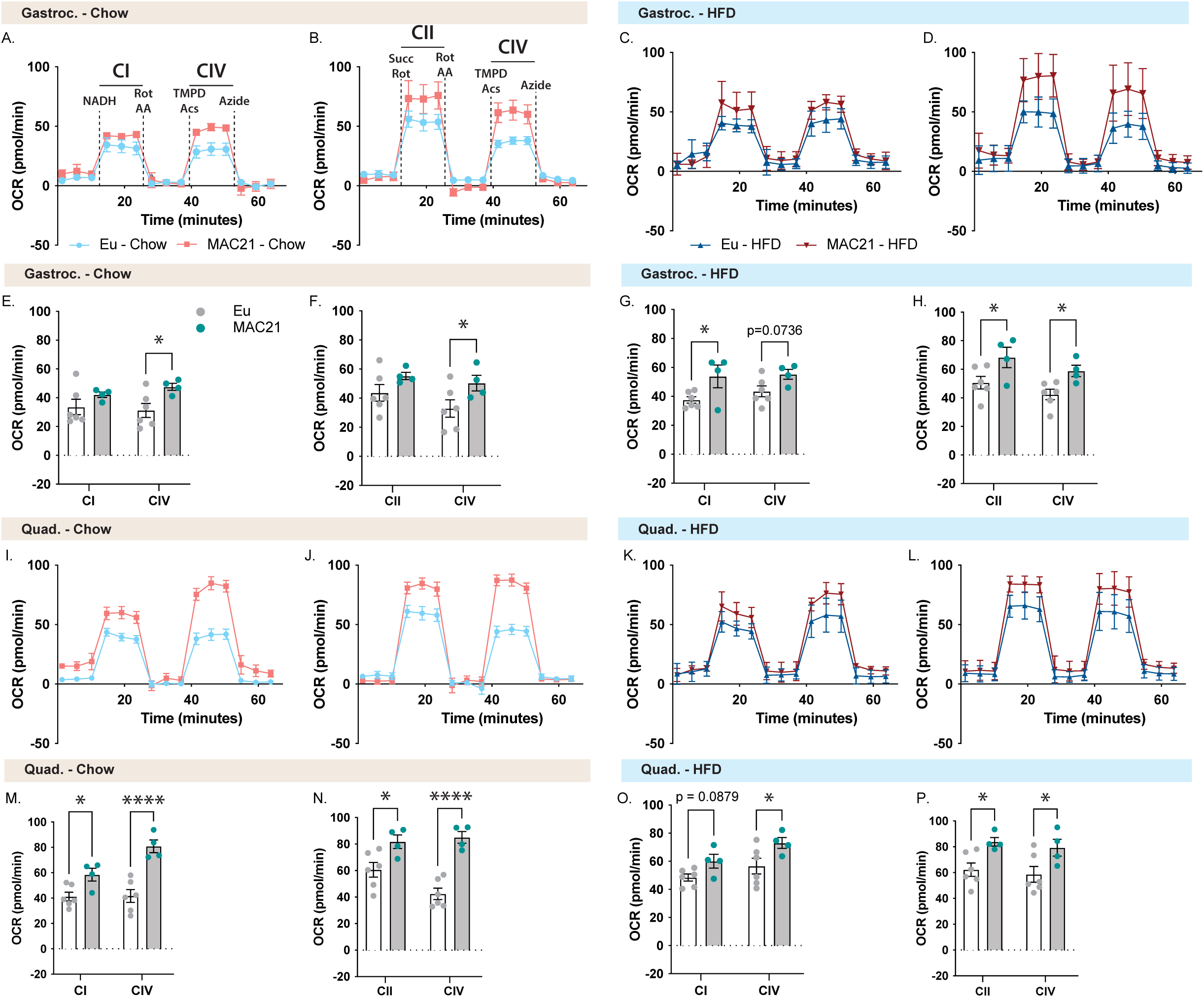
Mitochondrial respirometry analysis of gastrocnemius and quadricep muscle. **A and C)** Average group gastrocnemius (Gastroc) oxygen consumption rate (OCR) traces using NADH as a substrate for Euploid (Eu) and MAC21 male mice fed a standard chow or high-fat diet (HFD), respectively. **B and D)** Average group Gastroc OCR traces using succinate as a substrate in the presence of rotenone (Rot) for Euploid and MAC21 mice fed a standard chow or HFD, respectively. **E and G)** Mitochondrial complex I, and IV OCR quantification for A and C, respectively. **F and H)** Mitochondrial complex II and IV OCR quantification for B and D, respectively. **I and K)** Average group quadricep (Quad) OCR traces using NADH as a substrate for Euploid and MAC21 mice fed a standard chow or HFD, respectively. **J and L)** Average group Quad OCR traces using succinate as a substrate in the presence of rotenone (Rot) for Euploid and MAC21 mice fed a standard chow or HFD, respectively. **M and O)** Mitochondrial complex I and IV OCR quantification for I and K, respectively. **N and P)** Mitochondrial complex II and IV OCR quantification for J and L, respectively. All mice used for respirometry were house at ambient room temperature (22°C). AA, antimycin A; TMPD, N,N,N’,N’-tetramethyl-p-phenylenediamine; Acs, ascorbate; CI, mitochondrial complex I; CII, mitochondrial complex II; CIV, mitochondrial complex IV.

**Figure S17.**
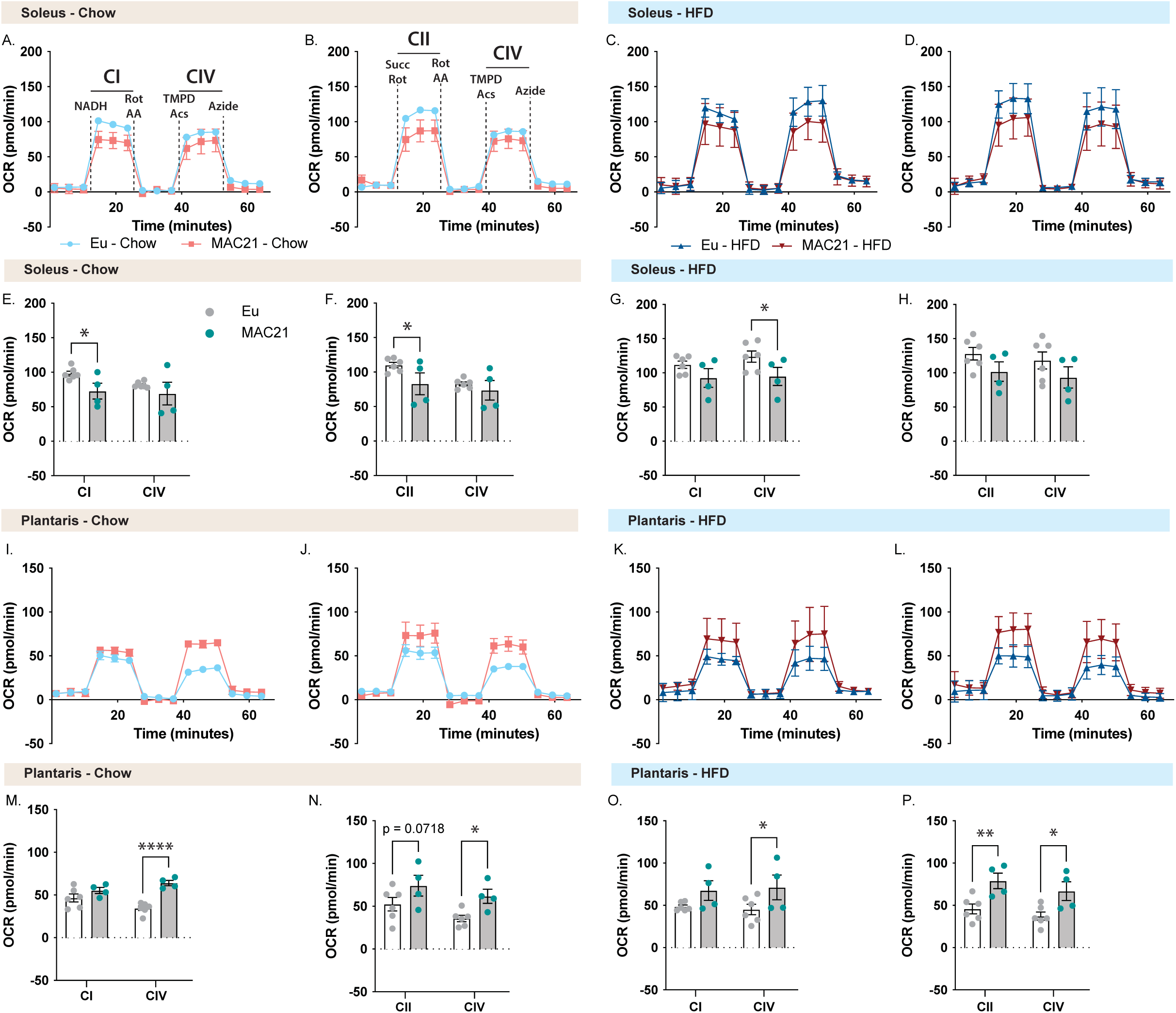
Mitochondrial respirometry analysis of soleus and plantaris muscle. **A and C)** Average group soleus oxygen consumption rate (OCR) traces using NADH as a substrate for Euploid (Eu) and MAC21 male mice fed a standard chow or high-fat diet (HFD), respectively. **B and D)** Average group soleus OCR traces using succinate as a substrate in the presence of Rotenone (Rot) for Euploid and MAC21 mice fed a standard chow or HFD, respectively. **E and G)** Mitochondrial complex I and IV OCR quantification for A and C, respectively. **F and H)** Mitochondrial complex II and IV OCR quantification for B and D, respectively. **I and K)** Average group plantaris OCR traces using NADH as a substrate for Euploid and MAC21 mice fed a standard chow or HFD, respectively. **J and L)** Average group plantaris OCR traces using succinate as a substrate in the presence of rotenone (Rot) for Euploid and MAC21 mice fed a standard chow or HFD, respectively. **M and O)** Mitochondrial complex I and IV OCR quantification for I and K, respectively. **N and P)** Mitochondrial complex II and IV OCR quantification for J and L, respectively. All mice used for respirometry were house at ambient room temperature (22°C). AA, antimycin A; TMPD, N,N,N’,N’-tetramethyl-p-phenylenediamine; Acs, ascorbate; CI, mitochondrial complex I; CII, mitochondrial complex II; CIV, mitochondrial complex IV.

**Figure S18.**
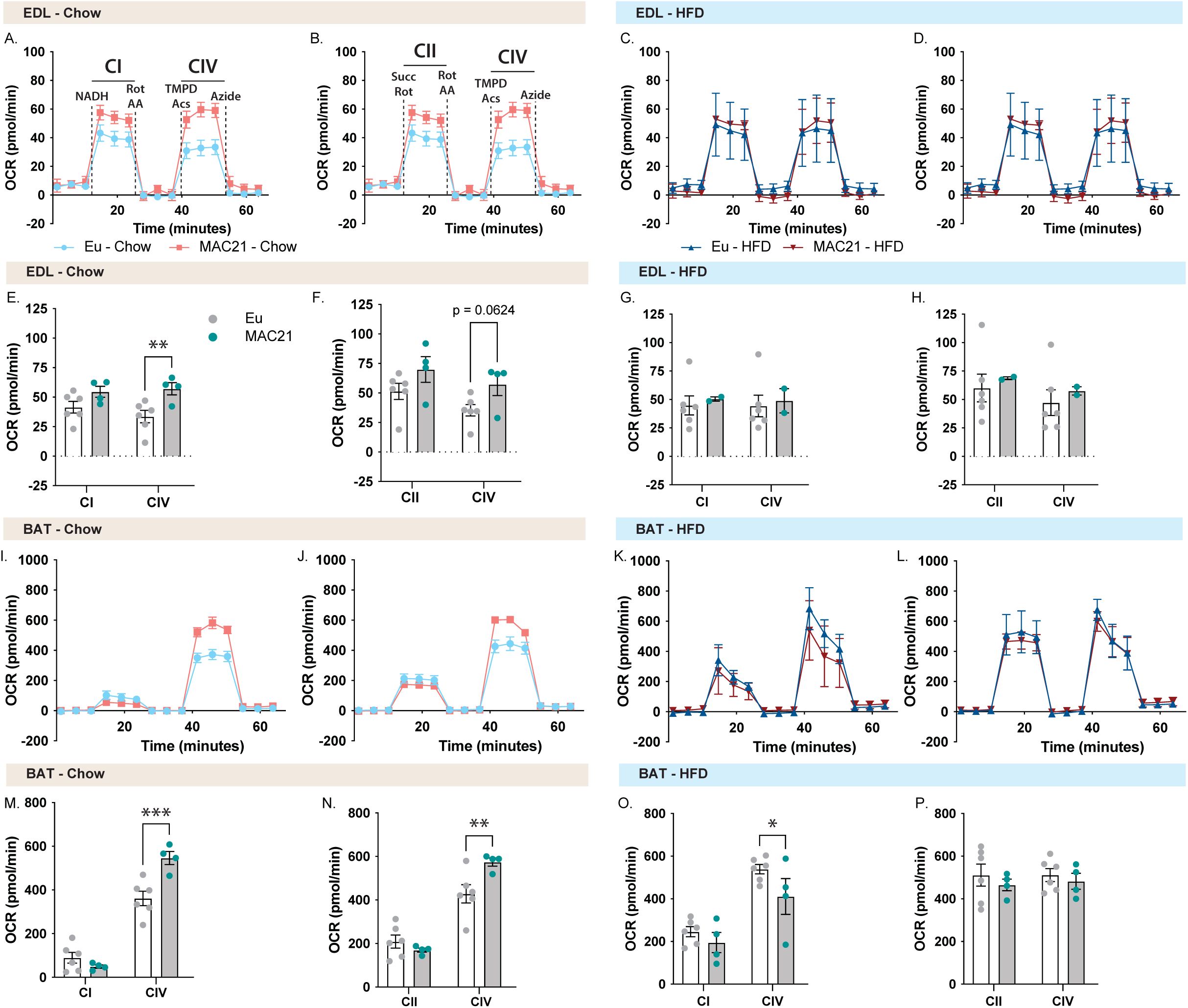
Mitochondrial respirometry analysis of EDL and BAT. **A and C)** Average group extensor digitorum longus (EDL) oxygen consumption rate (OCR) traces using NADH as a substrate for Euploid (Eu) and MAC21 male mice fed a standard chow or high-fat diet (HFD), respectively. **B and D)** Average group EDL OCR traces using succinate as a substrate in the presence of rotenone (Rot) for Euploid and MAC21 mice fed a standard chow or HFD, respectively. **E and G)** Mitochondrial complex I and IV OCR quantification for A and C, respectively. **F and H)** Mitochondrial complex II and IV OCR quantification for B and D, respectively. **I and K)** Average group brown adipose tissue (BAT) OCR traces using NADH as a substrate for Euploid and MAC21 mice fed a standard chow or HFD, respectively. **J and L)** Average group brown adipose tissue (BAT) OCR traces using succinate as a substrate in the presence of rotenone (Rot) for Euploid and MAC21 mice fed a standard chow or HFD, respectively. **M and O)** Mitochondrial complex I and IV OCR quantification for I and K, respectively. **N and P)** Mitochondrial complex II and IV OCR quantification for J and L, respectively. All mice used for respirometry were house at ambient room temperature (22°C). AA, antimycin A; TMPD, N,N,N’,N’-tetramethyl-p-phenylenediamine; Acs, ascorbate; CI, mitochondrial complex I; CII, mitochondrial complex II; CIV, mitochondrial complex IV.

**Figure S19.**
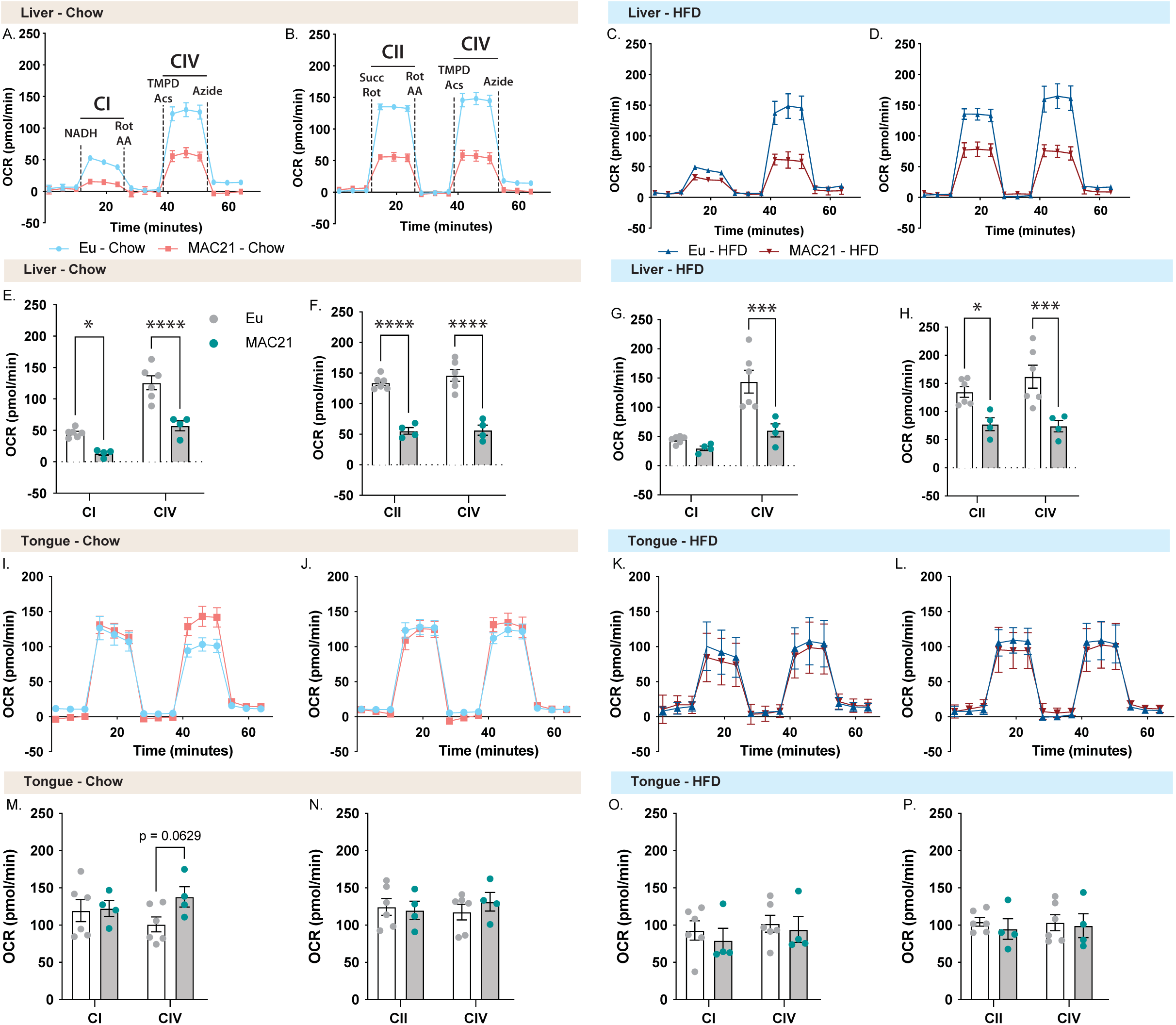
Mitochondrial respirometry analysis of liver and tongue. **A and C)** Average group liver oxygen consumption rate (OCR) traces using NADH as a substrate for Euploid (Eu) and MAC21 male mice fed a standard chow or high-fat diet (HFD), respectively. **B and D)** Average group liver OCR traces using succinate as a substrate in the presence of rotenone (Rot) for Euploid and MAC21 mice fed a standard chow or HFD, respectively. **E and G)** Mitochondrial complex I and IV OCR quantification for A and C, respectively. **F and H)** Mitochondrial complex II and IV OCR quantification for B and D, respectively. **I and K)** Average group tongue OCR traces using NADH as a substrate for Euploid and MAC21 mice fed a standard chow or HFD, respectively. **J and L)** Average group tongue OCR traces using succinate as a substrate in the presence of rotenone (Rot) for Euploid and MAC21 mice fed a standard chow or HFD, respectively. **M and O)** Mitochondrial complex I and IV OCR quantification for I and K, respectively. **N and P)** Mitochondrial complex II and IV OCR quantification for J and L, respectively. All mice used for respirometry were house at ambient room temperature (22°C). AA, antimycin A; TMPD, N,N,N’,N’-tetramethyl-p-phenylenediamine; Acs, ascorbate; CI, mitochondrial complex I; CII, mitochondrial complex II; CIV, mitochondrial complex IV.

**Figure S20.**
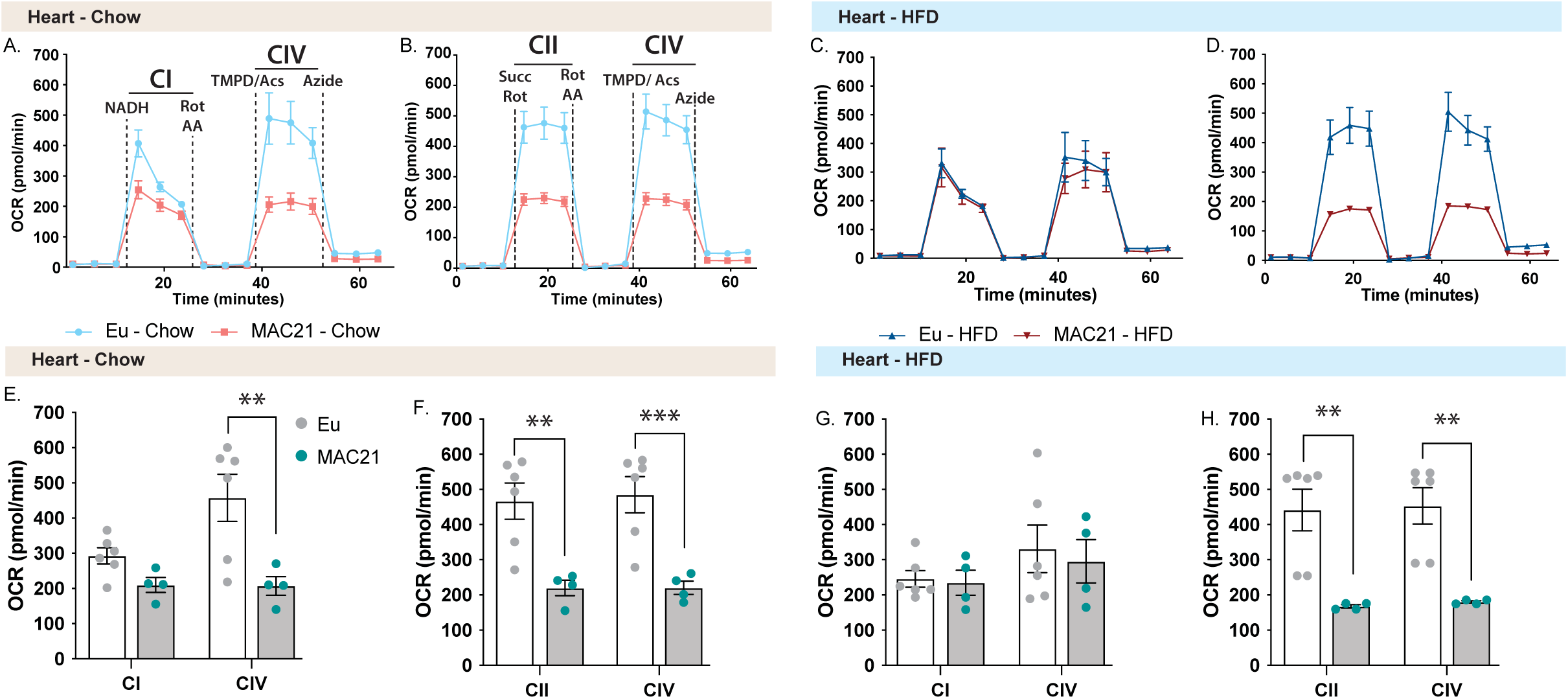
Mitochondrial respirometry analysis of heart. **A and C)** Average group heart oxygen consumption rate (OCR) traces using NADH as a substrate for Euploid (Eu) and MAC21 male mice fed a standard chow or high-fat diet (HFD), respectively. **B and D)** Average group heart OCR traces using succinate as a substrate in the presence of rotenone (Rot) for Euploid and MAC21 mice fed a standard chow or HFD, respectively. **E and G)** Mitochondrial complex I and IV OCR quantification for A and C, respectively. **F and H)** Mitochondrial complex II and IV OCR quantification for B and D, respectively. All mice used for respirometry were house at ambient room temperature (22°C). AA, antimycin A; TMPD, N,N,N’,N’-tetramethyl-p-phenylenediamine; Acs, ascorbate; CI, mitochondrial complex I; CII, mitochondrial complex II; CIV, mitochondrial complex IV.

**Figure S21.**
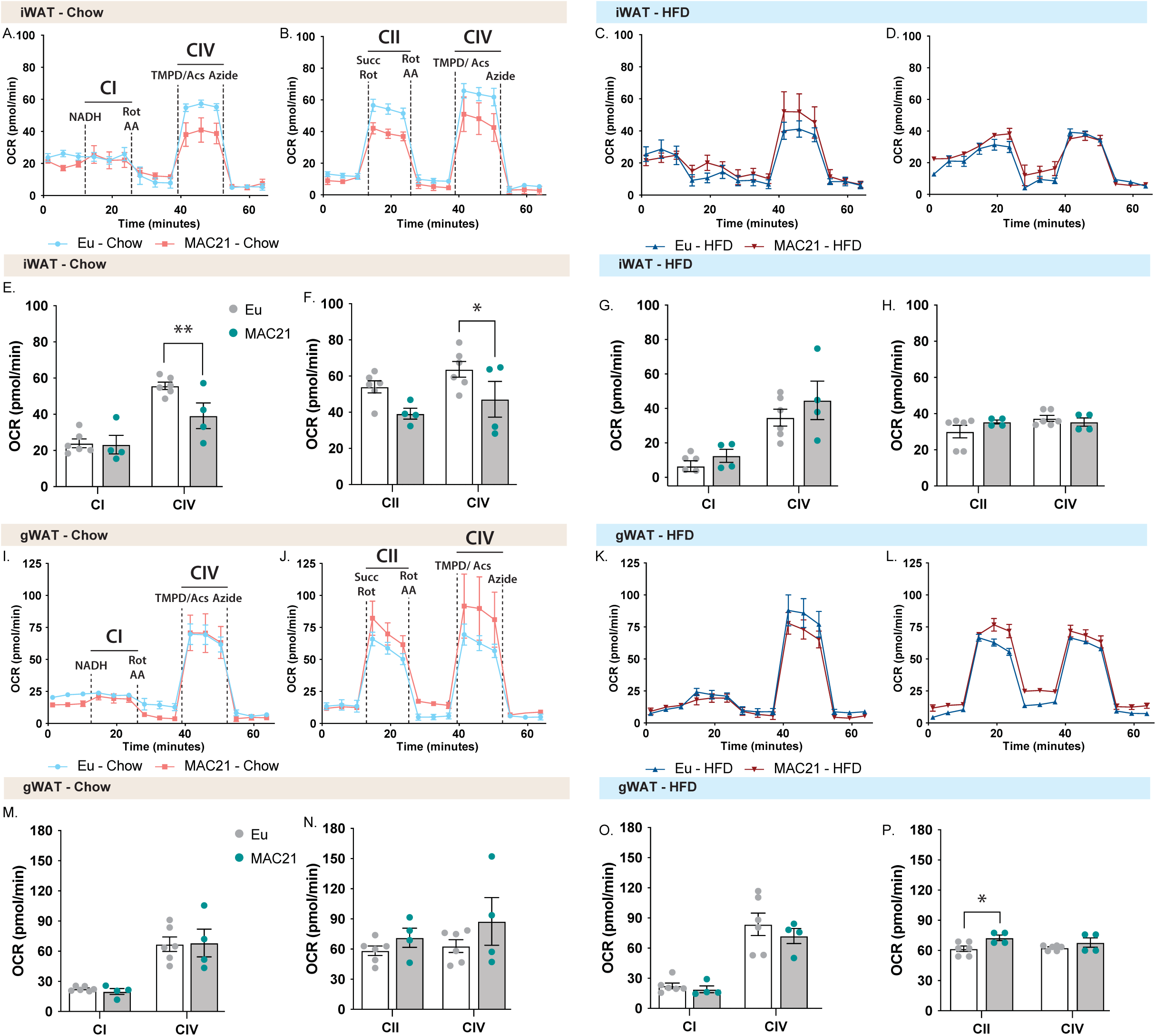
Mitochondrial respirometry analysis of inguinal (iWAT) and gonadal (gWAT) white adipose tissue. **A and C)** Average group iWAT oxygen consumption rate (OCR) traces using NADH as a substrate for Euploid (Eu) and MAC21 male mice fed a standard chow or high-fat diet (HFD), respectively. **B and D)** Average group liver OCR traces using succinate as a substrate in the presence of rotenone (Rot) for Euploid and MAC21 mice fed a standard chow or HFD, respectively. **E and G)** Mitochondrial complex I and IV OCR quantification for A and C, respectively. **F and H)** Mitochondrial complex II and IV OCR quantification for B and D, respectively. **I and K)** Average group gWAT OCR traces using NADH as a substrate for Euploid and MAC21 mice fed a standard chow or HFD, respectively. **J and L)** Average group gWAT OCR traces using succinate as a substrate in the presence of rotenone (Rot) for Euploid and MAC21 mice fed a standard chow or HFD, respectively. **M and O)** Mitochondrial complex I and IV OCR quantification for I and K, respectively. **N and P)** Mitochondrial complex II and IV OCR quantification for J and L, respectively. All mice used for respirometry were house at ambient room temperature (22°C). AA, antimycin A; TMPD, N,N,N’,N’-tetramethyl-p-phenylenediamine; Acs, ascorbate; CI, mitochondrial complex I; CII, mitochondrial complex II; CIV, mitochondrial complex IV.

**Figure S22.**
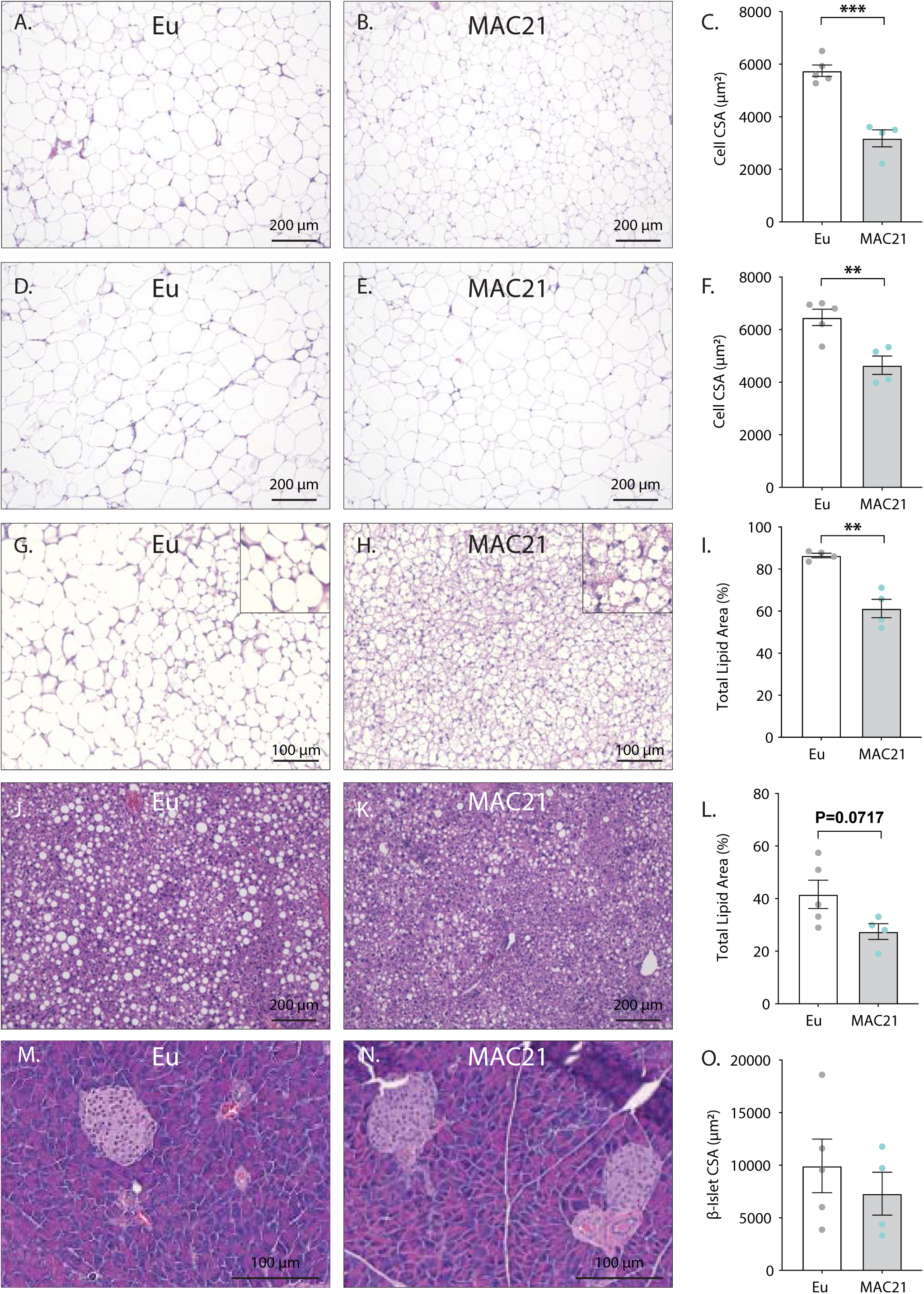
Histology of HFD-fed Euploid and MAC21 mice housed at thermoneutrality (30°C). Representative hematoxylin and eosin (H&E) stained sections of inguinal white adipose tissue (iWAT, A-B), gonadal white adipose tissue (gWAT, D and E), brown adipose tissue (BAT, H and I), liver (J and K), and pancreas (M and N). **C)** Average iWAT adipocyte cross-sectional area (CSA) quantification. **F)** Average gWAT adipocyte CSA quantification. **I)** Average BAT area covered by lipid droplets per focal plane. **L)** Average liver area covered by lipid droplets per focal plane. **O)** Average pancreas β-islet CSA quantification.

**Table S1.** Tissue weights of chow-fed Euploid and MAC21 male mice at termination of the study.

|  | Eu | MAC21 | *P* |
| --- | --- | --- | --- |
| *N* | 7 | 6 |  |
| BW (g) | **47.17 ± 3.73** | **26.77 ± 3.14** | **0.002** |
| gWAT ( g) | **1.47 ± 0.29** | **0.60 ± 0.19** | **0.040** |
| iWAT (g) | 1.15 ± 0.30 | 0.43 ± 0.20 | 0.080 |
| Liver (g) | **2.42 ± 0.31** | **1.54 ± 0.18** | **0.034** |
| Pancreas (g) | 0.30 ± 0.02 | 0.27 ± 0.01 | 0.109 |
| Kidney (g) | 0.19 ± 0.01 | 0.15 ± 0.02 | 0.051 |
| Heart (g) | **0.16 ± 0.01** | **0.14 ± 0.01** | **0.025** |

Eu: Euploid; BW: body weight; gWAT: gondal white adipose tissue; iWAT: inguinal white adipose tissue; BAT: brown adipose tissue

**Table S2.** Indirect calorimetry analysis of male (16.5 weeks of age) Euploid and MAC21 littermate mice fed a standard chow.

|  | Eu | MAC21 | P | Eu | MAC21 | P | Eu | MAC21 | P |
| --- | --- | --- | --- | --- | --- | --- | --- | --- | --- |
| **<u>Low-fat chow diet (Male)</u>** | **<u>Ad-libitum (dark cycle)</u>** |  |  | **<u>Ad-libitum (light cycle)</u>** |  |  | **<u>Ad-libitum (24 hr)</u>** |  |  |
| N | 8 | 9 |  | 8 | 9 |  | 8 | 9 |  |
| Body weight (g) |  |  |  |  |  |  | **<u>28.9 ± 1.09</u>** | **<u>20.6 ± 0.29</u>** | **<u><0.001</u>** |
| Food intake (kcal) | 9.76 ± 0.65 | 10.66 ± 0.58 | 0.320 | 2.44 ± 0.45 | 2.67 ± 0.20 | 0.639 | 12.2 ± 0.92 | 13.3 ± 0.56 | 0.302 |
| VO2 (mL/kg lean mass/h) | **<u>4381 ± 180</u>** | **<u>5674 ± 69</u>** | **<u><0.001</u>** | **<u>3589 ± 123</u>** | **<u>4616 ± 83</u>** | **<u><0.001</u>** | **<u>3984 ± 151</u>** | **<u>5145 ± 79</u>** | **<u><0.001</u>** |
| VCO2 (mL/kg lean mass/h) | **<u>4231 ± 182</u>** | **<u>5600 ± 97</u>** | **<u><0.001</u>** | **<u>3158 ± 122</u>** | **<u>4170 ± 109</u>** | **<u><0.001</u>** | **<u>3695 ± 150</u>** | **<u>4889 ± 81</u>** | **<u><0.001</u>** |
| RER (VCO2/VO2) | **<u>0.96 ± 0.006</u>** | **<u>0.99 ± 0.007</u>** | **<u>0.030</u>** | 0.88 ± 0.008 | 0.90 ± 0.010 | 0.099 | **<u>0.92 ± 0.006</u>** | **<u>0.94 ± 0.005</u>** | **<u>0.011</u>** |
| EE (kcal/kg lean mass/h) | **<u>21.9 ± 0.91</u>** | **<u>28.5 ± 0.49</u>** | **<u><0.001</u>** | **<u>17.6 ± 0.62</u>** | **<u>22.7 ± 0.45</u>** | **<u><0.001</u>** | **<u>19.8 ± 0.76</u>** | **<u>25.6 ± 0.40</u>** | **<u><0.001</u>** |
| Total activity (beam breaks) | **<u>30160 ± 2377</u>** | **<u>47055 ± 5532</u>** | **<u>0.017</u>** | **<u>9639 ± 1589</u>** | **<u>14530 ± 1107</u>** | **<u>0.021</u>** | **<u>39798 ± 3869</u>** | **<u>61586 ± 6366</u>** | **<u>0.013</u>** |
| Ambulatory activity (counts) | **<u>16577 ± 3138</u>** | **<u>28148 ± 4020</u>** | **<u>0.042</u>** | 4605 ± 1662 | 7135 ± 778 | 0.172 | 21181 ± 4736 | 35283 ± 4680 | 0.052 |
|  | **<u>Fasting (dark cycle)</u>** |  |  | **<u>Fasting (light cycle)</u>** |  |  | **<u>Fasting (24 hr)</u>** |  |  |
| Body weight (g) |  |  |  |  |  |  | **<u>25.5 ± 1.15</u>** | **<u>16.6 ± 0.41</u>** | **<u><0.001</u>** |
| Food intake (kcal) | 0 | 0 |  | 0 | 0 |  | 0 | 0 |  |
| VO2 (mL/kg lean mass/h) | **<u>3604 ± 118</u>** | **<u>5151 ± 149</u>** | **<u><0.001</u>** | **<u>2794 ± 60</u>** | **<u>4099 ± 199</u>** | **<u><0.001</u>** | **<u>3199 ± 83</u>** | **<u>4675 ± 169</u>** | **<u><0.001</u>** |
| VCO2 (mL/kg lean mass/h) | **<u>2800 ± 98</u>** | **<u>4115 ± 106</u>** | **<u><0.001</u>** | **<u>2107 ± 57</u>** | **<u>3098 ± 161</u>** | **<u><0.001</u>** | **<u>2454 ± 71</u>** | **<u>3606 ± 129</u>** | **<u><0.001</u>** |
| RER (VCO2/VO2) | 0.77 ± 0.002 | 0.78 ± 0.006 | 0.105 | 0.75 ± 0.006 | 0.75 ± 0.008 | 0.982 | 0.76 ± 0.003 | 0.77 ± 0.006 | 0.492 |
| EE (kcal/kg lean mass/h) | **<u>17.2 ± 0.57</u>** | **<u>25.1 ± 0.69</u>** | **<u><0.001</u>** | **<u>13.3 ± 0.30</u>** | **<u>19.5 ± 0.95</u>** | **<u><0.001</u>** | **<u>15.2 ± 0.41</u>** | **<u>22.3 ± 0.80</u>** | **<u><0.001</u>** |
| Total activity (beam breaks) | **<u>36169 ± 1838</u>** | **<u>87131 ± 11695</u>** | **<u>0.001</u>** | **<u>9221 ± 1546</u>** | **<u>56436 ± 14366</u>** | **<u>0.008</u>** | **<u>45389 ± 3068</u>** | **<u>143567 ± 25225</u>** | **<u>0.002</u>** |
| Ambulatory activity (counts) | **<u>21867 ± 1863</u>** | **<u>62399 ± 9337</u>** | **<u>0.001</u>** | **<u>4886 ± 1252</u>** | **<u>42324 ± 11664</u>** | **<u>0.009</u>** | **<u>26753 ± 2956</u>** | **<u>104723 ± 20414</u>** | **<u>0.003</u>** |
|  | **<u>Re-feed (dark cycle)</u>** |  |  | **<u>Re-feed (light cycle)</u>** |  |  | **<u>Re-feed (24 hr)</u>** |  |  |
| Body weight (g) |  |  |  |  |  |  | **<u>27.9 ± 1.03</u>** | **<u>19.4 ± 0.55</u>** | **<u><0.001</u>** |
| Food intake (kcal) | 12.9 ± 0.62 | 13.4 ± 0.36 | 0.524 | 3.41 ± 0.37 | 4.01 ± 0.47 | 0.337 | 17.6 ± 1.05 | 17.3 ± 0.49 | 0.836 |
| VO2 (mL/kg lean mass/h) | **<u>4216 ± 145</u>** | **<u>4980 ± 184</u>** | **<u>0.006</u>** | **<u>3459 ± 154</u>** | **<u>4356 ± 126</u>** | **<u><0.001</u>** | **<u>3850 ± 147</u>** | **<u>4678 ± 132</u>** | **<u><0.001</u>** |
| VCO2 (mL/kg lean mass/h) | **<u>4200 ± 159</u>** | **<u>4966 ± 179</u>** | **<u>0.008</u>** | **<u>3502 ± 186</u>** | **<u>4431 ± 177</u>** | **<u>0.003</u>** | **<u>3873 ± 167</u>** | **<u>4706 ± 132</u>** | **<u>0.001</u>** |
| RER (VCO2/VO2) | 1.00 ± 0.005 | 1.00 ± 0.006 | 0.517 | 1.01 ± 0.014 | 1.01 ± 0.020 | 0.920 | 1.01 ± 0.009 | 1.00 ± 0.011 | 0.921 |
| EE (kcal/kg lean mass/h) | **<u>21.3 ± 0.75</u>** | **<u>25.1 ± 0.92</u>** | **<u>0.006</u>** | **<u>17.5 ± 0.81</u>** | **<u>22.1 ± 0.69</u>** | **<u><0.001</u>** | **<u>19.5 ± 0.76</u>** | **<u>23.6 ± 0.66</u>** | **<u><0.001</u>** |
| Total activity (beam breaks) | **<u>49750 ± 4033</u>** | **<u>65119 ± 5703</u>** | **<u>0.048</u>** | 9175 ± 1219 | 12966 ± 1506 | 0.073 | 37102 ± 4356 | 45853 ± 7582 | 0.349 |
| Ambulatory activity (counts) | 27781 ± 4810 | 38270 ± 3686 | 0.099 | 4326 ± 1186 | 6693 ± 1045 | 0.153 | 19302 ± 4436 | 26753 ± 5461 | 0.314 |

Eu: Euploid; VO2: rate of oxygen consumption; VCO2: rate of carbon dioxide production; RER: respiratory exchange ratio; EE: energy expenditure

**Table S3.** Tissue weight of high fat diet-fed male mice (33 weeks of age; fed diet for 16.5 weeks) at termination of the study.

|  | Eu | MAC21 | *P* |
| --- | --- | --- | --- |
| *N* | 8 | 8 |  |
| Body weight (g) | **46.62 ± 2.16** | **28.16 ±1.76** | **<0.001** |
| Gondal fat (gWAT, g) | **1.09 ±0.11** | **0.64 ±0.12** | **0.016** |
| Inguinal fat (iWAT, g) | **1.02 ±0.06** | **0.32 ±0.08** | **<0.001** |
| Liver (g) | **2.02 ±0.22** | **1.39 ±0.08** | **0.018** |
| Pancreas (g) | 0.25 ±0.03 | 0.25 ±0.01 | 0.933 |
| Kidney (g) | **0.18 ±0.01** | **0.15 ±0.01** | **0.026** |
| Spleen (g) | 0.09 ±0.01 | 0.09 ±0.01 | 0.851 |
| Diaphragm (g) | **0.11 ±0.01** | **0.09 ±0.01** | **0.010** |
| Heart (g) | 0.18 ±0.02 | 0.16 ±0.01 | 0.266 |
| BAT (g) | **0.32 ±0.04** | **0.12 ±0.02** | **0.001** |
| Brain (g) | **0.45 ±0.01** | **0.48 ±0.005** | **0.004** |
| Tibia (mm) | 18.25 ±0.21 | 18.04 ±0.15 | 0.421 |

Eu: Euploidy; gWAT: gondal white adipose tissue; iWAT: inguinal white adipose tissue; BAT: brown adipose tissue

**Table S4.**
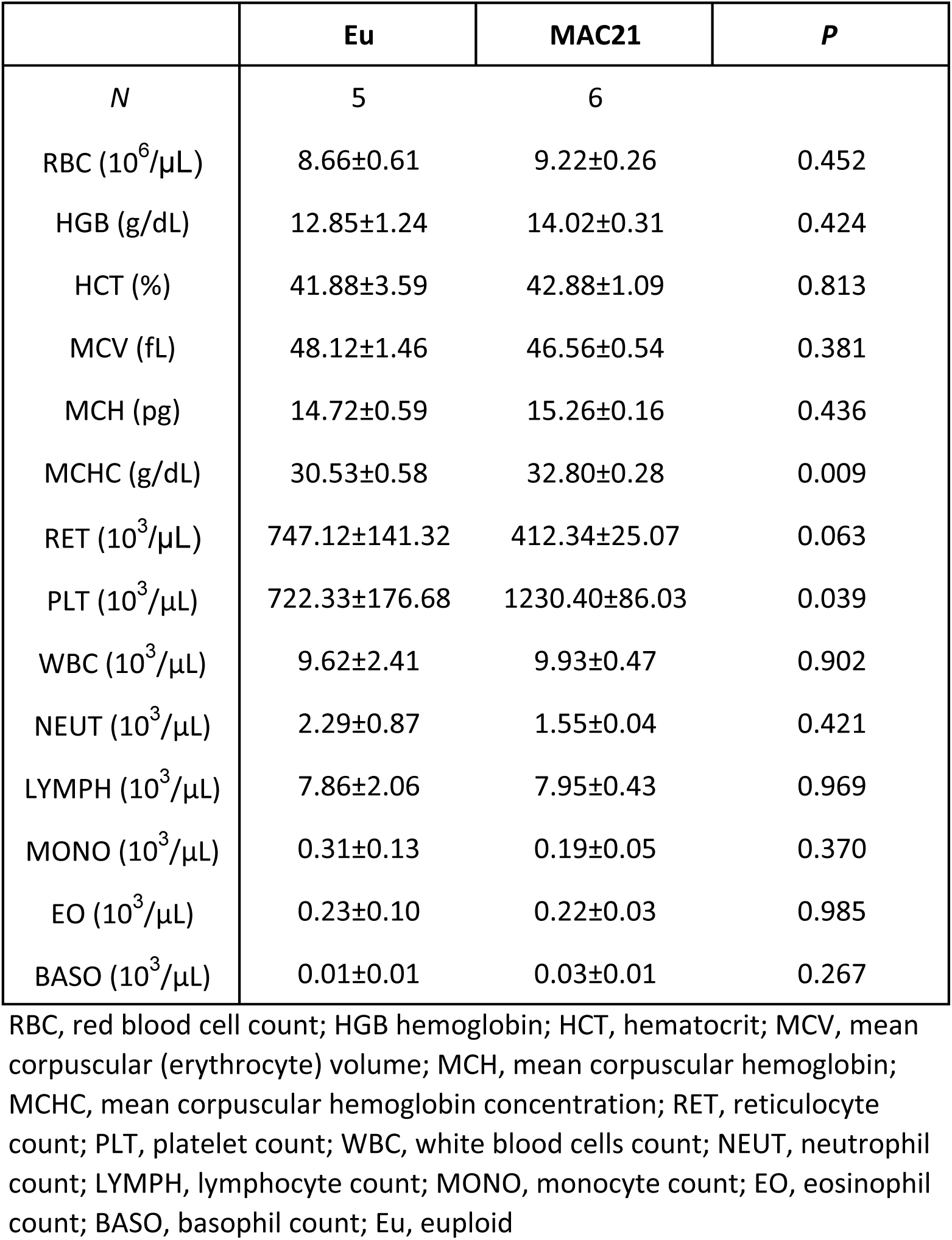
Complete blood count of Euploid and MAC21 mice fed a high-fat diet (50 weeks of age; on diet for 13 weeks)

|  | Eu | MAC21 | *P* |
| --- | --- | --- | --- |
| *N* | 5 | 6 |  |
| RBC (10<sup>6</sup>/μL) | 8.66±0.61 | 9.22±0.26 | 0.452 |
| HGB (g/dL) | 12.85±1.24 | 14.02±0.31 | 0.424 |
| HCT (%) | 41.88±3.59 | 42.88±1.09 | 0.813 |
| MCV (fL) | 48.12±1.46 | 46.56±0.54 | 0.381 |
| MCH (pg) | 14.72±0.59 | 15.26±0.16 | 0.436 |
| MCHC (g/dL) | 30.53±0.58 | 32.80±0.28 | 0.009 |
| RET (10<sup>3</sup>/μL) | 747.12±141.32 | 412.34±25.07 | 0.063 |
| PLT (10<sup>3</sup>/μL) | 722.33±176.68 | 1230.40±86.03 | 0.039 |
| WBC (10<sup>3</sup>/μL) | 9.62±2.41 | 9.93±0.47 | 0.902 |
| NEUT (10<sup>3</sup>/μL) | 2.29±0.87 | 1.55±0.04 | 0.421 |
| LYMPH (10<sup>3</sup>/μL) | 7.86±2.06 | 7.95±0.43 | 0.969 |
| MONO (10<sup>3</sup>/μL) | 0.31±0.13 | 0.19±0.05 | 0.370 |
| EO (10<sup>3</sup>/μL) | 0.23±0.10 | 0.22±0.03 | 0.985 |
| BASO (10<sup>3</sup>/μL) | 0.01±0.01 | 0.03±0.01 | 0.267 |

RBC, red blood cell count; HGB hemoglobin; HCT, hematocrit; MCV, mean corpuscular (erythrocyte) volume; MCH, mean corpuscular hemoglobin; MCHC, mean corpuscular hemoglobin concentration; RET, reticulocyte count; PLT, platelet count; WBC, white blood cells count; NEUT, neutrophil count; LYMPH, lymphocyte count; MONO, monocyte count; EO, eosinophil count; BASO, basophil count; Eu, euploid

**Table S5.** Indirect calorimetry analysis of male (25 weeks of age) Euploid and MAC21 littermate mice fed a high-fat diet (8.5 weeks on diet)

|  | Eu | MAC21 | P | Eu | MAC21 | P | Eu | MAC21 | P |
| --- | --- | --- | --- | --- | --- | --- | --- | --- | --- |
| **<u>High-fat diet (Male)</u>** | **<u>Ad-libitum (dark cycle)</u>** |  |  | **<u>Ad-libitum (light cycle)</u>** |  |  | **<u>Ad-libitum (24 hr)</u>** |  |  |
| N | 8 | 9 |  | 8 | 9 |  | 8 | 9 |  |
| Body weight (g) |  |  |  |  |  |  | **<u>43.0 ± 1.99</u>** | **<u>25.8 ± 1.06</u>** | **<u><0.001</u>** |
| Food intake (kcal) | 11.7 ± 1.02 | 12.6 ± 0.78 | 0.502 | 3.6 ± 0.61 | 3.6 ± 0.34 | 0.987 | 15.3 ± 1.20 | 16.2 ± 0.83 | 0.556 |
| VO2 (mL/kg lean mass/h) | **<u>4871 ± 107</u>** | **<u>6378 ± 316</u>** | **<u><0.001</u>** | **<u>4046 ± 74</u>** | **<u>5222 ± 250</u>** | **<u><0.001</u>** | **<u>4459 ± 84</u>** | **<u>5800 ± 281</u>** | **<u><0.001</u>** |
| VCO2 (mL/kg lean mass/h) | **<u>3956 ± 88</u>** | **<u>5316 ± 255</u>** | **<u><0.001</u>** | **<u>3270 ± 65</u>** | **<u>4274 ± 216</u>** | **<u><0.001</u>** | **<u>3613 ± 68</u>** | **<u>4795 ± 232</u>** | **<u><0.001</u>** |
| RER (VCO2/VO2) | **<u>0.81 ± 0.004</u>** | **<u>0.83 ± 0.003</u>** | **<u>0.001</u>** | 0.81 ± 0.004 | 0.82 ± 0.006 | 0.244 | **<u>0.81 ± 0.003</u>** | **<u>0.83 ± 0.004</u>** | **<u>0.011</u>** |
| EE (kcal/kg lean mass/h) | **<u>23.5 ± 0.51</u>** | **<u>30.9 ± 1.52</u>** | **<u><0.001</u>** | **<u>19.5 ± 0.36</u>** | **<u>25.2 ± 1.22</u>** | **<u><0.001</u>** | **<u>21.5 ± 0.40</u>** | **<u>28.0 ± 1.36</u>** | **<u><0.001</u>** |
| Total activity (beam breaks) | **<u>22378 ± 1644</u>** | **<u>41249 ± 4280</u>** | **<u>0.001</u>** | **<u>8153 ± 1249</u>** | **<u>14099 ± 1570</u>** | **<u>0.011</u>** | **<u>30531 ± 2553</u>** | **<u>16508 ± 5503</u>** | **<u>0.001</u>** |
| Ambulatory activity (counts) | **<u>11114 ± 1669</u>** | **<u>20126 ± 2350</u>** | **<u>0.008</u>** | 3571 ± 1295 | 5798 ± 730 | 0.144 | **<u>14685 ± 2859</u>** | **<u>25924 ± 2898</u>** | **<u>0.015</u>** |
|  | **<u>Fasting (dark cycle)</u>** |  |  | **<u>Fasting (light cycle)</u>** |  |  | **<u>Fasting (24 hr)</u>** |  |  |
| Body weight (g) |  |  |  |  |  |  | **<u>40.3 ± 1.89</u>** | **<u>23.1 ± 1.06</u>** | **<u><0.001</u>** |
| Food intake (kcal) | 0 | 0 |  | 0 | 0 |  | 0 | 0 |  |
| VO2 (mL/kg lean mass/h) | **<u>4389 ± 138</u>** | **<u>5063 ± 238</u>** | **<u>0.028</u>** | 3475 ± 84 | 3832 ± 176 | 0.089 | **<u>3932 ± 108</u>** | **<u>4447 ± 201</u>** | **<u>0.040</u>** |
| VCO2 (mL/kg lean mass/h) | **<u>3317 ± 101</u>** | **<u>3887 ± 172</u>** | **<u>0.013</u>** | 2650 ± 64 | 2953 ± 131 | 0.057 | **<u>2984 ± 80</u>** | **<u>3419 ± 146</u>** | **<u>0.020</u>** |
| RER (VCO2/VO2) | 0.76 ± 0.002 | 0.77 ± 0.007 | 0.096 | 0.76 ± 0.003 | 0.77 ± 0.004 | 0.149 | 0.76 ± 0.002 | 0.77 ± 0.005 | 0.109 |
| EE (kcal/kg lean mass/h) | **<u>20.8 ± 0.65</u>** | **<u>24.1 ± 1.12</u>** | **<u>0.024</u>** | 16.5 ± 0.40 | 18.3 ± 0.83 | 0.081 | **<u>18.7 ± 0.51</u>** | **<u>21.2 ± 0.94</u>** | **<u>0.035</u>** |
| Total activity (beam breaks) | **<u>25504 ± 3251</u>** | **<u>47893 ± 5189</u>** | **<u>0.003</u>** | **<u>6259 ± 815</u>** | **<u>14298 ± 1499</u>** | **<u><0.001</u>** | **<u>31763 ± 3793</u>** | **<u>62191 ± 6644</u>** | **<u>0.001</u>** |
| Ambulatory activity (counts) | **<u>13987 ± 1797</u>** | **<u>28435 ± 3571</u>** | **<u>0.003</u>** | **<u>2381 ± 392</u>** | **<u>7565 ± 1177</u>** | **<u><0.001</u>** | **<u>16368 ± 1981</u>** | **<u>36000 ± 4676</u>** | **<u>0.002</u>** |
|  | **<u>Re-feed (dark cycle)</u>** |  |  | **<u>Re-feed (light cycle)</u>** |  |  | **<u>Re-feed (24 hr)</u>** |  |  |
| Body weight (g) |  |  |  |  |  |  | **<u>41.4 ± 1.93</u>** | **<u>25.5 ± 0.90</u>** | **<u><0.001</u>** |
| Food intake (kcal) | 12.9 ± 0.62 | 13.4 ± 0.38 | 0.524 | 2.90 ± 0.62 | 3.6 ± 0.34 | 0.353 | **<u>13.0 ± 0.87</u>** | **<u>17.7 ± 0.82</u>** | **<u>0.002</u>** |
| VO2 (mL/kg lean mass/h) | **<u>4907 ± 133</u>** | **<u>5707 ± 225</u>** | **<u>0.008</u>** | **<u>3981 ± 77</u>** | **<u>4829 ± 187</u>** | **<u><0.001</u>** | **<u>4459 ± 100</u>** | **<u>5282 ± 203</u>** | **<u>0.003</u>** |
| VCO2 (mL/kg lean mass/h) | **<u>3916 ± 104</u>** | **<u>4812 ± 164</u>** | **<u><0.001</u>** | **<u>3233 ± 65</u>** | **<u>4130 ± 162</u>** | **<u><0.001</u>** | **<u>3586 ± 79</u>** | **<u>4482 ± 159</u>** | **<u><0.001</u>** |
| RER (VCO2/VO2) | **<u>0.80 ± 0.005</u>** | **<u>0.84 ± 0.008</u>** | **<u><0.001</u>** | **<u>0.81 ± 0.004</u>** | **<u>0.86 ± 0.011</u>** | **<u>0.002</u>** | **<u>0.80 ± 0.004</u>** | **<u>0.85 ± 0.009</u>** | **<u><0.001</u>** |
| EE (kcal/kg lean mass/h) | **<u>23.5 ± 0.63</u>** | **<u>27.7 ± 1.06</u>** | **<u>0.005</u>** | **<u>19.2 ± 0.37</u>** | **<u>23.5 ± 0.90</u>** | **<u><0.001</u>** | **<u>21.4 ± 0.48</u>** | **<u>25.7 ± 0.97</u>** | **<u>0.001</u>** |
| Total activity (beam breaks) | **<u>49750 ± 4033</u>** | **<u>65119 ± 6049</u>** | **<u>0.048</u>** | **<u>5118 ± 712</u>** | **<u>9820 ± 1222</u>** | **<u>0.005</u>** | **<u>27579 ± 2805</u>** | **<u>46524 ± 5236</u>** | **<u>0.007</u>** |
| Ambulatory activity (counts) | 27781 ± 4810 | 38270 ± 3910 | 0.100 | **<u>1618 ± 186</u>** | **<u>3424 ± 434</u>** | **<u>0.002</u>** | **<u>11970 ± 847</u>** | **<u>22377 ± 2708</u>** | **<u>0.003</u>** |

Eu: Euploid; VO2: rate of oxygen consumption; VCO2: rate of carbon dioxide production; RER: respiratory exchange ratio; EE: energy expenditure

**Table S6.**
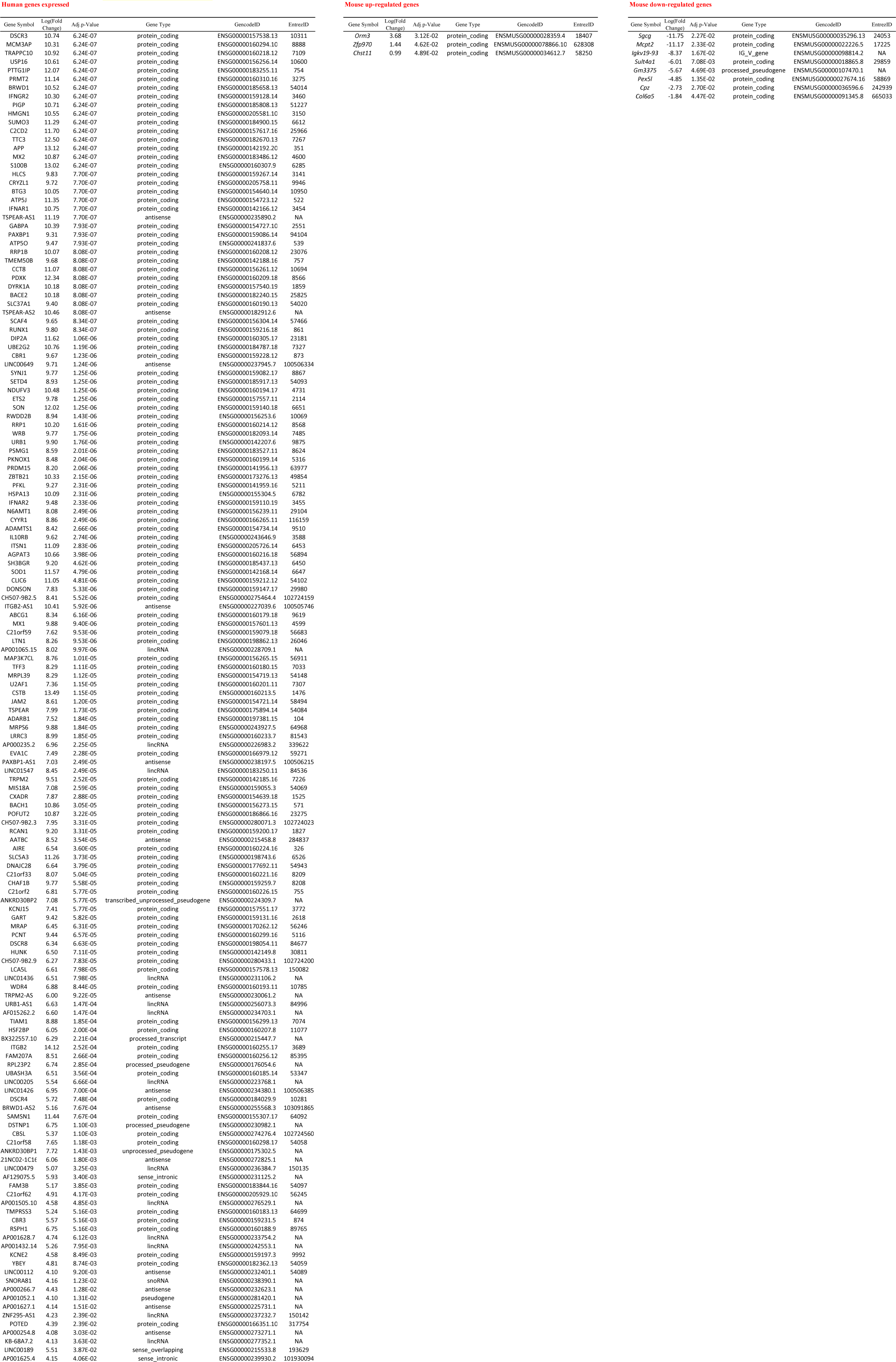
Gonadal White Adipose Tissue (HFD, 22°C)

**Table S7.**
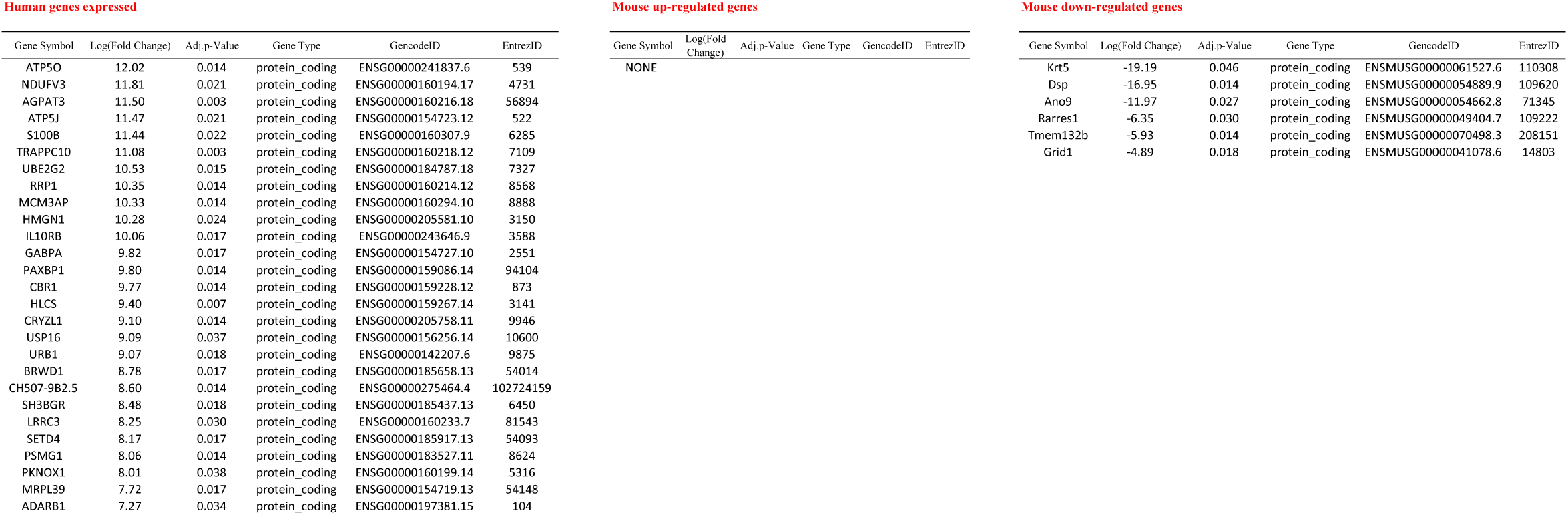
Inguinal White Adipose Tissue (HFD, 22°C)

**Table S8.**
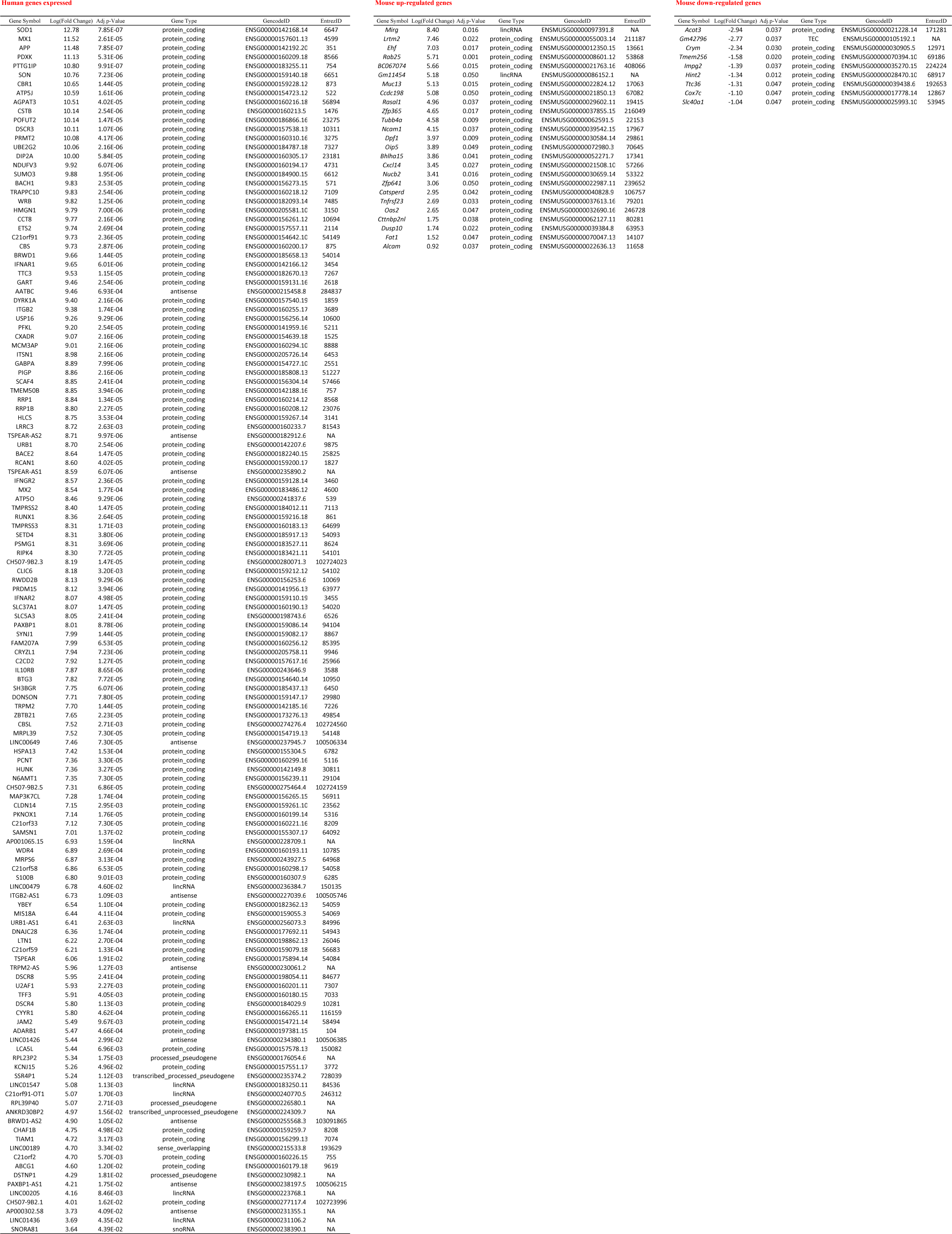
Liver (HFD, 22°C)

**Table S9.**
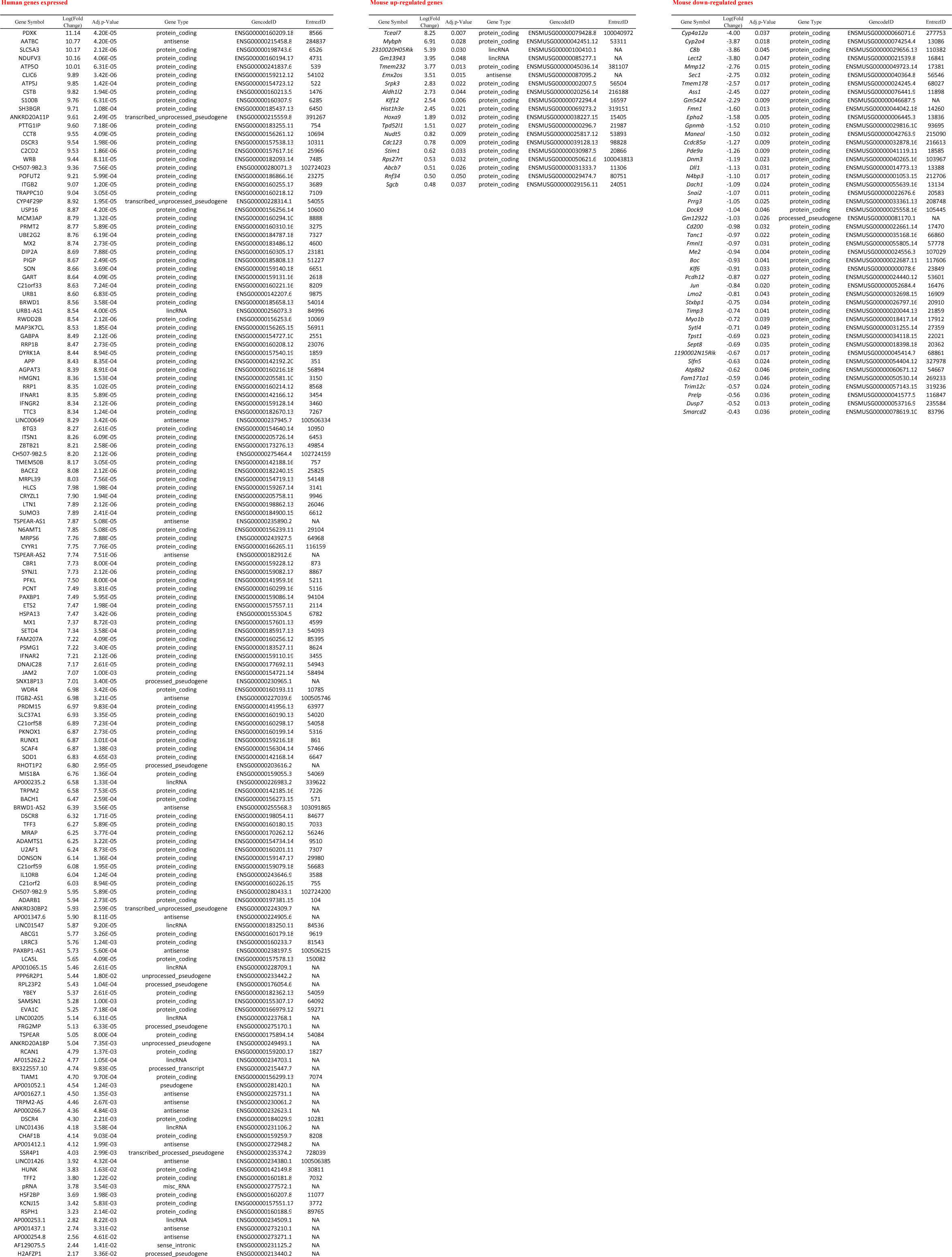
Brown Adipose Tissue (HFD, 22°C)

**Table S10.**
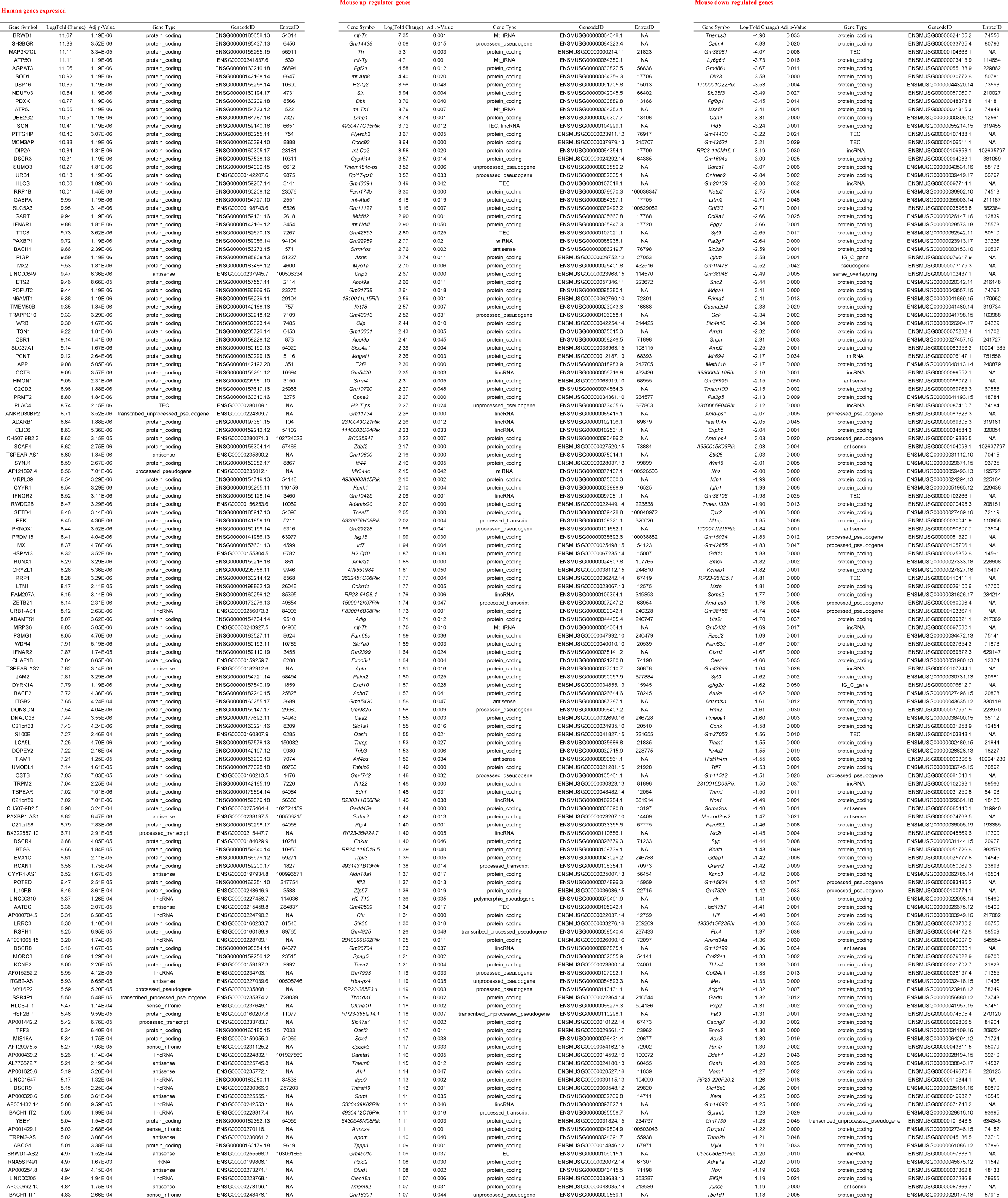
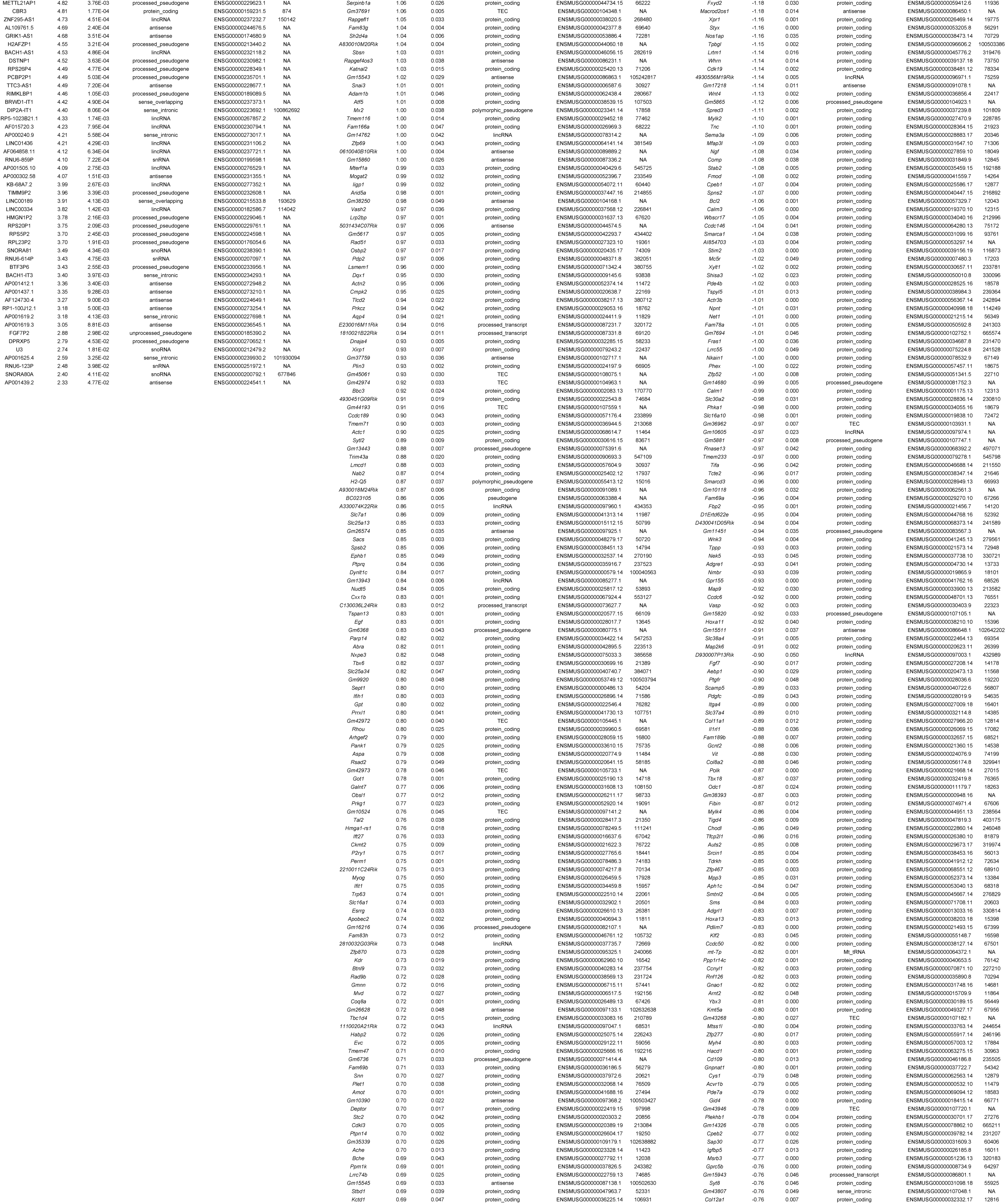
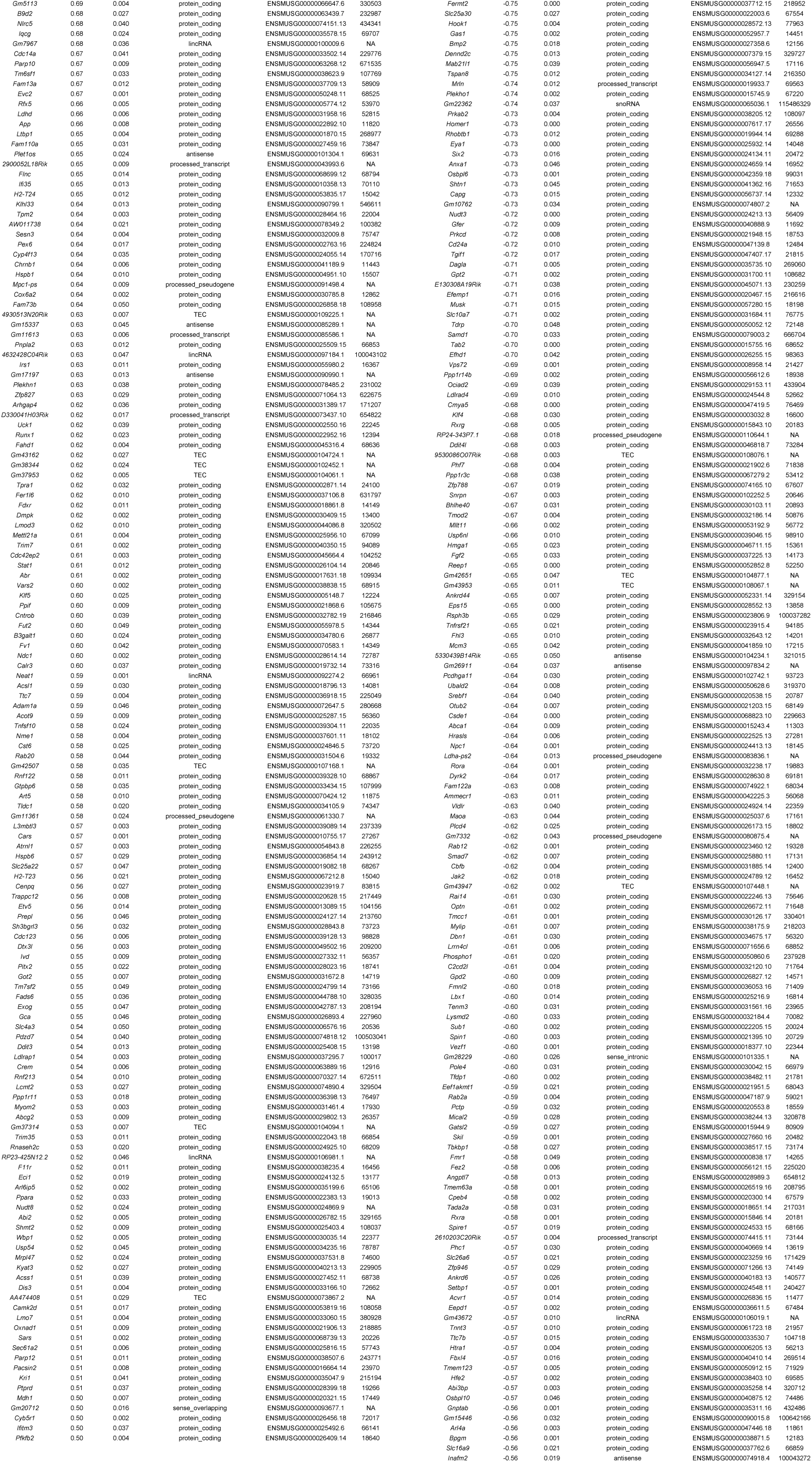
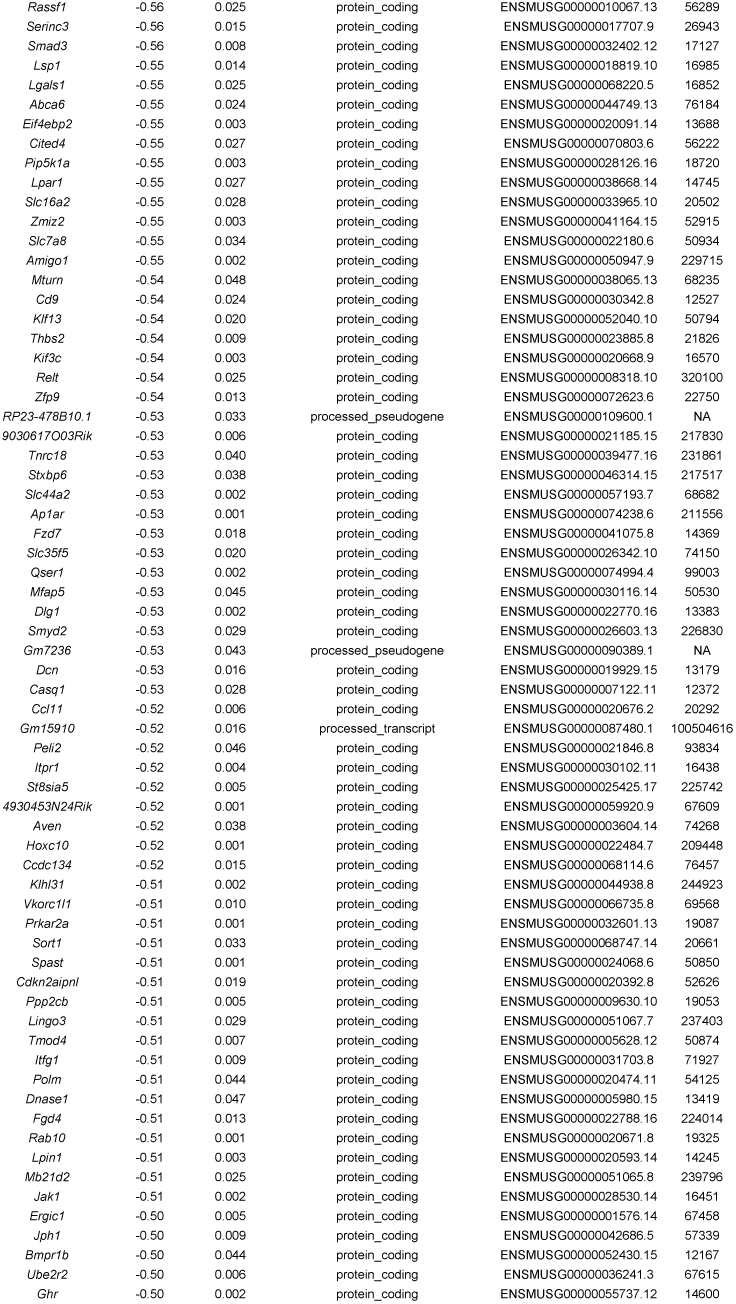
Skeletal muscle (HFD, 22°C)

**Table S11.** Indirect calorimetry analysis in thermoneutral conditions (31°C) after two-week acclimatization period of male (55 weeks of age) Euploid and MAC21 mice fed a high-fat diet (18 weeks on diet).

|  | Eu | MAC21 | P | Eu | MAC21 | P | Eu | MAC21 | P |
| --- | --- | --- | --- | --- | --- | --- | --- | --- | --- |
| **<u>High-fat diet (Male)</u>** | **<u>Ad-libitum (dark cycle)</u>** |  |  | **<u>Ad-libitum (light cycle)</u>** |  |  | **<u>Ad-libitum (24 hr)</u>** |  |  |
| N | 5 | 4 |  | 5 | 4 |  | 5 | 4 |  |
| Body weight (g) |  |  |  |  |  |  | **58.1 ± 6.18** | **36.0 ± 7.39** | **0.0020** |
| Food intake (kcal) |  |  |  |  |  |  | 7.67 ± 3.42 | 8.27 ± 3.87 | 0.814 |
| VO2 (mL/kg lean mass/h) | **3185 ± 120** | **4182 ± 103** | **<0.001** | **2480 ± 103** | **3134 ± 144** | **0.007** | **2833 ± 103** | **3658 ± 112** | **<0.001** |
| VCO2 (mL/kg lean mass/h) | **2230 ± 91** | **3017 ± 150** | **0.002** | **1766 ± 70** | **2360 ± 221** | **0.025** | **1998 ± 76** | **2688 ± 183** | **0.007** |
| RER (VCO2/VO2) | 0.70 ± 0.006 | 0.72 ± 0.024 | 0.342 | 0.71 ± 0.007 | 0.75 ± 0.033 | 0.245 | 0.70 ± 0.006 | 0.73 ± 0.028 | 0.282 |
| EE (kcal/kg lean mass/h) | **14.9 ± 0.57** | **19.7 ± 0.55** | **<0.001** | **11.6 ± 0.48** | **14.9 ± 0.82** | **0.009** | **13.3 ± 0.48** | **17.3 ± 0.64** | **0.001** |

Eu: Euploid; VO2: rate of oxygen consumption; VCO2: rate of carbon dioxide production; RER: respiratory exchange ratio; EE: energy expenditure

**Table S12.**
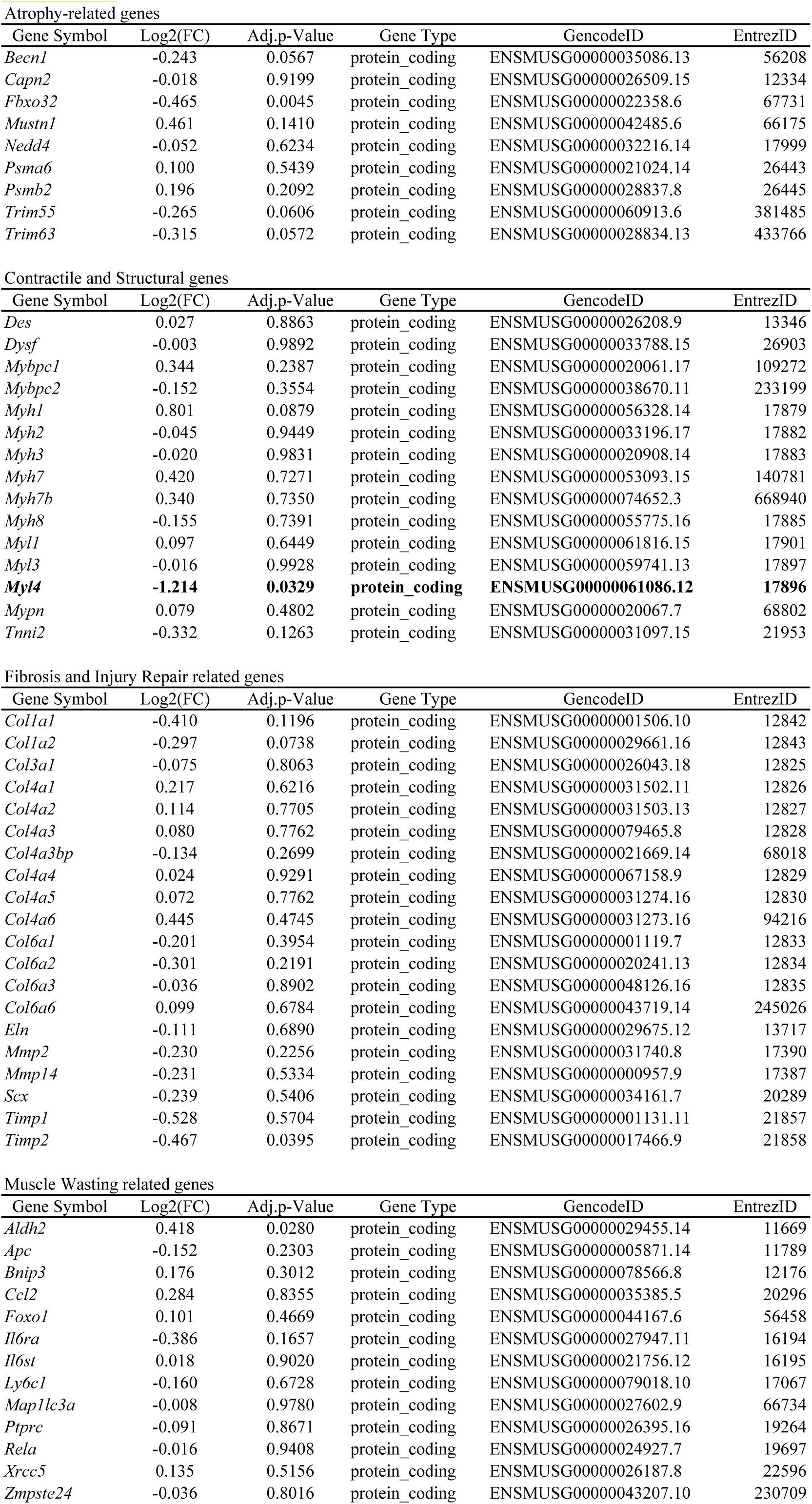
With the exeption of Myl4, none of the genes are signficantly upregulated above log2(FC) of 0.5

**Table S13.** Tissue weights of male MAC21 and Euploid littermates (55 weeks of age) housed at thermoneutrality (31°C) and fed an HFD (18 weeks on diet).

|  | Eu | MAC21 | P |
| --- | --- | --- | --- |
| N | 5 | 4 |  |
| Body weight (g) | 60.3 ± 2.85 | 36.9 ± 3.71 | 0.001 |
| Gondal fat (gWAT, g) | 1.62 ± 0.22 | 1.07 ± 0.05 | 0.065 |
| Inguinal fat (iWAT, g) | 1.60 ± 0.11 | 0.58 ± 0.11 | <0.001 |
| Liver (g) | 3.03 ± 0.33 | 1.66 ± 0.19 | 0.013 |
| BAT (g) | 0.25 ± 0.02 | 0.10 ± 0.02 | <0.001 |
| Pancreas (g) | 0.44 ± 0.02 | 0.34 ± 0.03 | 0.015 |

Eu: Euploid; gWAT: gondal white adipose tissue; iWAT: inguinal white adipose tissue; BAT: brown adipose tissue

**Table S14.**
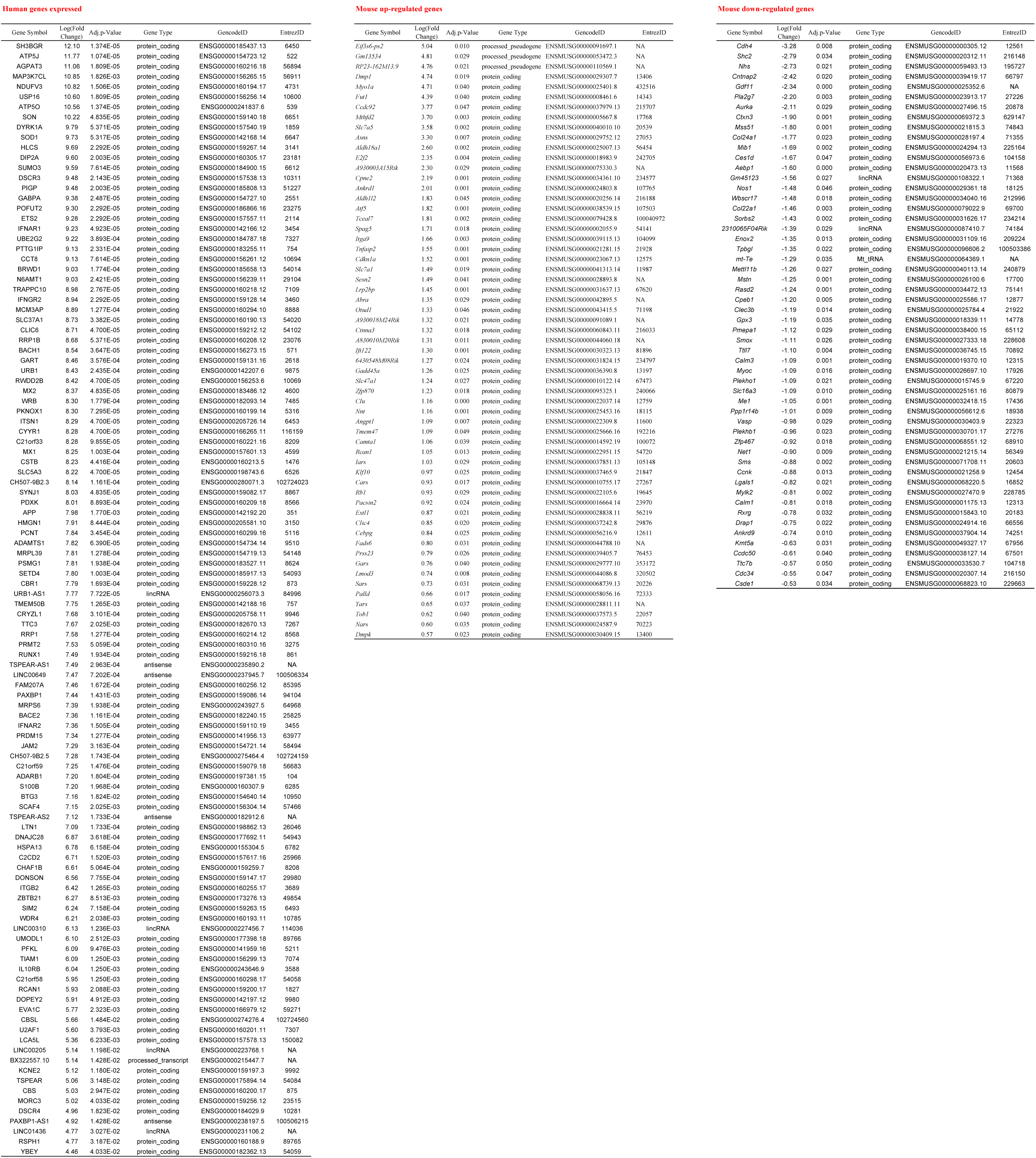
Skeletal muscle (HFD, 31°C)

**Table S15.**
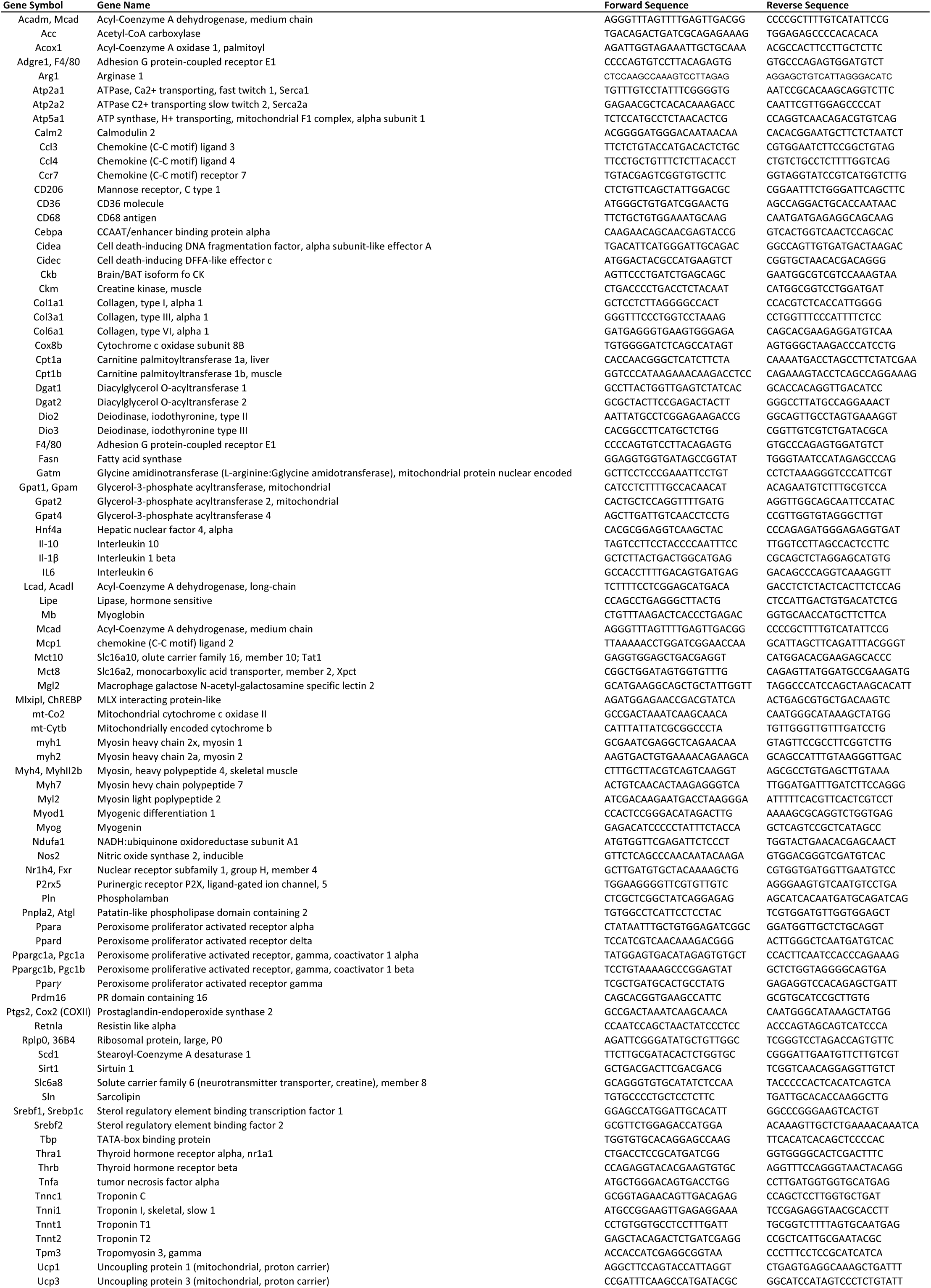
List of primers used for qPCR analysis of gene expression.

